# Plant phenotypic differentiation outweighs genetic variation in shaping the lettuce leaf microbiota

**DOI:** 10.1101/2024.10.16.618533

**Authors:** Arianna Capparotto, Guillaume Chesneau, Alessandra Tondello, Piergiorgio Stevanato, Tiziano Bonato, Andrea Squartini, Stéphane Hacquard, Marco Giovannetti

## Abstract

Lettuce is a widely consumed raw vegetable, making it crucial to understand and predict its leaf-associated microbial communities for the benefit of both plant and human health. While environmental factors are known to strongly influence plant leaf microbiomes, the role of plant-specific determinants in shaping microbial diversity remains unclear. In this study, we investigated the impact of three key plant factors -genetic distance, morphology and leaf micro- and macronutrient content- on the composition and diversity of lettuce leaf bacterial communities.

Using 131 fully-sequenced *Lactuca sativa* genotypes, we analyzed their leaf-associated bacterial communities via 16S rRNA amplicon sequencing. Our findings revealed that morphological classification, as defined by breeders, exerts a greater influence on bacterial community diversity than genetic distance or variations in leaf nutrient levels. Together with shoot traits they explained 13.9% of the observed bacterial diversity. Further analysis of 10 specific leaf morphological traits showed that heart formation, head height, and venation types significantly shaped bacterial richness and evenness, mainly acting on non-hub members.

The strong association between leaf morphology and bacterial community structure suggests that phenotypic traits play a disproportionately large, yet understudied, role in leaf microbiota establishment offering new potential for manipulation by breeders.

## INTRODUCTION

Lettuce (*Lactuca sativa*), a leafy vegetable from the Asteraceae family, is one of the most widely consumed green vegetables, holding a significant position in global agriculture [1]. The widespread consumption of lettuce can be largely attributed to its favorable nutritional profile: its leaves are low in calories and fats while being rich in fiber, vitamins, and essential minerals such as iron [2]. In addition to this, emerging evidence highlights the beneficial aspects of the “edible microbiome” associated with lettuce leaves [3, 4]. For example, bacteria present in vegetables, including lettuce, have been shown to enhance the diversity of the human gut microbiome [5]. Additionally, lactic acid bacteria found on the surface of lettuce leaves can survive digestion and persist in the gut, positively influencing its microbial composition [5, 6]. Given this evidence, advancing our understanding of the factors that shape the bacterial community on lettuce leaves could foster beneficial microbial populations that are important for both human nutrition and plant health, ultimately reducing resource losses.

While environmental factors, such as air temperature, humidity, solar radiation, wind, geographic location and local neighborhood identity account for most of the variation in leaf bacterial communities [7–9], emerging evidence also points to the impact of intrinsic plant determinants, such as genotypes, leaf morphology and secretion of secondary metabolites, as secondary but important contributors to this variation [10–12]. In lettuce, the impact of genotype has been demonstrated for fungi [13] and for culturable bacteria-associated communities [14, 15]. Recently, genome-wide association (GWA) has been employed to associate host genetic variation with phyllosphere microbiota differentiation, identifying genes potentially associated with microbiome variation [16]. However, the extent to which plant genetic variability shapes the assembly of microbial communities remains poorly understood, partly due to the limited availability of comprehensive leaf microbiota studies on non-model plants. In addition, increasing evidence supports the role of plant morphological and physiological traits in driving microbial community composition. For example, small- scale studies on lettuce have shown that macroscopic [14], as well as microscopic features like stomatal density, epidermal cell patterning, and cuticle hydrophobicity [17], create preferential attachment sites for fungi or bacteria. Regarding leaf nutrient homeostasis, studies in spinach, rocket salad [18], rice [19] and Cassava [20] have shown significant correlations between leaf minerals and diversity indices. Still, none have demonstrated a causative effect of leaf nutritional status on bacterial community distribution. These morphological and physiological differences can lead to uneven bacterial distribution across different plant types, underscoring the need for more integrative studies to understand the interplay between plant genetics, morphology, leaf nutrient content and microbial community structure.

While lettuce has not traditionally been used as a model organism, recent advances have considerably expanded our understanding of its evolutionary history and the molecular mechanisms underlying key plant traits. Notably, Wei and collaborators (2021) fully sequenced 445 *Lactuca* genotypes from accessions preserved by the Centre for Genetic Resources in the Netherlands (CGN). This extensive sequencing effort generated a detailed variation map, combined with available phenotypic traits, providing a valuable resource for exploring phylogenetic relationships and advancing breeding programs.

In this work, we took advantage of this comprehensive collection to conduct a large-scale field experiment using 131 *Lactuca sativa* genotypes. We hypothesized that key and manipulable plant determinants such as -genetic distance, leaf micro- and macronutrient content, and leaf morphology- may influence the leaf-associated bacterial community establishment and differentiation. Taking into account the complex interdependencies existing between plant genotype, phenotype and physiology [10, 21, 22], we uncovered the importance of considering phenotype-microbiome association to better understand, and possibly predict, the phyllosphere microbiota.

## MATERIAL AND METHODS

### Experimental design

In this study, 131 fully sequenced genotypes of *Lactuca sativa* were selected from the Centre for Genetic Resources (CGN, Wageningen, Netherlands). Seeds were sown and germinated in a growth chamber for 24 hours and then grown under greenhouse conditions for nearly one month at Bronte Garden (Mira, Italy). Plants were then transferred to the greenhouse of “Azienda Agricola Gambaro” (Noale, Italy). For the experiment, nine replicates per genotype were transferred into three blocks labeled as “A,” “B,” and “C” in the soil, following a randomized design outlined in Table S1. Each block consisted of 20 lines (numbered 1 to 20) and 20 columns (named a to t), maintaining a plant distance of 20x20 cm. Three replicates per genotype were cultivated inside each block. Throughout the experiment’s duration, plants were irrigated using drip irrigation.

### Sample collection

Leaf sample collection took place on May 20^th^-21^st^, 2022. During this process, a leaf from the mid-upper part of each plant was selected. Prior to sampling, leaves underwent surface washing with sterile water. The sample collection was done in racks of 96 collection microtubes (QIAGEN Hilden, Germany), following the sequential order outlined in Table S2 and Table S3. Control soil samples (germination substrate, growth soil stage 0, growth soil stage 1 and endophytes) were collected at three different time points as detailed in supplementary materials and methods. Immediately after collection, all samples were transported to the laboratory on ice and stored at -20°C. Leaf disks were lyophilized for long-term storage.

### Leaf mineral content quantification

Leaf disks were used to quantify leaf micro- and macro-nutrient content using the Inductively Coupled Plasma-Optical Emission Spectroscopy (ICP-OES) technology at SESA (Este, Italy). Briefly, samples were weighted with an ultra-analytical balance (sensitivity = 0.01 mg) and then digested at 95°C for 2 hours in nitric acid (HNO3 67-69%) using the DigiPREP (SCP Sciences). Samples were then diluted in 20 ml of ultrapure water and filtered through 25 mm diameter, 0.45-micron pore-size, removable syringe-tip filter membranes (cellulose acetate). 50 μl of yttrium was added as an internal standard to all samples before initiating the ICP-OES detection. Minerals that were poorly detected were excluded from the analysis. For the predominantly detected ones, samples falling below the limit of quantification (<LOQ) were fit in the model, following the procedure reported in a published method [23]. The detailed above-mentioned data and the processing steps are reported in Table S4.

### DNA extraction and sequencing

The leaf disk DNA isolation was performed on a large scale using BioSprint® technology (QIAGEN, Hilden, Germany) in 96-well collection microtubes. Seven empty wells were included as controls to identify potential contaminants from the extraction process. Conversely, DNA extraction from soil and seed samples was carried out using the DNeasy® PowerSoil® kit (QIAGEN Hilden, Germany) following the manufacturer’s instructions.

To isolate endophytes, a pool of 131 leaves underwent surface sterilization following the procedure outlined in previous methods [24] with some modifications. DNA isolation was then carried out using the CTAB protocol. DNA from all samples was quantified using Qubit™ dsDNA Assay Kits from Thermo Fisher Scientific, and their concentrations are detailed in Table S5. Subsequently, the DNA samples underwent MiSeq Illumina 16S amplicon sequencing at IGA Technology Services (IGATech, Udine, Italy) following the procedure reported in supplementary material and methods.

### Raw reads processing

Raw sequencing reads were subjected to processing using Dada2 (v.1.10) in QIIME2 (QIIME2-2023.2). The resulting dataset underwent additional refinement in R (version 4.3.2) to eliminate chloroplasts, eukarya, archaea, mitochondria (Table S6) and possible contaminations that occurred during extraction using the Decontam package (version 1.24.0). The final dataset consists of 12,173 ASVs and 417 samples. R’s Vegan package (version 2.6- 4) was employed to rarefy the dataset in a size-dependent manner. Given the disparate sampling depths across compartments, each was rarefied to a distinct sampling depth (leaf: sample size = 1,000 reads, soil: sample size = 14,000 reads, seeds: sample size = 200 reads, endophytes: sample size = 4,500 reads). Using the Phyloseq package (version 1.44.0), the rarefied datasets were merged into a Phyloseq object, comprising 9,889 ASVs and 259 samples (9,125 leaf ASVs and 230 leaf samples).

### Identification of groups of closely related genotypes and varieties

The lettuce phylogenetic tree, as delineated by [25], served as the basis for extracting the coordinates of the *Lactuca sativa* accessions tree. These were subsequently utilized to construct a novel phylogenetic tree in ITOL. A distant member belonging to the *Lactuca saligna* species (TKI-349) was selected as the outgroup to root the tree. Six distinct groups of closely related genotypes were identified based on branch ages. Seven breeder-defined varieties were used as proxies for plant phenotypes. Two genotypes, TKI-071 and TKI-119, did not clearly belong to any genetic distance group and were thus omitted from the analysis. Additionally, TKI-11 was excluded due to a lack of tree-coordinate information, and TKI-92 was removed because there were insufficient phenotypic parameters to impute the missing values. This process resulted in the exclusion of the oilseed variety group. The remaining samples were distributed as described in Table S7.

### Community composition and contributions of plant determinants to **β** diversity

To investigate statistically significant differences at various taxonomic ranks (up to the genus level) between groups of closely related genotypes and varieties, the Kruskal-Wallis test with Dunn’s *post hoc* test (*p-adjusted* method = “fdr”) was applied. Adjusted p-values less than 0.05 were considered significant. The hierarchical tree of the community at each taxonomic level was constructed using GraPhlAn (Graphical Phylogenetic Analysis), version 1.1.4, [26], in Python3.

β-diversity across leaf samples was assessed by the Bray-Curtis dissimilarity distance matrix in R using the vegan package. The influence of each plant factor in determining this diversity was tested with the Adonis2 implementation of Permutational Multivariate Analysis of Variance (PERMANOVA). To disentangle the independent and combined proportions of variance explained by these significant determinants, we employed the varpart function from the vegan package. To ensure the effectiveness of varpart, missing values in the dataset’s phenotypic traits were imputed based on their correlations. Initially, a correlation matrix among the phenotypic traits was computed (Fig. S1). Subsequently, factors demonstrating the strongest significant correlations were chosen to impute NA values, after fitting a correlation line through their data points. Table S8 represents the dataset before and after imputation. Other details are provided in supplementary material and methods.

### Integrative analysis of leaf bacterial communities: functional predictions, diversity, networks, and origins

Functional predictive analysis (Functional Annotation of Prokaryotic Taxa, FAPROTAX, [27]) was conducted using the microeco package [28] in R (version 1.8.0). Statistical differences in bacterial types between leaf morphological traits were assessed using the Kruskal-Wallis test with Dunn’s *post hoc* test.

Phenotypic traits exhibiting differences in α diversity were analyzed using the log2foldchange function from the DESeq2 package in R (version 1.40.2). The igraph package (version 2.0.3) was used to visualize the bacterial network for phenotypic groups showing differences in α diversity. The fast expectation–maximization microbial source tracking algorithm (FEAST) package (version 0.1.0, [29]) was used to calculate the relative share of ASVs between the leaf (considered the sink) and four different sources: seeds, germination soil, growth soil before (April 20, 2021) and after (May 21, 2021) plant transplantation, and the endophytic bacterial community.

A more detailed description of the methods and the script used for all the analyses are provided as supplementary material and methods and supplementary script, respectively.

## RESULTS

### Genetic distance and variety influence leaf bacterial community composition and diversity

To assess the impact of genetic distance and variety on the composition and shifts of leaf-associated bacteria, we first extracted and used the phylogenetic tree coordinates for 131 *Lactuca sativa* genotypes [25] to reconstruct the tree represented in Fig. 1. A partial overlap between variety groups and genetic distance groups was observed, as reported in Fig. 1 and Table 1. For instance, 81.94% of Butterhead plants overlap with genetic group 6, while Cos plants correspond mainly with group 2 (88.24%) and Crisp plants with group 4 (84.21%). However, other genetic groups (1, 3, and 5) are composed of several different varieties. The discrepancy between genetic groups and varieties underlies the importance of uncoupling phenotypic and genetic variation when studying the phyllosphere microbiota assembly.

**Fig. 1.**
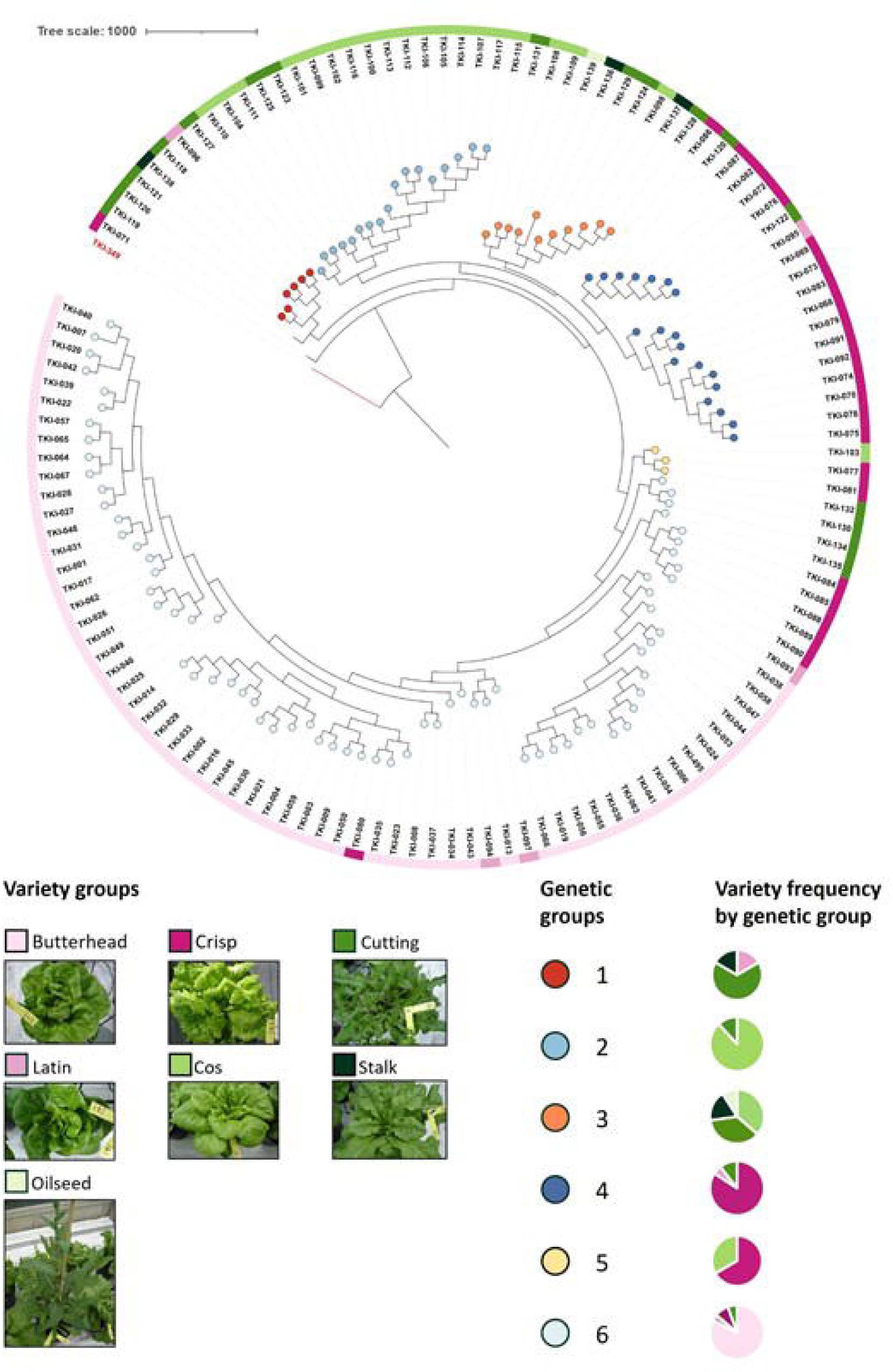
Neighbor-joining tree of 131 *Lactuca sativa* accessions. Tip colors indicate six groups of closely related genotypes, while colored squares represent different varieties. The phenotypic appearance of each variety group is depicted alongside. Bar charts show the proportion of each variety within each genetic group.

**Table 1.** Percentage of each variety type within groups of closely related genotypes.

|  | Butterhead | Cos | Crisp | Cutting | Latin | Stalk | Oilseed |
| --- | --- | --- | --- | --- | --- | --- | --- |
| 1 |  |  |  | 66.67% | 16.67% | 16.67% |  |
| 2 |  | 88.24% |  | 11.76% |  |  |  |
| 3 |  | 36.36% |  | 36.36% |  | 18.18% | 9.09% |
| 4 |  |  | 84.21% | 10.53% | 5.26% |  |  |
| 5 |  | 33.33% | 66.67% |  |  |  |  |
| 6 | 81.94% |  | 8.33% | 5.56% | 4.17% |  |  |

Overall, the leaf bacterial communities across all *Lactuca sativa* genotypes (Fig. 2A) were dominated by Proteobacteria (71.3%), followed by Bacteroidota (8.4%), Firmicutes (7%), Actinobacteriota (5.1%), Planctomycetota (1.1%) and Verrucomicrobiota (1.1%). The FEAST algorithm was implemented to identify the source most likely driving the establishment of the leaf microbial community. This analysis revealed that, among the potential sources and by taking into account field block position, the seed microbiome contributed with the largest relative share (average contribution=15,36%), followed by the leaf endophytic bacterial ASVs (average contribution=5,1%), the soil at harvesting stage (average contribution=1,2%), the soil at the transplanting stage (average contribution=0.63%), and finally, the germination substrate (average contribution=0.39%, Fig. S2A-C, Table S9).

**Fig. 2.**
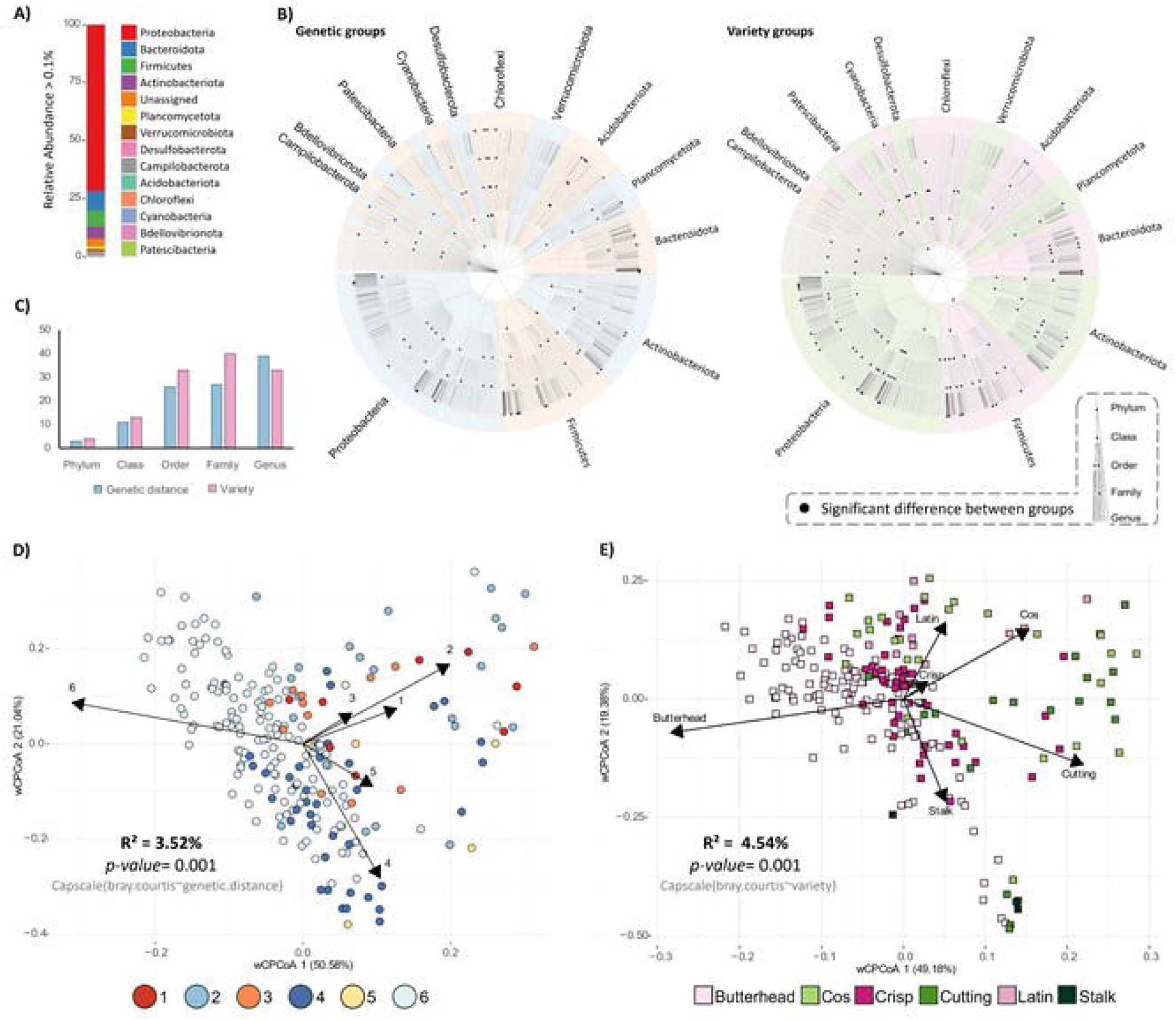
Leaf bacterial community composition and diversity as influenced by genetic distance and variety. A) Stacked bar plot showing the most abundant community members (relative abundance >0.1%), from top to bottom. B) Significant differences in taxa between groups of closely related genotypes and varieties. Most abundant members of the leaf-associated bacterial community (relative abundance >0.1%) are shown in the text. Nodes represent taxonomic ranks from Phylum (inner ring) to Genus (outer ring). Black dots indicate statistically significant differences of a node between groups of closely related genotypes (left side), and varieties (right side), assessed using Kruskal-Wallis and Dunn *post hoc* tests. Colors distinguish taxonomic groups between genetic groups (blue and orange, right side), and varieties (pink and green, left side). C) Disparities in significant nodes between varieties and groups of closely related genotypes across all taxonomic levels. D-E) Constrained Analysis of Principal Coordinates (CAP) performed on the Bray-Curtis distance matrix showing the contribution to leaf-associated bacterial community β-diversity of groups of closely related genotypes, and varieties, respectively. Each circle/square represents a single sample.

The effects of genetic distance and variety on bacterial community composition (Fig. 2B-2C) involve variation in different community members (Table S10). At higher taxonomic ranks - from Phylum to Family- variety determines more bacterial shifts. However, at the genus level, genetic distance has a greater impact, affecting more genera -39- than those influenced by variety -33- (Fig. 2C- Table S10). The influence of plant variety on bacterial community composition is further evidenced by its impact on bacterial β-diversity, which slightly surpasses the contribution of genetic distance. Specifically, plant genetic distance accounted for 3.52% (PERMANOVA, p-value= 0.001) of the total variation in leaf-associated bacterial communities, while variety accounted for 4.54% (PERMANOVA, p-value= 0.001, Fig. 2D- E). Moreover, across varieties butterhead plants show significantly higher Shannon and observed index compared to Cos and Cutting plants (p-value <0.05), while no difference in α- diversity is observed between genetic groups (Fig. S3). Altogether, these observations suggest that specific plant traits related to plant genotype and/or variety may differentially regulate leaf microbiota.

### Micro- and macronutrient concentrations partially affect leaf-associated bacterial **β**- diversity

To further investigate the plant leaf traits that may explain the differences in bacterial β- diversity observed across genetic and variety groups, we focused on a subset (Ca, Fe, K, Mg, Mn, Na, P, Zn) of leaf macro- and micro-elements. The strong correlations among most leaf elements (Fig. S4A), led us to perform a PCA ordination. Through this approach, we identified five PCs that collectively capture the combined contributions of these nutrient variables (Fig. 3A). Only PCs 2, 4, and 5, which capture opposing fluctuations between micro- and macronutrient concentration (Fig. S4B), had statistically significant small effects (PERMANOVA, p-value <0.05) on shaping bacterial β-diversity (Table S11).

**Fig. 3.**
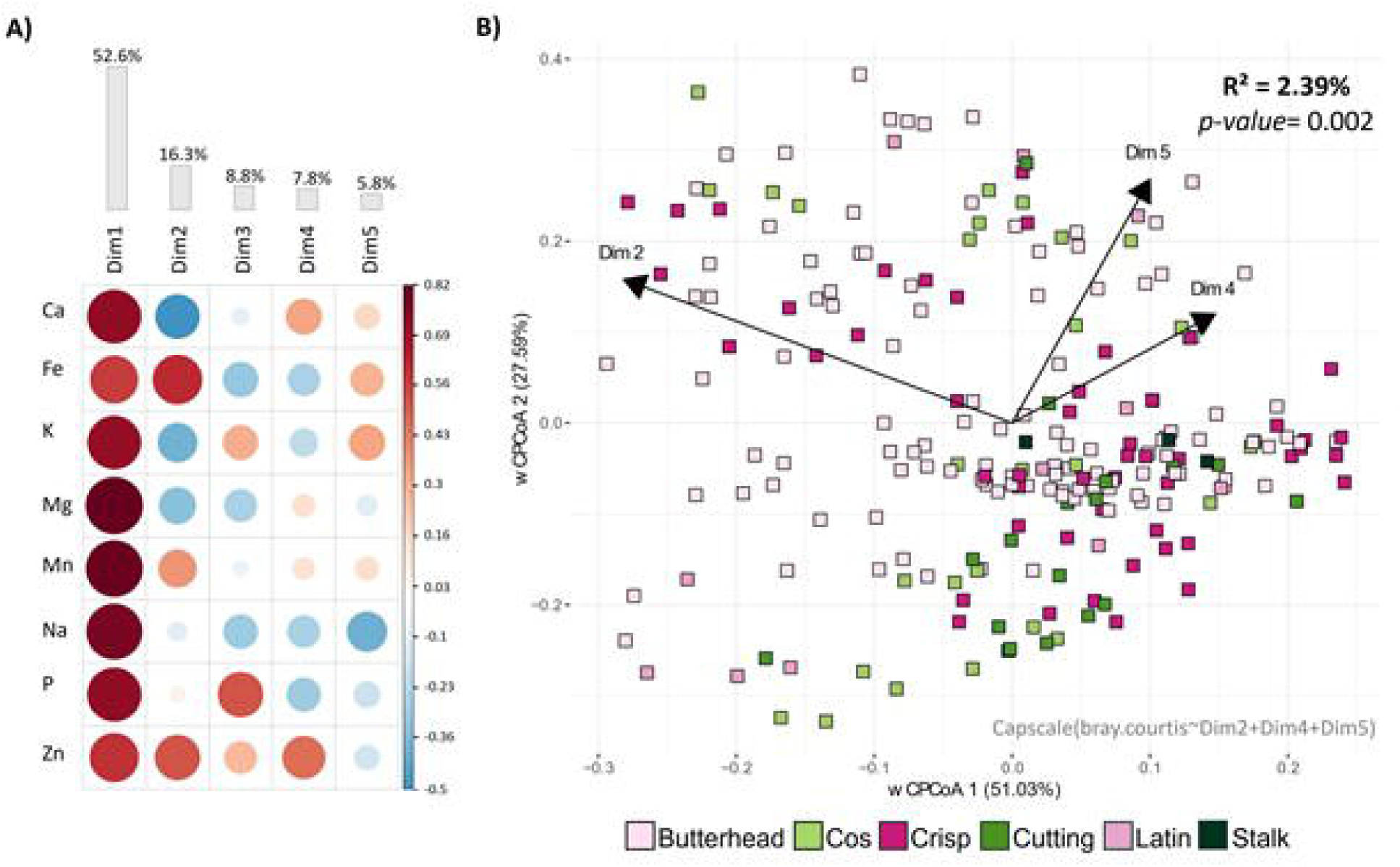
Correlogram and multivariate analysis linking mineral elements density with bacterial community distances. A) Correlation matrix depicting the relationships between micro- and macronutrients and their PCs. Red indicates positive correlations, while blue indicates negative correlations. Larger dots indicate stronger correlations. B) Constrained Analysis of Principal Coordinates (CAP) performed on the Bray-Curtis distance matrix illustrating the influence of significant mineral PCs (arrows) on bacterial diversity. Each square represents a single sample, color- coded according to the variety group.

We constrained the bacterial Bray-Curtis dissimilarity distance matrix to reflect only the total variation explained by the significant dimensions 2, 4, and 5. The PERMANOVA statistical analysis revealed that leaf micro- and macro-nutrient content accounted for 2.38% (PERMANOVA, p-value=0.002) of the bacterial β-diversity (Fig. 3B). In addition, they also demonstrated a minor impact on bacterial α-diversity with Dimension 2 being slightly significantly positively correlated with bacterial Shannon and observed indices (Fig. S5A-B).

Overall, these observations suggest that leaf nutritional status exerts a limited influence on the plant leaf-associated microbiota. Moreover, the results underscore the importance of considering the combined effects of micro- and macro-nutrients rather than their individual contributions (Fig. S4B-D).

### Leaf morphological traits disentangle the variety contribution to **β**-diversity

Given the significant impact of variety on bacterial variation among the three factors investigated, we hypothesized that increasing the resolution of this trait, particularly by including plant phenotypic parameters, could provide a more detailed understanding of factors shaping the leaf-associated bacterial community. We thus included 17 phenotypic parameters in the analysis (Table S8). These parameters, already characterized for each genotype by the collection curators and available on the CNG lettuce collection database, encompass a range of traits: leaf characteristics (e.g., shape, venation), pigmentation-related attributes (e.g., leaf color, anthocyanin content), as well as traits related to juvenile stages (e.g., seedling cotyledon shape) and adult plants (e.g., heart formation, head height).

The PERMANOVA statistical analysis revealed that out of the 17 phenotypic parameters investigated, 10 were significantly (PERMANOVA, p-value <0.05) associated with leaf bacterial β-diversity (Table S12). Among these, the most impactful parameter was heart formation (R^2^=0.012, p-value=0.001), followed by head height (R^2^=0.01, p-value=0.001), head shape (R^2^=0.012, p-value=0.001) and leaf venation (R^2^=0.01, p-value=0.001).

We found that the above-mentioned leaf morphological traits explained 6.52% of the variation in community composition (PERMANOVA, p-value=0.001, Fig 4) and also allowed a clear clustering among different lettuce varieties. For example, Butterhead lettuce, known for its tight heart formation capacity, is strongly associated with high values in traits such as head shape, heart formation, and head leaf overlap.

**Fig. 4.**
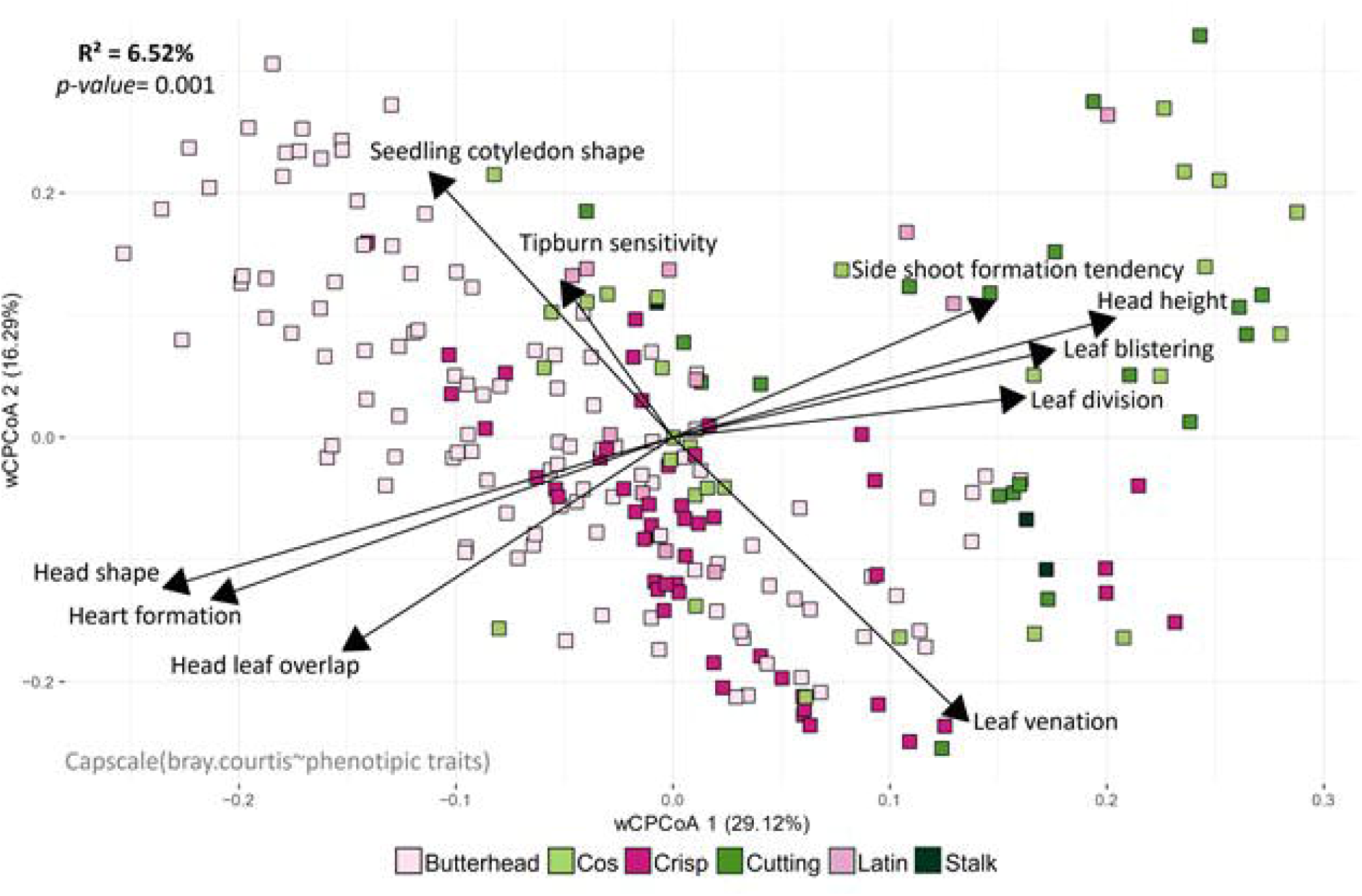
Multivariate dissection of the variety contribution to bacterial β-diversity. A) Constrained Analysis of Principal Coordinates (CAP) performed on the Bray-Curtis distance matrix illustrating how leaf morphological traits (arrows) contribute to leaf- associated bacterial β-diversity. Samples are color-coded according to the variety groups to emphasize their association with plant phenotypes. Each square represents a single sample.

To dissect the individual and combined contributions of genetic distance, variety, leaf micro- and macronutrient content, and plant morphological parameters, we conducted a Variation Partitioning analysis. The results align with those from the Constrained Analysis of Principal Coordinates, especially in terms of the variance explained by each individual trait (Fig. 5, Fig. 2D-E, 3B, and 4A). Collectively, these factors accounted for 13.9% of the total β- diversity in the leaf-associated bacterial community. Fig. 5A shows some overlap in the variance explained by these factors, but their combined effect never exceeds the contribution of individual factors (Table S13-S14).

**Fig. 5.**
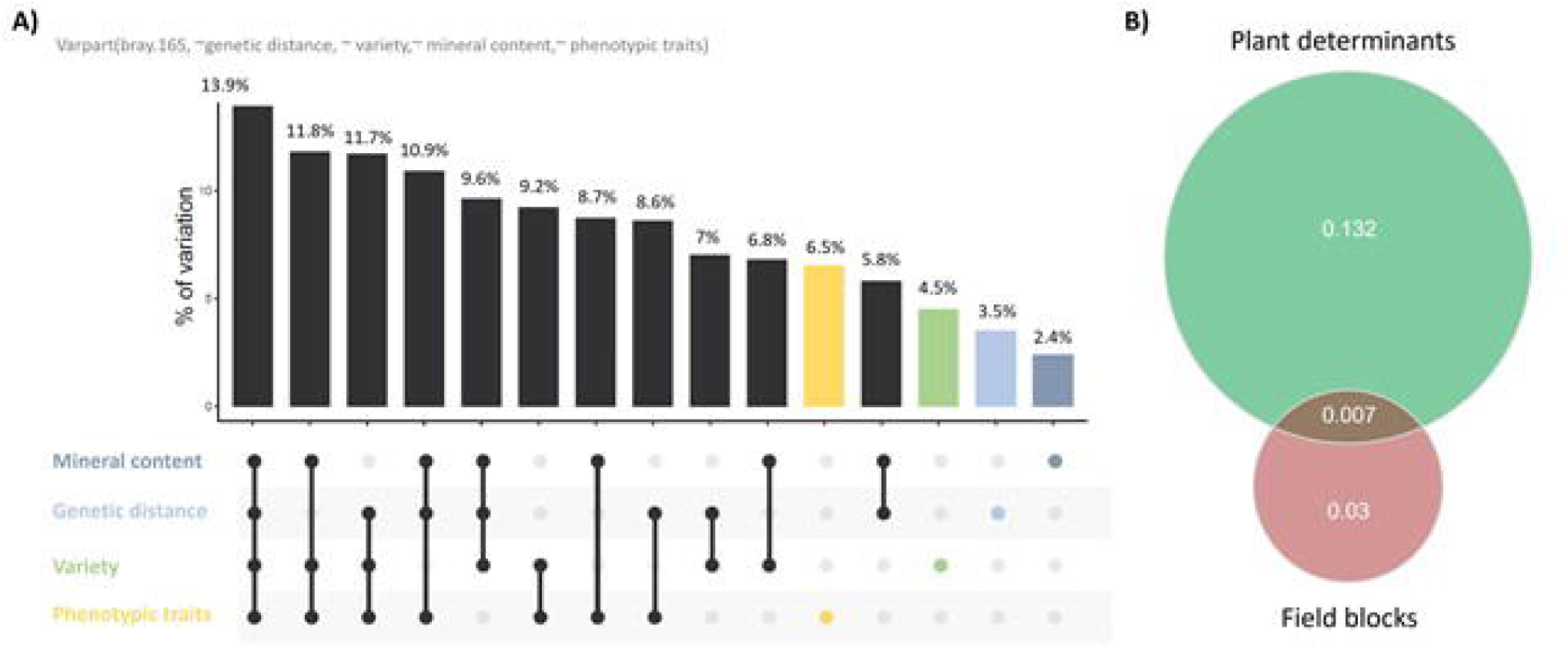
Proportion of leaf-associated bacterial 16S diversity explained by plant determinants. A) UpSet plot representing the proportion of bacterial β-diversity (R²) explained by each individual plant determinant and their combinations. Colors distinguish the contributions of individual variables from those of combined variables (in black). B) Venn diagram showing the individual contributions and the overlap of known plant and environmental factors to leaf- associated bacterial β-diversity.

In summary, these results suggest that despite the substantial environmental exposure of leaves, a significant portion of bacterial diversity is still linked to intrinsic plant characteristics. These factors influence diversity at multiple levels, including bacterial composition, α-diversity, β-diversity, with leaf morphological traits contributing with the largest share.

### Non-hub community members modulate leaf bacterial composition and richness in relation to shoot phenotypic traits

Among the significant phenotypic traits, heart formation, head height and leaf venation, which explained most of the microbiota variation and drive distinct microbial communities, were found to strongly influence α diversity (Fig. 6 and Fig. S6). The grouping values for these parameters were previously characterized by breeders. Specifically, in Fig. 6A-B, higher numbers correspond to a progressive increase in heart formation capacity and head height.

**Fig. 6.**
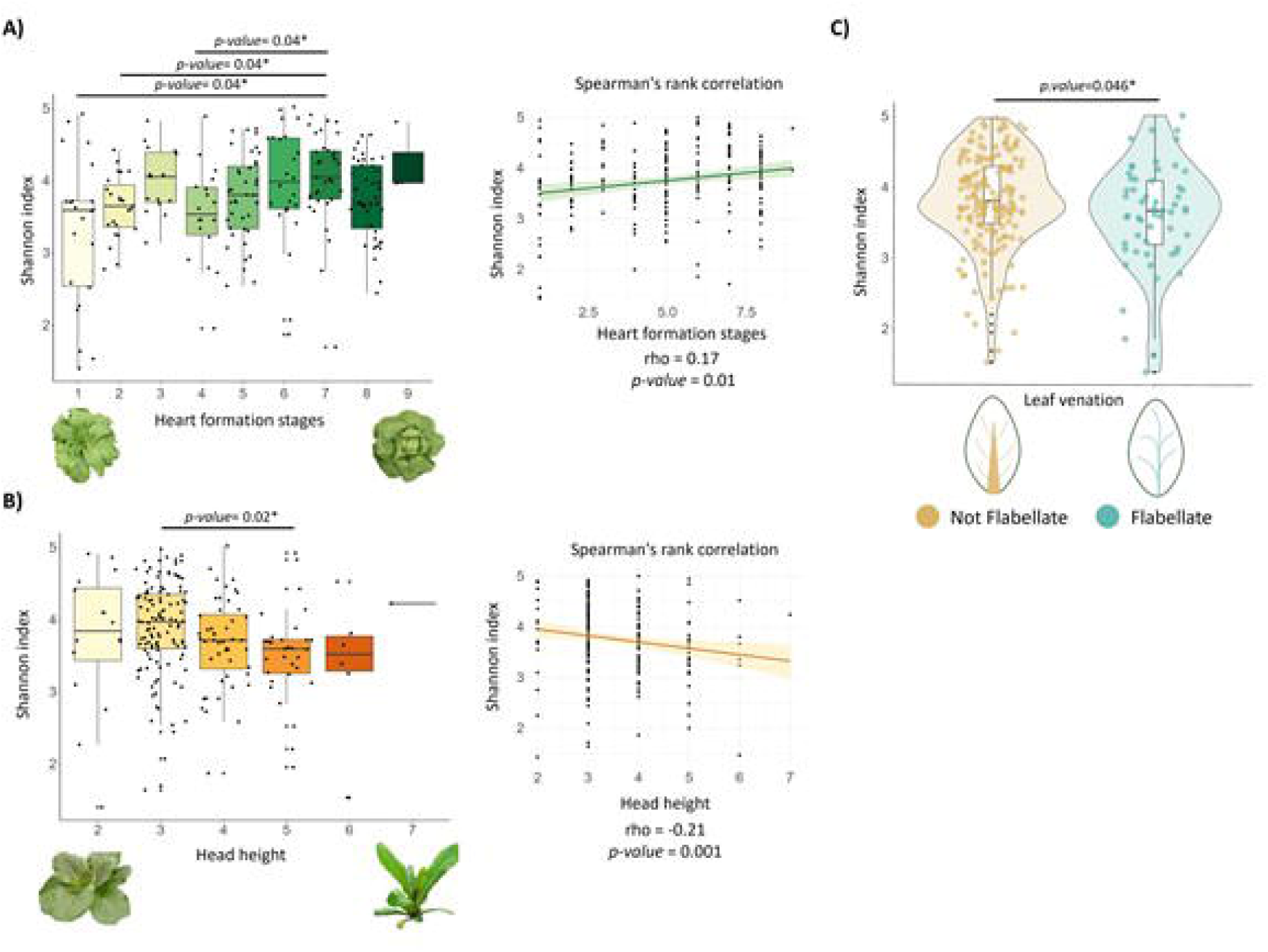
Effect of phenotypic traits on bacterial α-diversity. A-C) Differences in Shannon diversity index across A) Heart formation stages, B) Head heights, and C) Leaf venation types. Statistical differences between groups were assessed using Dunn *post hoc* and Wilcoxon–Mann–Whitney tests. Spearman’s rank correlation coefficient was used to evaluate the relationship between the tested variables and the α-diversity Shannon index in panels A and B. Credit: created in BioRender. Capparotto, A. (2024) BioRender.com/f29j677”.

After assessing the homogeneous distribution of phenotypic trait groups across field blocks (Fig. S8), we found significant differences in α-diversity (Dunn *post-hoc*, p-value <0.05) between different heart formation stages and plant heights, with a positive correlation between the Shannon index and heart formation stages (Spearman Rho = 0.17, p-value=0.01) and a negative correlation between the Shannon index and plant height (Spearman Rho = - 0.21, p-value=0.001), alongside differences in α-diversity based on head shapes (Dunn *post-hoc*, p-value <0.05, Fig. 6A-B, Fig. S6).

Additionally, leaf venation was found to impact bacterial α-diversity (Fig. 6C). Indeed, not- flabellate venation types host a bacterial community with a significantly higher Shannon index compared to flabellate types (Wilcoxon–Mann–Whitney test, p-value=0.046). Given the nutritional significance of lettuce, we further investigated the potential association between various phenotypic traits and the relative abundance of human-associated bacteria. By comparing our 16S rRNA data with the prokaryotic taxa listed in the FAPROTAX database, we found that plants with a not-flabellate venation type exhibited a greater capacity to recruit higher relative abundances of mammal gut bacteria, as well as human gut-associated and human-associated bacteria (Wilcoxon–Mann–Whitney test, p-value <0.05, Fig. S7).

To investigate which ASVs are differentially enriched between these phenotypic groups and whether these changes are driven by hub taxa within the community, we conducted differential abundance analysis (Fig. 7A) and community network analysis (Fig. 7B-C). Notably, none of the ASVs that differ in abundance between plant types are hub taxa (Fig. 7A), except for ASV59 from the Burkholderiales order, identified as a hub taxon in short plants (Fig. S9A-B), and ASV44 in not flabellate venation type (Fig. S9A-C). Across all traits analyzed, hub members mostly belong to the Parvibaculales, MBAE14, Burkholderiales, and Thiomicrospirales orders, remaining stable across phenotypic traits (Fig. 7C- S9B). The only exception is leaf venation type (Fig. S9A-C), where hub members differ between non- flabellate and flabellate groups. The non-flabellate group is entirely composed of Rhizobiales, while the flabellate group displays a pattern similar to that observed in heart formation and head height. Fig. S10 also highlights that the hierarchical cluster, according to bacterial relative abundance across heart formation stages, clearly isolates hub taxa from the remaining bacterial community.

**Fig. 7.**
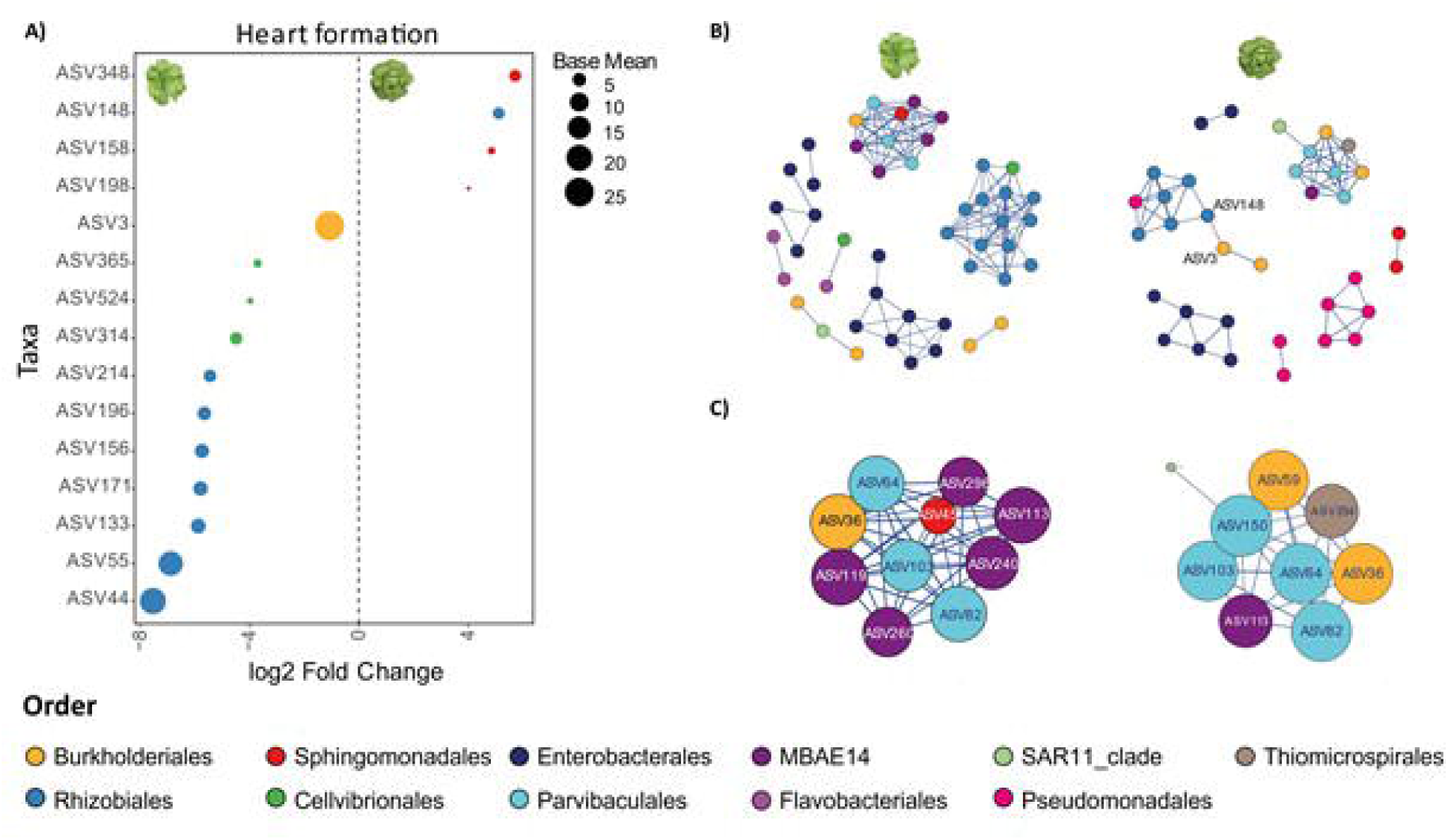
Identification of differentially abundant ASVs between heart formation capacity groups. A) ASVs with significant changes in relative abundance (p-value < 0.05) between groups 1 and 2 (open-hearted plants, left plot) and group 7 (tight-hearted plants, right plot). Colors correspond to the ASVs classification at the order level, as indicated in the legend. The size of the ASVs represents the base mean effect size, indicating the magnitude of change relative to the reference. B) Leaf bacterial community network in open-hearted plants (left) and tight-hearted plants (right). Dots represent ASVs, colored according to their order. Only significant correlations (Spearman correlation > 0.65) are depicted: positive correlations are shown in blue, and negative correlations in red. C) Hub ASVs of the community. Colors indicate the order, while size corresponds to the hub score.

Overall, these findings suggest that changes in ASV relative abundance do not involve hub taxa, which remain stable and show only minor variations between shoot morphologies. This implies that the observed diversity between groups with contrasting phenotypic traits is driven by shifts in the more abundant ASVs that are peripheral within the community network.

In summary, our findings indicate that despite the considerable environmental exposure of leaves, a significant portion of bacterial diversity is strongly associated with intrinsic plant characteristics, with leaf morphological traits playing a pivotal role. These traits influence diversity on multiple fronts, including α-diversity, β-diversity, and the enrichment of specific ASVs, without affecting hub species.

## DISCUSSION

Building on the concept that plant phenotypic traits and leaf nutrient content are direct consequences of the plant’s genetic makeup, which in turn shapes the structure of the phyllosphere bacterial community [10], we investigated the complex interplay among these factors. Specifically, we examined how different leaf physicochemical features, partially driven by genotype, influence the structure and diversity of leaf-associated bacterial communities.

Although the individual contributions of genotype, micro- and macronutrient content, and phenotypic traits may be modest, together they account for up to 13.9% of the total diversity of leaf-associated bacterial communities. This suggests that, despite the substantial influence of environmental variation on bacterial communities, plant-specific determinants still allow for the prediction of a considerable piece of diversity observed in leaves. This is in line with other works on lettuce [13, 17, 30] and Arabidopsis, where the genotype has been reported to explain nearly 10% of the variation in phyllosphere-associated bacterial communities across three studied genotypes [31]. Accordingly, GWAS on Arabidopsis reported that the host genome explained around 10% of the variance of phyllosphere bacterial and fungal communities [32, 33]. Interestingly, in our experimental setup, leaf morphological traits played a central role in setting the plant-bacterial interaction stage compared to genetic distance and leaf micro- and macronutrient content, in line with results on spinach, rocket salad and *Ipomoea hederacea* [18, 34]. This insight is crucial for breeders, as it highlights the potential to manipulate leaf microbiome through plant traits and to discriminate between leaf morphology’s direct and indirect effects, as shown by plant-derived metabolites from silver birch [35].

Among all analyzed traits, shoot morphological characteristics showed a great influence on causing shifts in both α- and β-diversity. These findings are consistent with those of [14], which highlighted the role of plant morphotype in differentiating bacterial profiles in lettuce. More specifically, our results indicate that increased heart formation capacity, decreased head heights and a not-flabellate leaf venation type are associated with greater species richness. We hypothesize that the development of a tightly formed heart may create a distinct ecological niche compared to plants with a more open structure, altering water retention and distribution, and favoring bacterial proliferation [36, 37]. In the case of low head height, the proximity to the soil can expose leaves to a greater reservoir of soil bacteria, increasing the potential for leaf bacterial contamination.

In addition to α-diversity, we reported how the different plant forms associate divergent bacterial members. For example, open-hearted plants were enriched with several ASVs from the Rhizobiales, Cellvibrionales, and Burkholderiales orders. Notably, members of the Rhizobiales and Burkholderiales orders are generally recognized as soil bacteria [38–40], suggesting that their presence in open-hearted plant types may be due to larger environmental exposure, similarly to plants grown close to the soil surface [14, 38]. Notably, two Enterobacteriaceae ASVs enriched in shorter plants belong to the genus *Pantoea*, whose members have been shown to inhibit the growth of certain human pathogens in both pre-and post-harvest conditions [41, 42]. For leaf venation, the analysis reported in Fig. S7 revealed a stronger association of mammalian gut bacteria with non-flabellate venation types, although these results require further validation through additional experiments. This suggests that vein density in non-flabellate venation types, where a midrib gives rise to other veins, may be higher, creating more favorable conditions for bacterial proliferation than in flabellate venation types, where veins grow radially from the base [43].

In accordance with the results of and colleagues [17], positive interactions between bacterial members dominated the community. The only negative interaction within the community was detected in the closed heart type (Fig. 7B) between ASV3 (from the Burkholderiales order) and ASV148 (from the Rhizobiales order). The fact that this negative interaction exists exclusively between two ASVs that are differentially enriched in the two plant forms suggests that phenotypical differences, along with the resulting physicochemical variations, may contribute to creating distinct microenvironmental niche conditions that favor specific taxa [10, 44]. At most taxonomic levels, variety was the primary plant driver of community changes; however, at the genus level, genetic distance had a greater influence, suggesting a possible role of genotype fine-tuning in shaping the leaf bacterial community [45, 46].

Overall, our findings highlight the benefits of working with large and well-documented datasets and underscore the importance of enhancing phenotyping efforts to increase the resolution of analyses. In this study, we explored the complex interconnection of three plant factors: the morphological traits of the leaves, the genotype, and the micro and macronutrients, examining their individual and combined effects on the leaf-associated bacterial communities. These observations lay the groundwork for advancing plant breeding research and developing plant varieties with an enhanced ability to support beneficial microbiota.

## Supporting information

Supplemental R markdown

## ACKNOWLEDGEMENTS

We thank Daniele Castelli for his assistance with sample collection. We are grateful to Johannes Herpell and Enrico Bortoletto for their support with data analysis. We also thank the curators of the CNG seed collection, Robbert Van Treuren and Lynn Vorstenbosch, for the lettuce seeds and the photos used in Fig. 1. We are grateful to Bronte Garden and Azienda Agricola Gambaro for providing the space and materials necessary to conduct the experiments. This work was supported by grants from the European Union (NextGenerationEU 2021 STARS Grants@Unipd program P-NICHE and Next Generation EU, Mission 4, Component 1, CUPD53D23022080001 PRIN-PNRR prot. P2022WL8TS to MG), the Ministry of the University and Research (PON Ricerca e Innovazione 2014-2020 PhD fellowship DOT1471523 to AC); the University of Padova (Progetti di Ricerca Dipartimentali - PRID, grant number BIRD214519 to MG) and by Consorzio Interuniversitario di Biotecnologie (mobility grant to AC).

## DATA AVAILABILITY STATEMENTS

The datasets generated during the current study are available in the NCBI SRA repository as Bioproject PRJNA1173114.

## SUPPLEMENTARY TABLES

Table S1. Experimental field design

Table S2. Randomized plant sample ID for endophyte extraction

Table S3. Sampling rack design

Table S4. All steps of micro- and macronutrient quantification through ICP-OES

Table S5. Sample DNA quantification

Table S6. Reads lost at each contamination removal step

Table S7. Sample size and distribution across varieties and genetic distance groups

Table S8. Leaf morphological traits before and after the NA imputation

Table S9. FEAST function output

Table S10. Statistically different taxa among varieties and genetic distance groups

Table S11. Mineral principal components contribution to bacterial β-diversity

Table S12. Single phenotypic trait contribution to β-diversity

Table S13. Varpart function output

Table S14. Statistical analysis of varpart model components

