## Supplementary figures for "Plant phenotypic differentiation outweighs genetic variation in shaping the lettuce leaf microbiota"

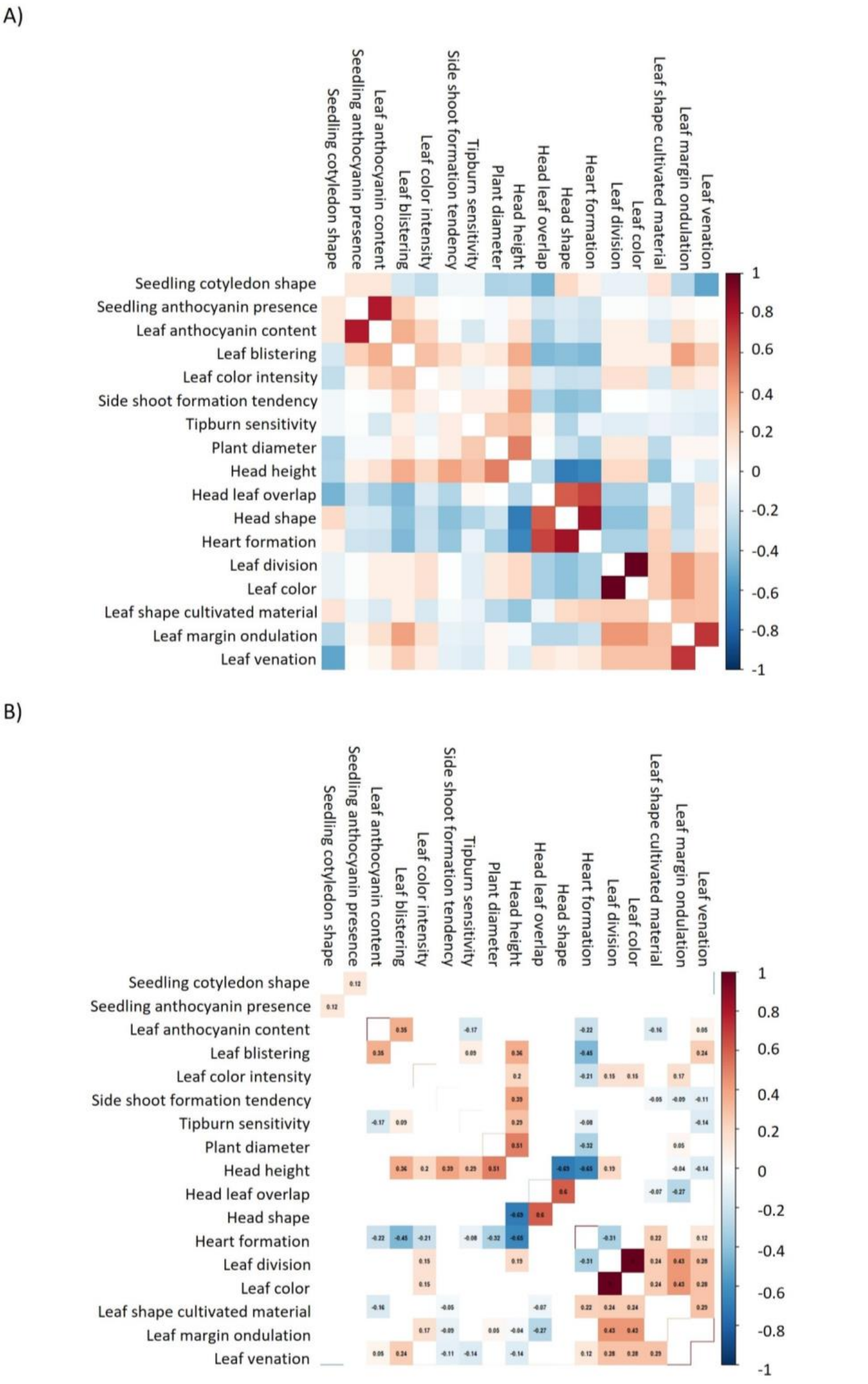


**Fig. S1. Correlation matrix between plant morphological traits before NA imputation.**

Panel A displays all relationships among the plant morphological traits, while Panel B shows only the significant correlations along with the pairwise-complete correlation coefficient calculated using the "*corr*" function in R. Imputation was based solely on the significant correlations reported in Panel B. Colors differentiate positive correlations (in red) from negative correlations (in blue), with color intensity indicating the strength of the correlation (darker colors represent stronger correlations).

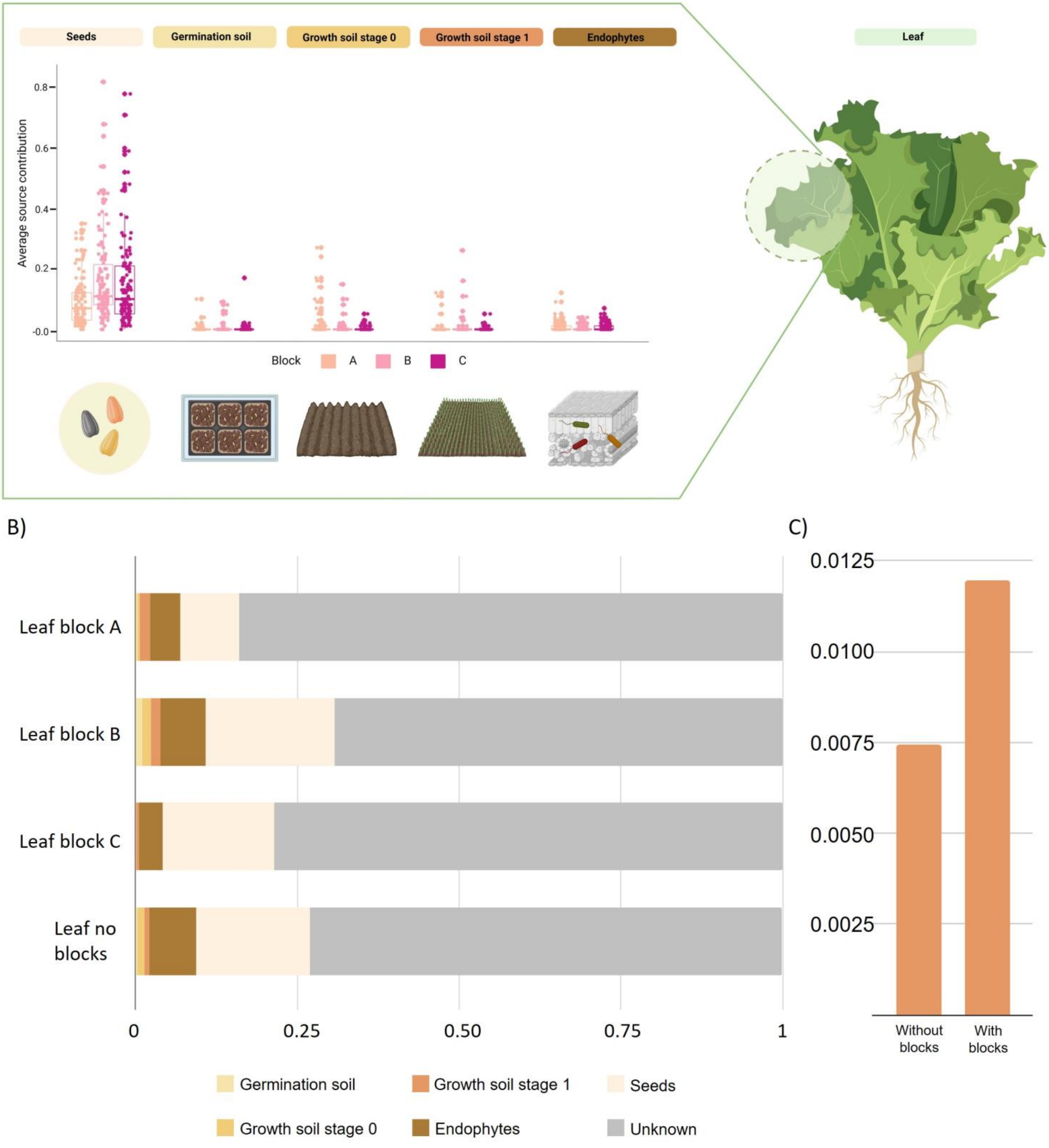


**Fig. S2. Estimated sources of the leaf-associated bacterial community.**

A) The share of each source (seeds, germination soil, growth soils, and endophytes) in contributing to the leaf bacterial community according to FEAST. Contributions were calculated separately for each sample within the three field blocks, which are represented by different colors in the boxplots. B) Proportions of known and unknown contributions based on analyses by blocks (Leaf Block A, B, and C) and a separate analysis conducted without considering field blocks (last bar). C) Variation in the average source contribution for growth stage 1, comparing results without considering blocks (first bar) and with blocks taken into account (second bar). Credit: created in BioRender. Capparotto, A. (2024) BioRender.com/a47q702 ".

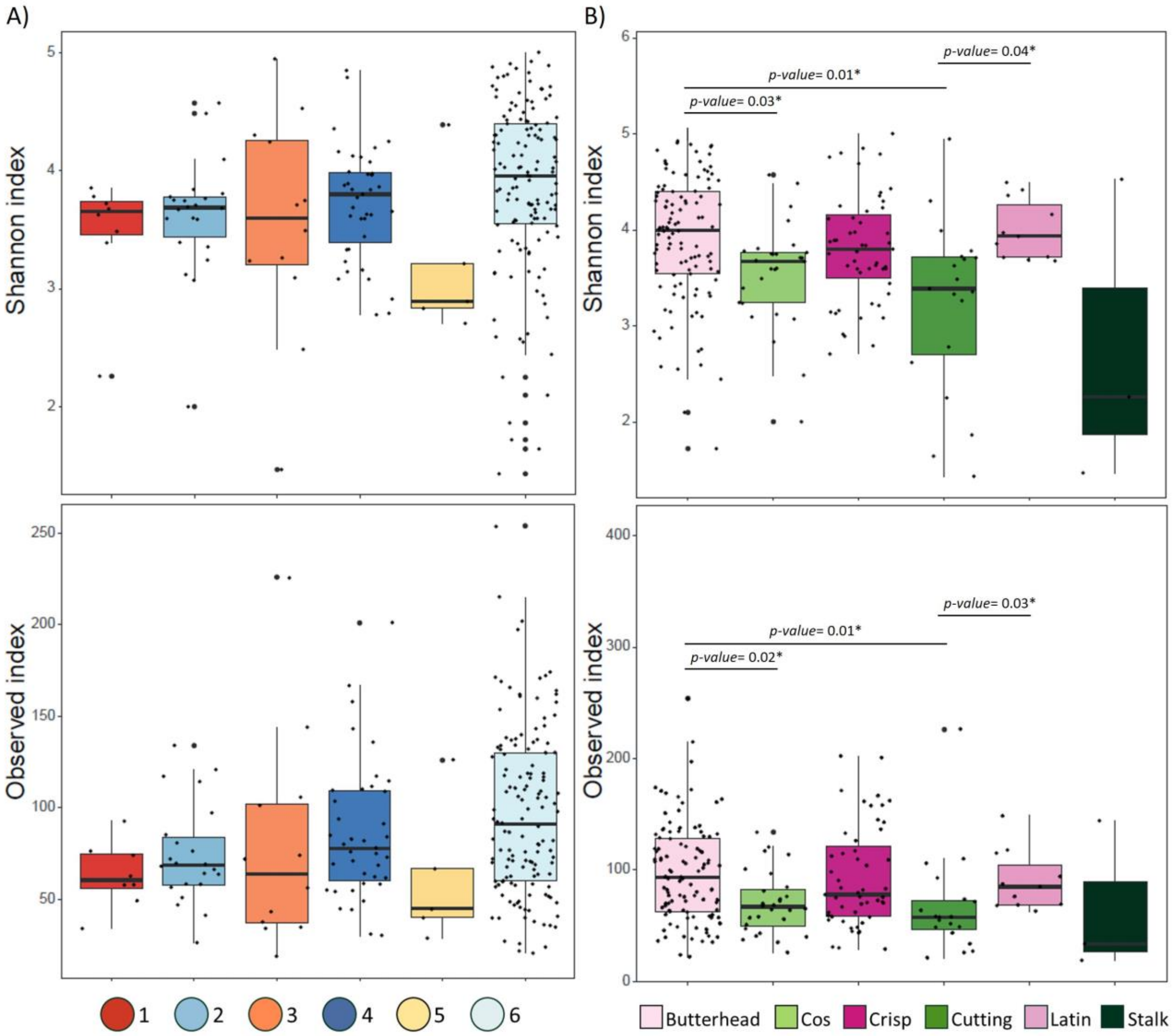


**Fig. S3. α-diversity variation across groups of closely related genotypes and varieties.**
Shannon (Above) and Observed (below) indexes were used to evaluate richness across different genetic groups (A) and varieties (B). Statistical differences between groups were evaluated by Kruskal-Wallis followed by Dunn *post-hoc* tests.

A)

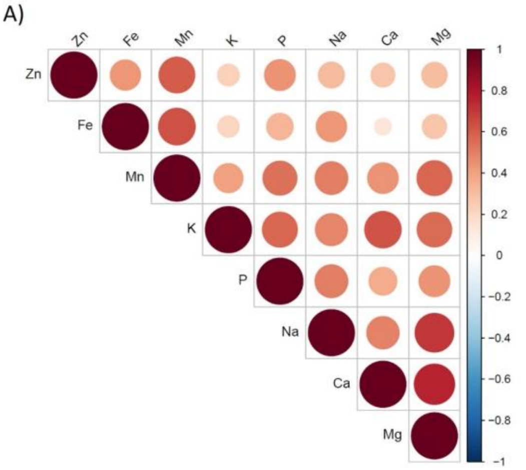


C)

Adonis2(bray.16S~element)

| Element | $R^2$ | *p-value* |
|---|---|---|
| K | ns | ns |
| Ca | ns | ns |
| Mg | ns | ns |
| Na | ns | ns |
| Fe | ns | ns |
| P | ns | ns |
| Mn | ns | ns |
| Zn | ns | ns |

B)

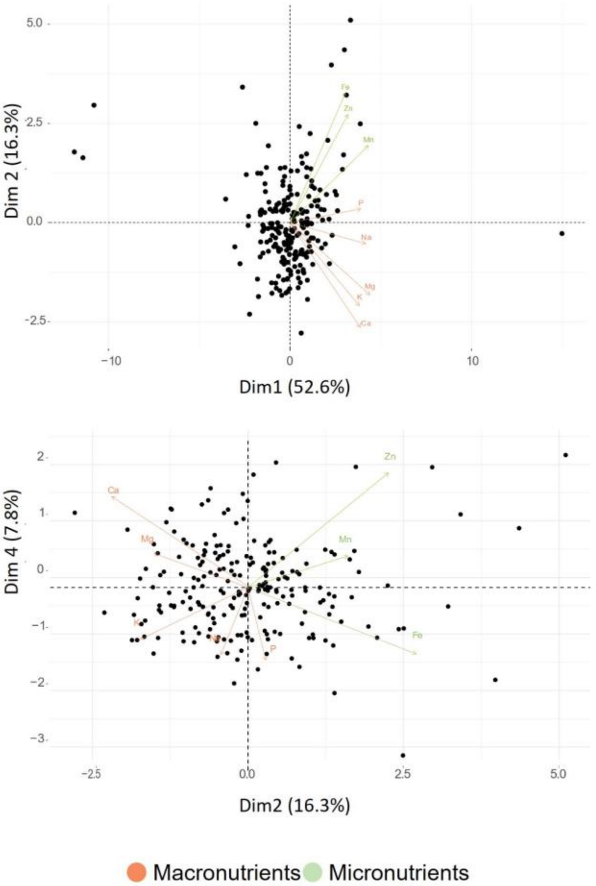


D)

Adonis2(bray.16S~variety(or genetic distance)/element)

| Element | Genetic distance | variety |
|---|---|---|
| K | ns | ns |
| Ca | ns | ns |
| Mg | ns | ns |
| Na | ns | ns |
| Fe | ns | ns |
| P | ns | $R^2$=0.034, *p-value*=0.002 |
| Mn | ns | ns |
| Zn | ns | $R^2$=0.031, *p-value*=0.014 |

**Fig. S4. Combined and independent mineral contributions to bacterial β-diversity.**
A) correlation matrix among micro- and macroelements. Colors differentiate positive correlations (in red) from negative correlations (in blue), with color intensity indicating the strength of the correlation (darker colors represent stronger correlations). B) Principal component analysis (PCA) of leaf micro- and macronutrient contents. Colors differentiate macronutrients (in pink) from micronutrients (in green). The upper panel shows the distribution along the first dimension (which does not significantly impact β-diversity) and the second dimension (which significantly influences β-diversity). The lower panel illustrates nutrient distribution according to dimensions 2 and 4, which account for significant variation in β-diversity. C) Independent mineral contributions to bacterial β-diversity. PERMANOVA analysis was performed to determine the effects of individual minerals on β-diversity. D) Nested mineral contributions to bacterial β-diversity by genetic distance and variety groups. PERMANOVA was applied to reveal effects.

A)

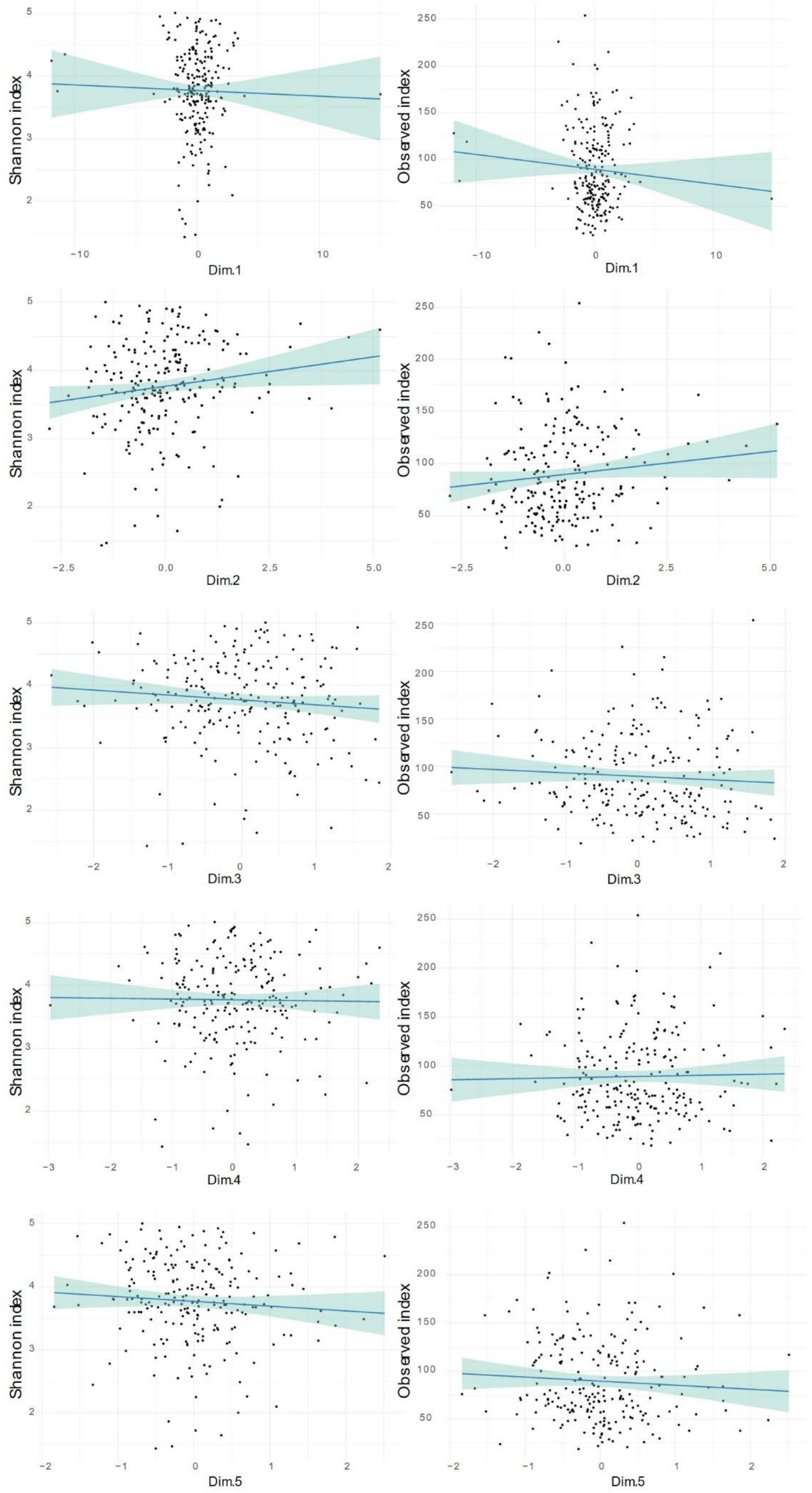


B)

| | Dim 1 | Dim2 | Dim3 | Dim4 | Dim5 |
|---|---|---|---|---|---|
| Observed | ns | *p-value* = 0.049<br>rho = 0.13 | ns | ns | ns |
| Shannon | ns | p-value = 0.019<br>rho = 0.16 | ns | ns | ns |

**Fig. S5. Correlation between mineral concentrations and bacterial α-diversity.**
In panel A, Spearman's rank correlation coefficient was used to evaluate correlations between each mineral dimension and the Shannon and Observed α-diversity indexes. Statistical analysis and Spearman's rho coefficients for each component are reported in panel B.

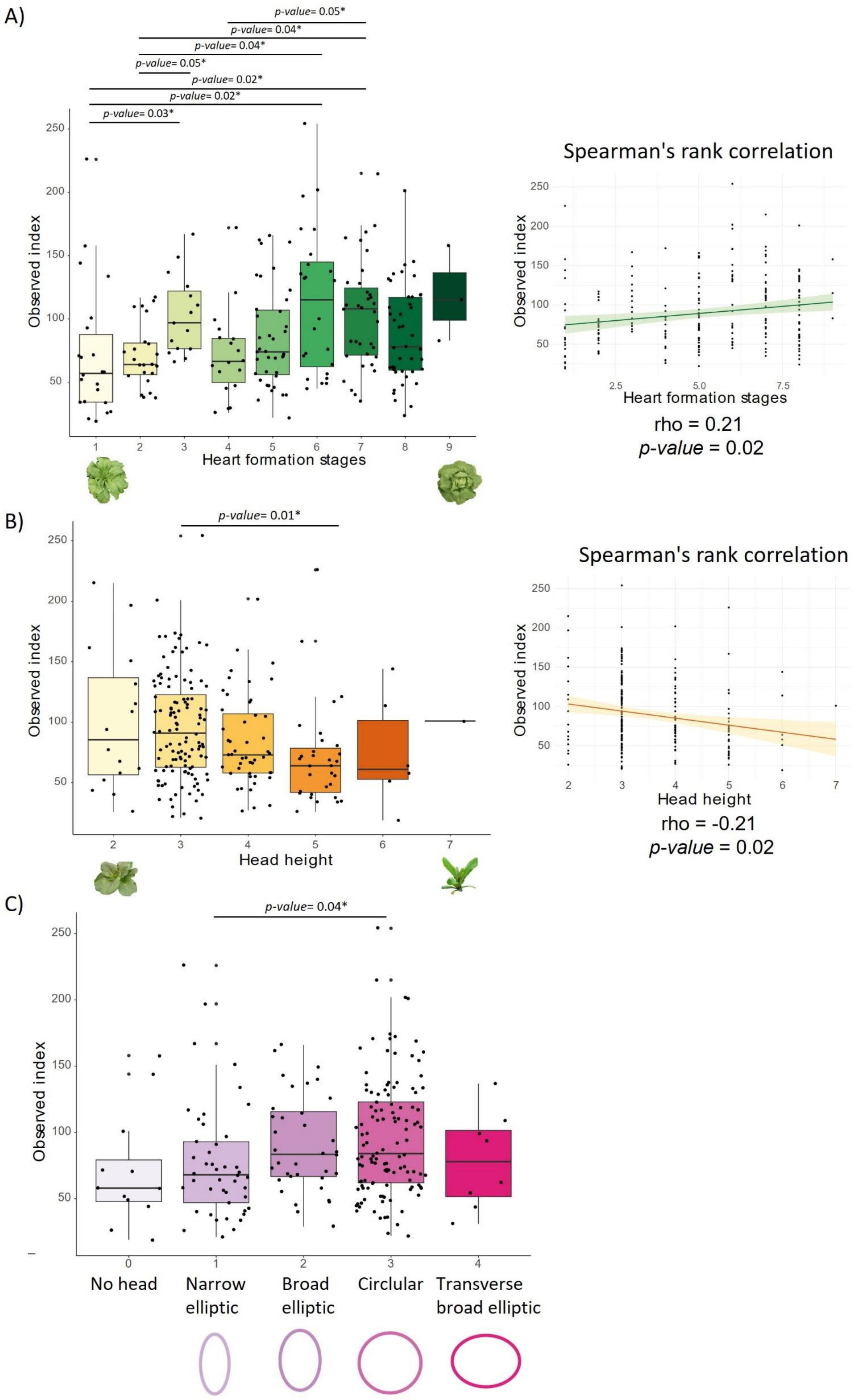


**Fig. S6. Shoot phenotypic traits strongly affect bacterial α-diversity.**

Variation in observed bacterial richness is analyzed across A) Heart formation stages B) Head height and C) Lettuce head shape type. Statistical differences between groups were calculated using the Kruskal-Wallis test followed by Dunn's *post-hoc* tests. Spearman's rank correlation, with Spearman's rho coefficient, was applied in panels A and B to evaluate correlations between: A) Different heart formation stages and B) Head heights.

A)

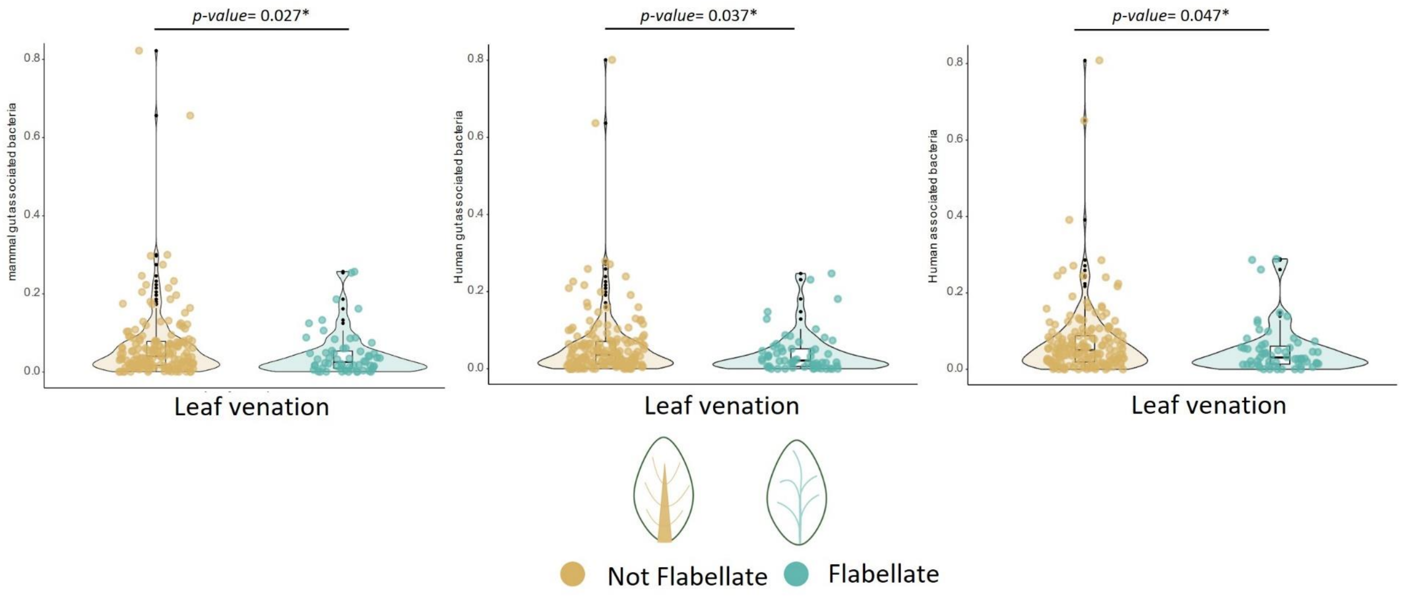


B)

| | Mammal gut bacteria | Human associated bacteria | Human gut bacteria |
|---|---|---|---|
| Seedling cotyledon shape | ns | ns | ns |
| Seedling anthocyanin presence | ns | ns | ns |
| Leaf shape cultivated material | ns | ns | ns |
| Leaf blistering | ns | ns | ns |
| Leaf margin undulation | ns | ns | ns |
| Leaf venation | *p-value*= 0,02 | *p-value*= 0,04 | *p-value*= 0,047 |
| Leaf division | ns | ns | ns |
| Leaf color | ns | ns | ns |
| Leaf color intensity | ns | ns | ns |
| Leaf anthocyanin content | ns | ns | ns |
| Side shoot formation tendency | ns | ns | ns |
| Tipburn sensitivity | ns | ns | ns |
| Plant diameter | ns | ns | ns |
| Head shape | ns | ns | ns |
| Head leafs overlap | ns | ns | ns |
| Head height | ns | ns | ns |
| Heart formation | ns | ns | ns |

**Fig. S7. Variation in leaf venation types affects the association with mammal gut bacteria.** Differences in A) the abundance of mammal gut-, human gut-associated bacteria and human-associated bacteria between flabellate and non-flabellate venation types. Statistical analyses were performed using the Kruskal-Wallis test followed by Dunn's *post-hoc* tests. Bacterial identities were extracted from the FAPROTAX database. B) The table presents the statistical analyses conducted on other plant traits.

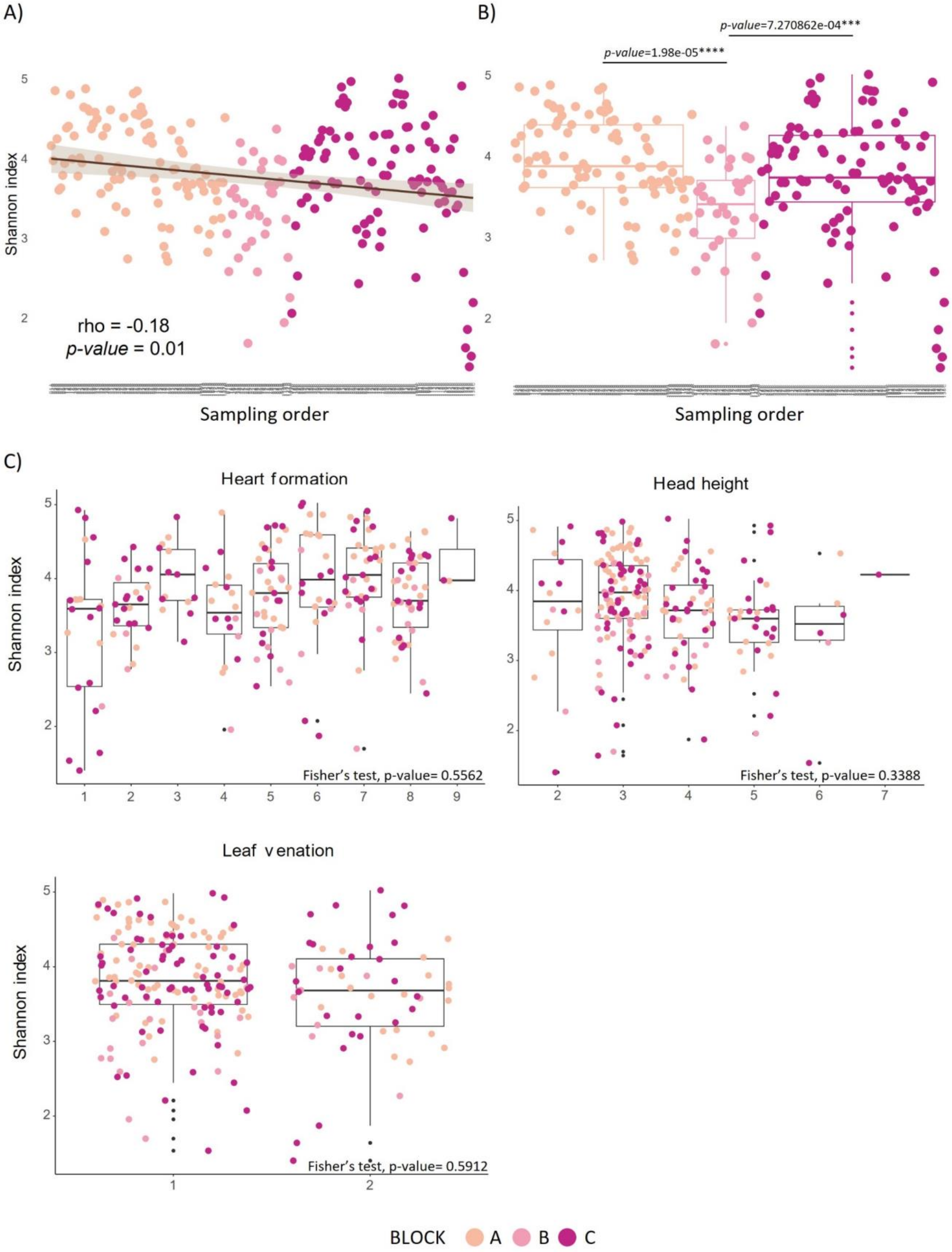


**Fig. S8. Sequential sampling influence on α-diversity.**

A) Variation in Shannon α-diversity based on sampling order (left side) and field block (right side). The X-axis shows sample IDs according to their sampling order, with colors representing different field blocks. In the left panel, Spearman's rank correlation, using Spearman's ρ coefficient, was applied to identify significant correlations. In the right panel, differences in Shannon diversity among field blocks were assessed using the Kruskal-Wallis test followed by Dunn's post-hoc tests. B) Influence of the block of origin of each sample on differences in α-diversity among the studied phenotypic traits.

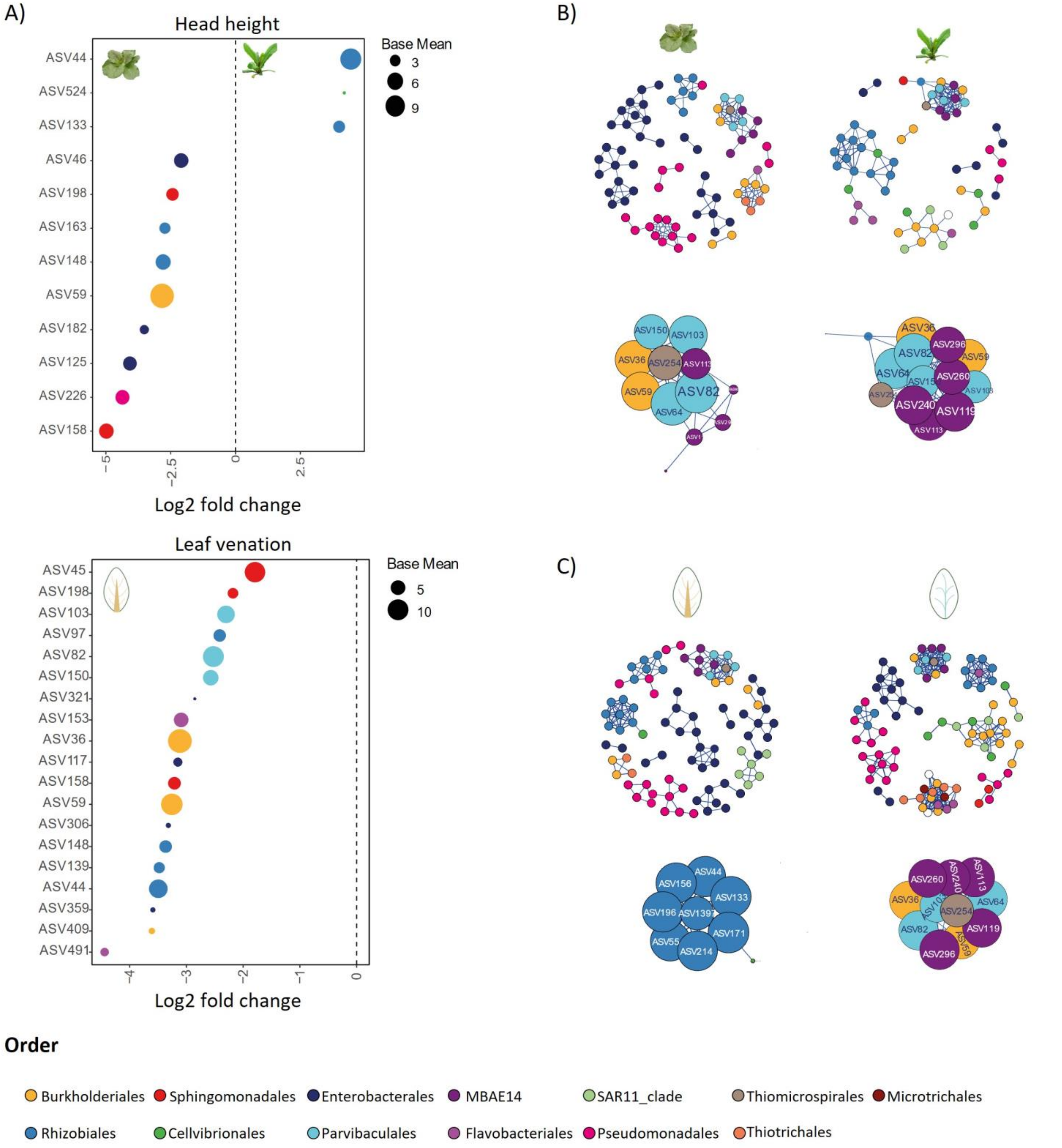


**Fig. S9. Differentially abundant ASVs between head height groups and leaf venation types.**
A) The upper panel shows ASVs with different relative abundances between head height Groups 3 (short plants, left side) and 5 (tall plants, right side). The lower panel represents ASVs with differing relative abundances between non-flabellate venation types (left side) and flabellate types (right side). Colors indicate the order level, as shown in the figure legend. The size of the dots reflects the base mean effect size. The community network for the two forms of each phenotypic trait is shown in panel B) for head height and panel C) for leaf venation. In each panel, the community network is displayed at the top, while hub community members are depicted at the bottom. Positive correlations are colored blue, negative correlations are colored red, and only significant correlations (Spearman correlation > 0.65) are shown. The size of the ASVs in the hub corresponds to the hub score.

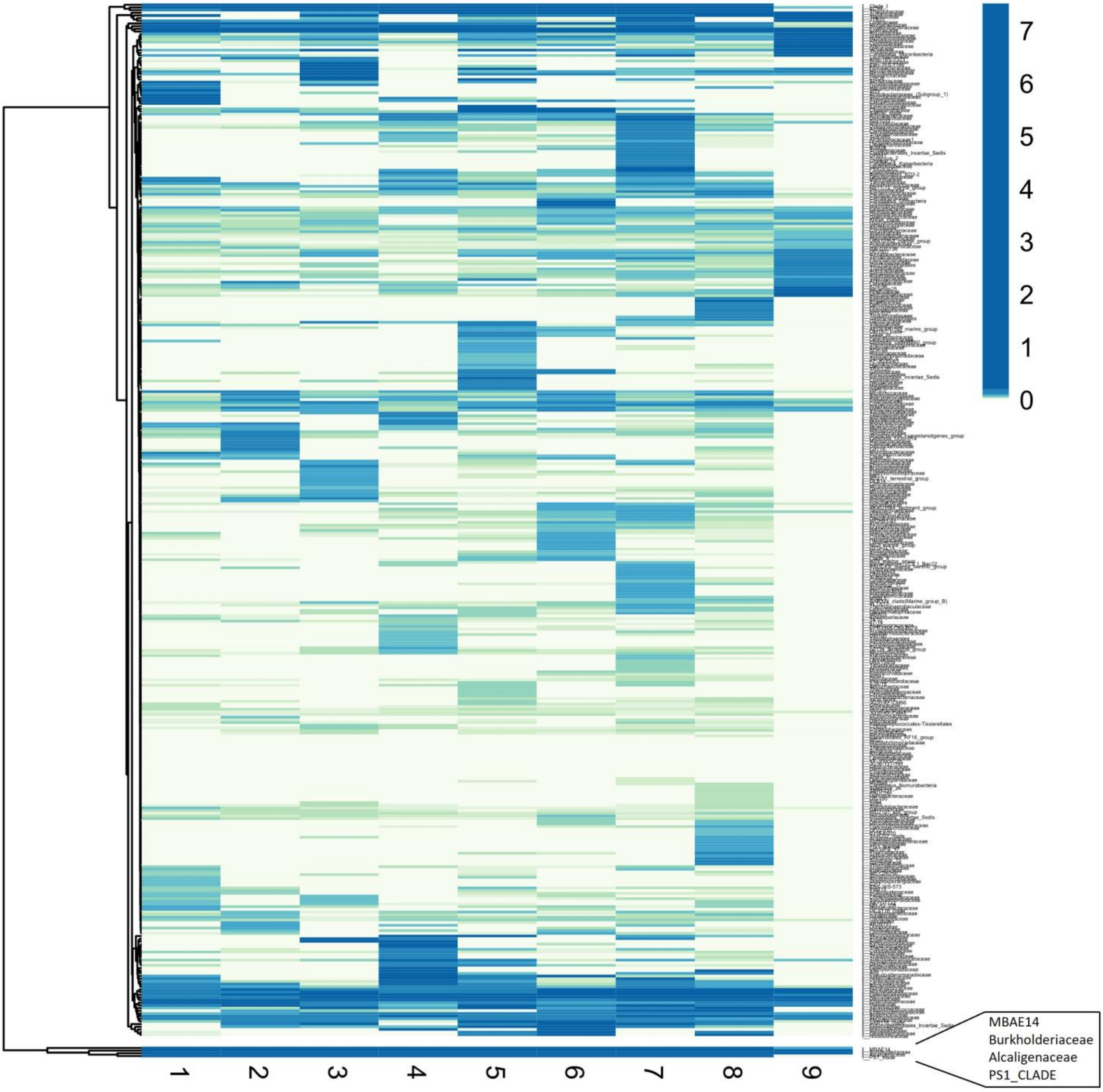


**Fig. S10. Relative abundance of leaf-associated bacterial community families across different heart formation groups.**

In the heatmap, the X-axis represents the heart formation stages, while the Y-axis shows the families, which are clustered according to their relative abundances across groups. Low relative abundances are depicted in light blue, whereas high relative abundances are indicated by dark blue.

### SUPPLEMENTARY MATERIAL AND METHODS

#### Sample collection

Using a single-hole punch, we gathered three leaf disks (3 mm diameter) from each leaf, originating from three distinct areas: the central vein, the right leaf lamina, and the apical part of the leaf. Leaf disks from plants of the same genotype and grown in the same block were combined, resulting in a total of 9 leaf disks per genotype in each block and three final replicates for each genotype. Control soil samples were collected at three different time points: on March 15$^{th}$, 2022 (germination substrate), on April 21$^{st}$, 2022 (Growth soil stage 0), and on May 21$^{st}$, 2022 (Growth soil stage 1). In the latter two instances, sample collection involved pooling samples from three different points in the field (upper, median, and lower regions). Leaf samples for endophyte control were collected on May 30$^{th}$, 2022. Specifically, a leaf from the mid-upper part of the plant was collected for each genotype following the randomized design detailed in Table S2.

#### Leaf mineral content quantification

Minerals that were poorly detected (Al, As, B, Ba, Be, Cd, Co, Cr, Cu, Li, Mo, Ni, Pb, Sb, Se, Sn) were excluded from the analysis, and only the predominantly detected ones (Ca, Fe, K, Mg, Mn, Na, P, Zn) were considered. For the latter, samples falling below the limit of quantification (<LOQ) were fit in the model, following the procedure reported in a published method (Stuart L. Beal 2001).

In brief, the LOQ was initially calculated using equation 1 (Eq.1):

Eq 1. $LOQ = \frac{\left(\frac{A(mg)\ x\ 20\ ml}{1000\ ml}\right)}{B}$

Where 'A' represents the concentration of the initial point on the linear absorbance curve (mg/L), which is specific to each mineral; '20 ml' means the volume used for sample dilution; and 'B' denotes the average weight of the leaves (0.002132 g)."

Finally, the concentration of these samples (mg/g) was calculated following equation 2 (Eq.2):

Eq 2. $[Sample] = \frac{LOQ}{\sqrt{2}}$

#### DNA extraction and sequencing

To isolate endophytes, under biological hood conditions, leaves were first submerged in 70% ethanol (2 minutes), followed by immersion in 5% hypochlorite (5 minutes), and then in 70% ethanol (30 seconds). Finally, leaves were washed twice in sterile water before being homogenized with a sterile blender.

All DNA samples underwent MiSeq Illumina 16S amplicon sequencing at IGA Technology Services (IGATech, Udine, Italy). To summarize the process briefly, 2 × 250 bp stretches of 16S rDNA spanning the V3-V4 region were amplified using the 16S-341F (5’- CCTACGGGNBGCASCAG -3’) and 16S-805R (5’- GACTACNVGGGTATCTAATCC -3’) primers. During the initial amplification step, PNA clamping was applied to block the amplification of host chloroplast and mitochondrial 16S sequences, adhering to the manufacturer's protocol (PNA Bio Inc, Newbury Park, CA).

**Raw reads processing**

Forward and reverse reads were filtered and trimmed at 250 bp before merging. The merged reads underwent chimera removal and taxa assignment utilizing the Silva database (Silva database trained on the V3-V4 region, 2023). Only highly abundant Amplicon Sequence Variants (ASVs) with a relative abundance greater than 0.1% and samples with at least 100 reads were retained for subsequent analyses.

**Community composition and contributions of plant determinants to β diversity**

For a matter of visualization, the arrows in the Constrained Analysis of Principal Coordinates (CAP) were scaled down by a factor of 2.

Given the relatively minor impact of individual micronutrients and macronutrients on β diversity and the strong correlation existing among them, we chose to extract their principal components (PCs). Principal component analysis (PCA) was thus conducted in R, utilizing the Factoextra and FactoMineR packages. Only significant plant determinants were integrated into the varpart function following testing against the Bray-Curtis distance matrix.

**Integrative analysis of leaf bacterial communities: functional predictions, diversity, networks, and origins**

For the log2foldchange function application, variables were transformed into a binary format to include only those groups where significant α diversity differences were detected. Samples with a p-value $< 0.05$ were considered.

To enhance the community network representation using igraph we applied different minimum prevalence thresholds (the number of samples in which ASVs were not expressed): 6 for both heart formation groups, 10 for plants with high head heights, 4 for those with short head heights, 12 for the non-flabellate leaf venation type, and 5 for the flabellate type. Spearman's correlation coefficient was employed to assess the relationships between taxa within the network, and only significant correlations (Spearman's correlation $> 0.65$) were included in the final image.
