## Supplemental R markdown for "Plant phenotypic differentiation outweighs genetic variation in shaping the lettuce leaf microbiota": Supplementary script.html


### Supplementary script

###### Arianna Capparotto et al

#### 2024-08-01

```
knitr::kable(head(mtcars), "latex", longtable = TRUE)
```

**1-Processing and clean data from QIIME2**

##### 1.1 - Denoising table from DADA2.

```
Denoising <- read.csv("yourPath/Denoising.tsv", header = TRUE, sep = "\t", dec = ".")

Denoising
```

##### 1.2 - Construct phyloseq objects

```
#libraries

library("phyloseq")
library("ggplot2")      
library("readxl")       
library("dplyr")       
library("tibble")
library("tidyverse")
library("magrittr")


#Upload tables

##ASV Table
asv16S <- read.table(file="yourPath/16S_table-FINAL.txt",sep="\t",dec = ".",header=TRUE) 
##Tax Table
tax16S <- read.table(file="yourPath/taxonomy.tsv",sep="\t",dec = ".",header=TRUE)
##Design
  design <- read.table(file="yourPath/Design.txt",sep="\t",dec = ",",header=TRUE)

##Sequences
library(Biostrings)
refseq16S <- readDNAStringSet("yourPath/dna-sequences.fasta")

#Manage data for Phyloseq
asv16S <- asv16S %>%
    tibble::column_to_rownames("OTU.ID") 
tax16S <- tax16S %>% 
    tibble::column_to_rownames("OTU.ID")

#Collapse Tax Table
library(stringr)
tax16S[c("Kingdom","Phylum", "Class", "Order", "Family", "Genus", "Species")] 
<- str_split_fixed(tax16S$Taxon, ';', 7)
tax16S[tax16S == ""] <- NA 
tax16S[tax16S == " "] <- NA  
design <- design %>% 
    tibble::column_to_rownames("sampleID") 

otu_mat <- as.matrix(asv16S)
tax_mat <- as.matrix(tax16S)

#Create Phyloseq Object  
ps16S_0 <- phyloseq(tax_table(tax_mat),
                  otu_table(otu_mat, taxa_are_rows = TRUE), sample_data(design), refseq(refseq16S))
```

##### 1.3 - Mitocondria, chloroplast and Archaea Contaminations removal

```
# TAXA Calculate the number of taxa before filtering
num_taxa_before <- ntaxa(ps16S_0)


# Filtered ASV based on taxonomy
ps16S_1a <- subset_taxa(ps16S_0, !is.na(Taxon) & !Taxon %in% c("Unassigned"))
ps16S_1b <- subset_taxa(ps16S_1a, !Order %in% c(" o__Chloroplast"))
ps16S_1c <- subset_taxa(ps16S_1b, !Order %in% c("o__Mitochondria"))
ps16S_1d <- subset_taxa(ps16S_1c, !Family %in% c("f__Mitochondria"))
ps16S_1e <- subset_taxa(ps16S_1d, !Family %in% c("f__Chloroplast"))
ps16S_1f <- subset_taxa(ps16S_1e, !Genus %in% c(" g__Mitochondria"))
ps16S_1g <- subset_taxa(ps16S_1f, !Taxon %in% c("d__Eukaryota"))
ps16S_1h <- subset_taxa(ps16S_1g, !Kingdom %in% c("d__Eukaryota"))
ps16S_1i <- subset_taxa(ps16S_1h, !Kingdom %in% c("d__Archaea"))
```

##### 1.4 - Remove blank samples contamination

```
################## decontam
library(here)
packageVersion("here")
# 1.0.1
library(decontam)
packageVersion("decontam")
packageVersion("decontam")
# 1.16.0
library(phyloseq)
packageVersion("phyloseq")
# 1.40.0
library(Biostrings)
packageVersion("Biostrings")
# 2.64.0
library(tidyverse)
packageVersion("tidyverse")
# 1.3.2

d16S_leaves <- subset_samples(ps16S_1i, tissue %in% c("leaf"))
d16S_leaves <- subset_taxa(d16S_leaves, !taxa_sums(d16S_leaves) == 0)
d16S_soil <- subset_samples(ps16S_1i, tissue %in% c("soil"))
d16S_soil <- subset_taxa(d16S_soil, !taxa_sums(d16S_soil) == 0)
d16S_endophytes <- subset_samples(ps16S_1i, tissue %in% c("endophytes"))
d16S_endophytes <- subset_taxa(d16S_endophytes, !taxa_sums(d16S_endophytes) == 0)
d16S_seeds <- subset_samples(ps16S_1i, tissue %in% c("seed"))
d16S_seeds <- subset_taxa(d16S_seeds, !taxa_sums(d16S_seeds) == 0)

# import clean phyloseq from blanks
phyloseq_bianchi <- readRDS("ps_blanks.rds")

# prepare dataset
ps16S_1i.2 <- merge_phyloseq(d16S_leaves, phyloseq_bianchi)
sample_data(ps16S_1i.2)$FEAST[394:400] <- paste("blank")
base::as.data.frame(phyloseq::sample_data(ps16S_1i.2))
base::as.data.frame(phyloseq::sample_data(d16S_leaves))
view(sample_data(ps16S_1i.2))

# inspect libraries sizes

library("gtable")

df <- as.data.frame(sample_data(ps16S_1i.2))
df$LibrarySize <- sample_sums(ps16S_1i.2)
df <- df[order(df$LibrarySize), ]
df$Index <- seq(nrow(df))
df$FEAST <- as.factor(df$FEAST)
ggplot(data = df, aes(x = Index, y = LibrarySize, color = FEAST)) + geom_point() +
    theme(legend.title = element_text(size = 5), legend.text = element_text(size = 5),
        legend.key.size = unit(0.5, "cm"), legend.key.width = unit(0.5, "cm"))

# Prevalence based-decontamination

sample_data(ps16S_1i.2)$is.neg <- sample_data(ps16S_1i.2)$FEAST == "blank"

contamdf.prev <- isContaminant(ps16S_1i.2, method = "prevalence", neg = "is.neg")
table(contamdf.prev$contaminant)

# Make phyloseq object of presence-absence in negative controls and true
# samples
ps.pa <- transform_sample_counts(ps16S_1i.2, function(abund) 1 * (abund > 0))
ps.pa.neg <- prune_samples(sample_data(ps.pa)$FEAST == "blank", ps.pa)
ps.pa.pos <- prune_samples(sample_data(ps.pa)$FEAST == "leaf", ps.pa)
# Make data.frame of prevalence in positive and negative samples
df.pa <- data.frame(pa.pos = taxa_sums(ps.pa.pos), pa.neg = taxa_sums(ps.pa.neg),
    contaminant = contamdf.prev$contaminant)
ggplot(data = df.pa, aes(x = pa.neg, y = pa.pos, color = contaminant)) + geom_point() +
    xlab("Prevalence (Negative Controls)") + ylab("Prevalence (True Samples)")

# trim contaminants

ps.noncontam <- prune_taxa(!contamdf.prev$contaminant, ps16S_1i.2)
ps.noncontam

ps16S_1i.3 <- merge_phyloseq(ps.noncontam, d16S_endophytes, d16S_seeds, d16S_soil)
view(otu_table(ps16S_1i.3))

ps16S_1i.3 = subset_samples(ps16S_1i.3, FEAST != "blank")
```

##### 1.5 - Create final tables resuming the process

```
num_taxa_after1 <- data.frame(Before_filtering = ntaxa(ps16S_0), Remove_Unassigned = ntaxa(ps16S_1a),
    remove_chloroplast = ntaxa(ps16S_1e), remove_mithocondria = ntaxa(ps16S_1f),
    remove_Eukaryota = ntaxa(ps16S_1h), remove_Archaea = ntaxa(ps16S_1i), romove_blanks = ntaxa(ps16S_1i.3))

# READS Before after filtering Calculate the total number of reads per sample
total_reads <- colSums(otu_table(ps16S_0))
total_reads_Unassigned <- colSums(otu_table(ps16S_1a))
total_reads_Chloro <- colSums(otu_table(ps16S_1e))
total_reads_Mito <- colSums(otu_table(ps16S_1f))
total_reads_Eukaria <- colSums(otu_table(ps16S_1h))
total_reads_Archaea <- colSums(otu_table(ps16S_1i))
total_reads_blanks <- colSums(otu_table(ps16S_1i.3))


# Gather
total_filtering_reads <- cbind(row.names(total_reads), total_reads, total_reads_Unassigned,
    total_reads_Chloro, total_reads_Mito, total_reads_Eukaria, total_reads_Archaea,
    total_reads_blanks)
total_filtering_reads_f <- merge(total_filtering_reads, design, by = "row.names")
write.csv(total_filtering_reads_f, file = "total_filtering_reads_f.csv", row.names = FALSE)
write.csv(total_filtering_reads, file = "total_filtering_reads.csv", row.names = FALSE)


# Create a data frame with sample IDs and total reads
reads_per_sample_table <- data.frame(SampleID = sample_names(ps16S_1i.3), TotalReads = total_reads,
    TotalReads_Unassigned = total_reads_Unassigned, TotalReads_Bacterial = total_reads_Chloro,
    total_reads_Eukaria = total_reads_Eukaria, total_reads_blanks = total_reads_blanks,
    Percent_Bacteria = 100 * total_reads_blanks/total_reads)
```

##### 1.6 - Remove samples with less than 100 counts and set a relative abundance

```
# Rename ASV
dna.16S <- taxa_names(ps16S_1i.3)
names(dna.16S) <- taxa_names(ps16S_1i.3)
ps16S_2 <- merge_phyloseq(ps16S_1i.3, dna.16S)
taxa_names(ps16S_2) <- paste0("ASV", seq(ntaxa(ps16S_2)))

# Subset Samples (remove sample with less than 100 counts)
ps16S_3 <- prune_samples(sample_sums(ps16S_2) >= 100, ps16S_2)
ps16S_4 <- filter_taxa(ps16S_3, function(x) sum(x) > 0, TRUE)

# Filter ASV (keep ASV > 0.1% RA per sample)
ps16S_5 <- transform_sample_counts(ps16S_4, function(x) ifelse((x/sum(x)) >= 0.001,
    x, 0))
ps16S_6 <- filter_taxa(ps16S_5, function(x) sum(x) > 0, TRUE)
otu_table16S <- data.frame(otu_table(ps16S_6))
ps16S_6_w <- ps16S_6

saveRDS(ps16S_5, "ps16S_less_clean_noblanks.rds")
saveRDS(ps16S_6, "ps16S_clean_noblanks.rds")
```

##### 1.7 - phylogenetic tree

```
# create phylogenetic tree Neighbor-Joining tree were constructed the packages
# DECIPHER v 2.12.0 and Phangorn v 2.5.5 using default paramaters.

library("DECIPHER")
packageVersion("DECIPHER")

seqs.16S <- refseq(ps16S_6)
alignment.16S <- AlignSeqs(DNAStringSet(seqs.16S), anchor = NA)
library("phangorn")
packageVersion("phangorn")
phang.align.16S <- phyDat(as(alignment.16S, "matrix"), type = "DNA")
dm.16S <- dist.ml(phang.align.16S)  #measure of distances 
treeNJ.16S <- NJ(dm.16S)  # Note, tip order != sequence order
fit.16S <- pml(treeNJ.16S, data = phang.align.16S)
fit.GTR.16S <- update(fit.16S, k = 4, inv = 0.2)
fit.GTR.16S <- optim.pml(fit.GTR.16S, model = "GTR", optInv = TRUE, optGamma = TRUE,
    rearrangement = "stochastic", control = pml.control(trace = 0))
d16S.tree <- phyloseq(tax_table(ps16S_6), sample_data(ps16S_6), otu_table(ps16S_6,
    taxa_are_rows = FALSE), refseq(ps16S_6), phy_tree(treeNJ.16S))
saveRDS(d16S.tree, "16S_PS_clean_tree.rds")


# import phylogenetic tree
library(ape)
treeNJ.16S <- readRDS("treeNJ.16S.decontam.rds")

# join the phylogenetic tree

ps16S_6 <- phyloseq(tax_table(ps16S_6), sample_data(ps16S_6), otu_table(ps16S_6),
    refseq(ps16S_6), phy_tree(treeNJ.16S))

write.tree(phy_tree(ps16S_6), "output.tree.noblanks")

saveRDS(ps16S_6, "ps16S_clean_noblanks_withtree.rds")
```

##### 1.8 - Size dependent rarefaction for compartment

```
# create a dataset for each compartment

# leaf
d16S_filtered_leaves <- subset_samples(ps16S_6_w, tissue %in% c("leaf"))
d16S_filtered_leaves <- subset_taxa(d16S_filtered_leaves, !taxa_sums(d16S_filtered_leaves) ==
    0)

# seed
d16S_filtered_seed <- subset_samples(ps16S_6_w, tissue %in% c("seed"))
d16S_filtered_seed <- subset_taxa(d16S_filtered_seed, !taxa_sums(d16S_filtered_seed) ==
    0)

# soil
d16S_filtered_soil <- subset_samples(ps16S_6_w, tissue %in% c("soil"))
d16S_filtered_soil <- subset_taxa(d16S_filtered_soil, !taxa_sums(d16S_filtered_soil) ==
    0)

# endophytes
d16S_filtered_endophyte <- subset_samples(ps16S_6_w, tissue %in% c("endophytes"))
d16S_filtered_endophyte <- subset_taxa(d16S_filtered_endophyte, !taxa_sums(d16S_filtered_endophyte) ==
    0)

# rarefy each compartment

library("ranacapa")

r16S_leaf <- rarefy_even_depth(d16S_filtered_leaves, sample.size = 1000, rngseed = 900)

r16S_soil <- rarefy_even_depth(d16S_filtered_soil, sample.size = 14000, rngseed = 900)

r16S_seeds <- rarefy_even_depth(d16S_filtered_seed, sample.size = 200, rngseed = 900)

r16S_endophytes <- rarefy_even_depth(d16S_filtered_endophyte, sample.size = 4500,
    rngseed = 900)

# create the merged phyloseq object with each rarefied compartment

phy_tree <- readRDS("treeNJ.16S.decontam.rds")
merged_phylo <- merge_phyloseq(r16S_leaf, r16S_soil, r16S_seeds, r16S_endophytes)
vector_asv <- row.names(otu_table(merged_phylo))
vector_sample <- row.names(sample_data(merged_phylo))
phy_tree_filtered <- prune_taxa(vector_asv, phy_tree)
merged_phylo_tree <- phyloseq(tax_table(merged_phylo), sample_data(merged_phylo),
    otu_table(merged_phylo), refseq(merged_phylo), phy_tree(phy_tree))

# visualize the output of rarefaction

rcurve16S <- ggrare(merged_phylo_tree, color = "tissue", step = 999, se = FALSE) +
    labs(title = "16S rRNA gene")
rcurve16S

rcurve16Sf_rarefied <- rcurve16S + theme(strip.background = element_rect(color = "white",
    fill = "#585858", size = 1, linetype = "solid"), strip.text.x = element_text(size = 12,
    color = "white"), axis.title.x = element_text(size = 12, face = "bold"), axis.title.y = element_text(size = 12,
    face = "bold"), plot.title = element_text(size = 16), axis.text.x = element_text(face = "bold"),
    axis.text.y = element_text(face = "bold")) + ylab("Species richness") + xlab("Sample size") +
    ggtitle("16S rRNA gene") + facet_grid(~tissue, scale = "free")

rcurve16Sf_rarefied
```

***2-Community composition*** # 2- Community
composition ### 2.1 - Figure 2A

```
library(ggplot2)

#The most abundnat members of the community (Relative abundance >0.1%)
# Prepare the data
data <- data.frame(
  Phylum = c("Proteobacteria", "Bacteroidota", "Firmicutes", "Actinobacteriota", "Unassigned", 
             "Planctomycetota", "Verrucomicrobiota", "Desulfobacterota", "Campilobacterota", 
             "Acidobacteriota", "Chloroflexi", "Cyanobacteria", "Bdellovibrionota", "Patescibacteria"),
  Percentage = c(71.31652174, 8.385217391, 6.978695652, 5.0765217390,3.416956522, 1.089130435, 1.065217391, 0.517826087, 0.466086957, 0.399565217, 0.311304348, 0.28826087, 0.276086957, 0.106521739))


my_colors <- c("#E41A1C", "#377EB8", "#4DAF4A", "#984EA3", "#FF7F00",
               "#FFFF33", "#A65628", "#F781BF", "#999999", "#66C2A5",
               "#FC8D62", "#8DA0CB", "#E78AC3", "#A6D854")

# Reorder the levels of Phylum based on their abundance
data$Phylum <- factor(data$Phylum, levels = data$Phylum[order(data$Percentage, decreasing = T)])

# Create the stacked bar plot
stacked_ggp <- ggplot(data, aes(x = "", y = Percentage, fill = Phylum, group = Phylum)) +
  geom_bar(stat = 'identity', position = 'stack', width = 1) +
  scale_fill_manual(values = my_colors) +
  guides(fill = guide_legend(title = "Factors Explaining Variation")) +
  labs(title = "Stacked Bar Plot of Phylum Percentages") +
  theme_bw() +
  theme(panel.border = element_blank(),
        panel.grid.major = element_blank(),
        panel.grid.minor = element_blank(),
        legend.position = "right")

stacked_ggp
```

##### 2.2 - Figure 2B- - Statistical analysis to find taxa differences between varieties and groups of closely related genotypes

```
#To do this part, the community threshold (relative abundance >0.1%) has been recalculated using only leaf samples, as follows:" 

ps16S_3<-subset_samples(ps16S_6_w, tissue %in% c("leaf"))
ps16S_4 <- filter_taxa(ps16S_3, function(x) sum(x) > 0, TRUE)

#And a new phyloseq object (r16S_leaf) have been created with the same procedure reported above (Part 1), containing only leaf samples. 

#Obtain a table made of OTU + TAXA table to calculate the relative abundance at each single taxonomic rack (phylum, class, order, family, genus, species)

#convergence table
taxa.table_1<-tax_table(r16S_leaf)
taxa.table_1<-as.data.frame(taxa.table_1)
otu.table_1<-as.data.frame(otu_table(r16S_leaf))
convergence_table_1<- (cbind(taxa.table_1[4:9], otu.table_1))
write.csv(convergence_table_1, "convergence_table_1.csv")
write.csv(samples, "sample_data_leaf.csv")

#Then, create the tables (one for each taxonomic rank) by following these instructions: 

  #1) Open the Excel file and create a pivot table from the converged table by clicking on "Recommended Pivot Table." Choose the pivot table with the desired taxonomic rank (e.g., Phylum).

  #2) Select all samples.

  #3) Copy the table into a new Excel file, transposing the entire table.

  #4) Calculate the relative abundance using the formula:

  #Relative abundance = Count in each cell/ Sum of all counts per sample
 
  #5) Ensure that the sum of the relative abundances for each sample equals 1.

  #6) Save the table as .csv file.

#Import data and perform statistical analysis by running the following code:

#libraries
library(ggExtra)
library(multcomp)
library(ggpubr)
library(rstatix)
library(patchwork)
library(tidyverse)

#Import dataset - change every time the table imported at the taxonomic levels you want to perform statistic
  data <- read.csv("C:/yourPath/species.csv", #phylum, order, class, family, genus
                 header=T,sep = ";", na.string=c("na", "NA", "null")) 

#convert the table to have your taxa in a single column
data %>%
 pivot_longer(., cols = (2:294),  names_to= "species" , values_to = "Abundance")-> data2 

#tranform to factor
data2$genotype<-as.factor(data2$genotype)
data2$species <- as.factor(data2$species) #change with the taxa you are examining
data2$variety <- as.factor(data2$variety)
data2$genetic.distance <- as.factor(data2$genetic.distance)

#Statistic - change "species" with the taxa you are examining and "genetic distance" with the factor for which you want to assess differences

#Krukall Wallis test
x <- unique(data2$species)
models_genetic.distance  <- sapply(x, function(My){
  kruskal.test(Abundance~ genetic.distance, data=data2, species== My)
}, simplify=FALSE)
models_genetic.distance 

table_genetic.distance  <- do.call(rbind.data.frame, models_genetic.distance )
table_genetic.distance 

#Dunn Post hoc test
data2 %>% 
  group_by(species) %>% 
  dunn_test(Abundance~ genetic.distance, p.adjust.method = "fdr") -> dunns_species_genetic.distance

write.csv(dunns_species_genetic.distance, "dunns_species_genetic.distance.csv")

#Now move to phyton3 and GraPhlAn graphical interface. 
    #1 - form the table " "dunns_species_genetic.distance" select all the taxa for which there is significant differences 
    #2 - Create a table with unique significant taxa names
    #3 - Create figure 2B-C by following the script of Segata et al., 2015
```

***3-Beta diversity analysis*** # 3- Beta
diversity analysis ### 3.1 - Figures 2D-E

```
#library
library(vegan)

#Create a subset of the phyloseq object to include only leaf samples
r16S_L <- subset_samples(merged_phylo_tree, tissue %in% c("leaf"))
r16S_L <- subset_taxa(r16S_L, !taxa_sums(r16S_L) == 0)
r16S_L= subset_samples(r16S_L, genotype2 != "11") #no in nature tree
r16S_L= subset_samples(r16S_L, genotype2 != "71") #no belonging to genetic group
r16S_L= subset_samples(r16S_L, genotype2 != "119") #no belonging to genetic group
r16S_L= subset_samples(r16S_L, genotype2 != "92") #Too much Na values 

#PERMANOVA statistical analysis (Adonis2)

bray.16S <- phyloseq::distance(r16S_L, method = "bray")
mdat.16S_bray <- as(sample_data(r16S_L), "data.frame")

adonis.16STiss_bray <- adonis2(bray.16S ~ genetic.distance, #change to variety
                                data = mdat.16S_bray, perm = 999) 
print(adonis.16STiss_bray)

#Figure 2E-F - Constrained Analysis of Principal Coordinates (CAP) 

Cap_cluster<- capscale(bray.16S~genetic.distance,#change to variety
                       data = mdat.16S_bray, add=FALSE, subset, na.action = na.omit)

##ANOVA-like permutation analysis
perm_test <- anova(Cap_cluster, permutations = 999)
p_value <- perm_test$Pr[1]
p_value

##Run variability table function
variability_table <- function(cca){

        chi <- c(cca$tot.chi,
                       cca$CCA$tot.chi, cca$CA$tot.chi)
        variability_table <- cbind(chi, chi/chi[1])
        colnames(variability_table) <- c("inertia", "proportion")
        rownames(variability_table) <- c("total", "constrained", "unconstrained")
        return(variability_table)
}


##Generate variability table
var_tbl <- variability_table(Cap_cluster)
ci <- quantile(perm_test$F.perm, c(.05,.95))*perm_test$chi[1]/var_tbl["total", "inertia"]
eig_CCA <- Cap_cluster$CCA$eig
eig_CA <- Cap_cluster$CA$eig
variance <- var_tbl["constrained", "proportion"]
variance_NMDS <- var_tbl["unconstrained", "proportion"]

##Extract unweighted and  unweighted CCA values and unconstrained CA values (here you basically extract the positions of your samples in the ordination space defined by the constrained/unconstrained axes)

###Extract colnames
colnames <- c(paste("u", colnames(Cap_cluster$CCA$u)[1], sep=""), paste("u", colnames(Cap_cluster$CCA$u)[2], sep=""), paste("w", colnames(Cap_cluster$CCA$wa)[1], sep=""), paste("w", colnames(Cap_cluster$CCA$wa)[2], sep=""), paste(colnames(Cap_cluster$CA$u)[1], sep=""), paste(colnames(Cap_cluster$CA$u)[2], sep=""))
  
 ### Extract datapoints 
points = data.frame(c(Cap_cluster$CCA$u[, 1]),
                    c(Cap_cluster$CCA$u[, 2]),
                    c(Cap_cluster$CCA$w[, 1]),
                    c(Cap_cluster$CCA$w[, 2]),
                    c(Cap_cluster$CA$u[, 1]),
                    c(Cap_cluster$CA$u[, 2]))
colnames(points) <- colnames
rownames(points)


##Calculate axis explained variation
eig_tot <- sum(Cap_cluster$CCA$eig)
eigenvalues <- Cap_cluster$CCA$eig/eig_tot
eigenvalues

##Bind with metadata


df <- cbind(points, mdat.16S_bray[match(rownames(points), rownames(mdat.16S_bray)),])


#Extract Arrows

#script for extracting genetic distance arrows
arrow <- data.frame(Cap_cluster$CCA$biplot)
Cap_cluster2<- capscale(bray.16S ~ genetic.distance-1,  data = mdat.16S_bray, add=FALSE,na.action=na.exclude , subset)
genetic.distance1<- data.frame(Cap_cluster2$CCA$biplot)[1,]
arrows2 <- rbind(arrow,genetic.distance1)
new_row_names <- sub("genetic.distance", "", row.names(arrows2))
row.names(arrows2)<-new_row_names
arrows2$names <- row.names(arrows2)


#script for extracting variety arrows
Cap_cluster3<- capscale(bray.16S ~ variety-1,  data = mdat.16S_bray, add=FALSE,na.action=na.exclude , subset)
varietyButterhead<- data.frame(Cap_cluster3$CCA$biplot)[1,]
arrows2 <- rbind(arrow,varietyButterhead)
new_row_names <- sub("variety", "", row.names(arrows2))
row.names(arrows2)<-new_row_names

#Colored palette

#genetic distance
colors <- c("#d73027","#91bfdb", "#fc8d59","#4575b4", "#fee090","#e0f3f8")

#Variety
colors_var<- c( "#fde0ef","#a1d76a","#c51b7d","#4d9221","#e9a3c9","#013220","#e6f5d0")


#Plot
pwCPOA <- ggplot(df, aes(x=wCAP1, y=wCAP2)) +
  geom_point(aes(fill=genetic.distance), shape=22, size=3.5, na.value = "grey50") +
  scale_fill_manual(values = colors) + #colors_var for variery
  labs(
    x = paste("wCPCoA 1 (", format(100 * eig_CCA[1] / sum(eig_CCA), digits=4), "%)", sep=""),
    y = paste("wCPCoA 2 (", format(100 * eig_CCA[2] / sum(eig_CCA), digits=4), "%)", sep="")
  ) +
  ggtitle(paste(format(100 * variance, digits=3), " % of variance; p=", format(p_value, digits=2), sep="")) +
  theme(
    legend.position = "right",
    panel.grid.major = element_line(color = "#EDEDED"),  
    panel.grid.minor = element_line(color = "#EDEDED"),  
    panel.background = element_rect(fill = "white"),   
    plot.background = element_rect(fill = "white")     
  ) +
  geom_segment(data = arrows2, aes(x = 0, y = 0, xend = CAP1, yend = CAP2), size = 0.5, 
               arrows2 = arrow(type = "closed", length = unit(0.2, "inches"))) +
  geom_text(data = arrows2, aes(x = CAP1, y = CAP2, label = row.names(arrows2)), vjust = -1.5, hjust = NA)


pwCPOA
```

##### 3.2 - Figures 3A and S4A-B

```
#libraries
library("FactoMineR")
library("factoextra")

#transform to factor and numeric
mdat.16S_bray$variety<-as.factor(mdat.16S_bray$variety)
mdat.16S_bray$genetic.distance<-as.factor(mdat.16S_bray$genetic.distance)
mdat.16S_bray$block<-as.factor(mdat.16S_bray$block)
mdat.16S_bray$P<-as.numeric(mdat.16S_bray$P)
mdat.16S_bray$Ca<-as.numeric(mdat.16S_bray$Ca)
mdat.16S_bray$Fe<-as.numeric(mdat.16S_bray$Fe)
mdat.16S_bray$K<-as.numeric(mdat.16S_bray$K)
mdat.16S_bray$Mg<-as.numeric(mdat.16S_bray$Mg)
mdat.16S_bray$Mn<-as.numeric(mdat.16S_bray$Mn)
mdat.16S_bray$Na<-as.numeric(mdat.16S_bray$Na)
mdat.16S_bray$Zn<-as.numeric(mdat.16S_bray$Zn)

#adjust the table with the right row.names
row.names(mdat.16S_bray)<-NULL
row.names(mdat.16S_bray)<- mdat.16S_bray$sampleID3
view(mdat.16S_bray)

# log transform mineral data 
mdat.16S_bray_active <- log(mdat.16S_bray[20:27])

#Extract principal components
res.pca <- PCA(mdat.16S_bray_active, graph = FALSE)
print(res.pca)
var <- get_pca_ind(res.pca)
var<-as.data.frame(var$coord)

mineral_content<-var[, c("Dim.2", "Dim.4", "Dim.5"), drop = FALSE] #to use in varpart

#Figure S3A- correlation matrix among micro- and macronutrients

library(gplots)
cor_matrix <- cor(mdat.16S_bray_active, use = "complete.obs")

#color palette
default_colors <- colorRampPalette(c("#053061", "#2166AC", "#4393C3", "#92C5DE", "#D1E5F0", 
                                     "#FFFFFF", "#FDDBC7", "#F4A582", "#D6604D", "#B2182B", 
                                     "#67001F"))(200)

#plot
corrplot(cor_matrix, type = "upper", order = "hclust", col=default_colors,
         tl.col = "black", tl.srt = 45)


#Figure 3A - visualize Correlations between variables and dimensions
#library
library("corrplot")

default_colors <- colorRampPalette(c("#053061", "#2166AC", "#4393C3", "#92C5DE", "#D1E5F0", 
                                     "#FFFFFF", "#FDDBC7", "#F4A582", "#D6604D", "#B2182B", 
                                     "#67001F"))(200)

corrplot(res.pca$var$cor, is.corr=FALSE, col=default_colors, tl.cex=0.8, cl.cex=0.8, addrect=2)

#Figure S4A

fviz_pca_biplot(res.pca,
                axes = c(1,2),
                legend.title = "Mineral variation",
                col.var = "black",
                pointshape = 19,
                pointsize = 2.5,
                labelsize = 3,
                label = "var",
                title = "Mineral variation"
)

#Figure S4A
fviz_pca_biplot(res.pca,
                axes = c(2,4),
                legend.title = "Mineral variation",
                col.var = "black",
                pointshape = 19,
                pointsize = 2.5,
                labelsize = 3,
                label = "var",
                title = "Mineral variation"
)
```

##### 3.3 - Figures 3B, S4D, and Table S10

```
#add mineral content coordinates to mdat.16S_bray table 
mdat.16S_bray <- as(sample_data(r16S_L), "data.frame")
mdat.16S_bray$sampleID  <- rownames(mdat.16S_bray)
rownames(mdat.16S_bray) <- NULL
mdat.16S_bray <- cbind(sampleID = mdat.16S_bray$sampleID, mdat.16S_bray[, -ncol(mdat.16S_bray)])
mdat.16S_bray$sample_ID <- NULL
sampleID3_values <- rownames(mdat.16S_bray)
var <- var %>% mutate(sampleID3 = rownames(var))
mdat.16S_bray <- left_join(mdat.16S_bray, var, by = "sampleID3")
rownames(mdat.16S_bray) <- mdat.16S_bray[,1]

#Tranfrom mineral into numeric
mdat.16S_bray$P<-as.numeric(mdat.16S_bray$P)
mdat.16S_bray$Ca<-as.numeric(mdat.16S_bray$Ca)
mdat.16S_bray$Fe<-as.numeric(mdat.16S_bray$Fe)
mdat.16S_bray$K<-as.numeric(mdat.16S_bray$K)
mdat.16S_bray$Mg<-as.numeric(mdat.16S_bray$Mg)
mdat.16S_bray$Mn<-as.numeric(mdat.16S_bray$Mn)
mdat.16S_bray$Na<-as.numeric(mdat.16S_bray$Na)
mdat.16S_bray$Zn<-as.numeric(mdat.16S_bray$Zn)
mdat.16S_bray$Dim.1<-as.numeric(mdat.16S_bray$Dim.1)
mdat.16S_bray$Dim.2<-as.numeric(mdat.16S_bray$Dim.2)
mdat.16S_bray$Dim.3<-as.numeric(mdat.16S_bray$Dim.3)
mdat.16S_bray$Dim.4<-as.numeric(mdat.16S_bray$Dim.4)
mdat.16S_bray$Dim.5<-as.numeric(mdat.16S_bray$Dim.5)

#Table S10 - PERMANOVA (Adonis2) to test significance of dimensions

adonis.16Sb_phenotypic_traits <- adonis2(bray.16S~Dim.2, #Change dimension
                                data = mdat.16S_bray, perm = 999, na.action=na.exclude )
adonis.16Sb_phenotypic_traits

#Figure S4D - nested PERMANOVA (Adonis 2)
adonis.16Sb_phenotypic_traits <- adonis2(bray.16S~genetic.distance/Fe, #Change genetic distance with variety and Fe with another micro/macro nutrient
                                data = mdat.16S_bray, perm = 999, na.action=na.exclude )
adonis.16Sb_phenotypic_traits

#Figure 3B - Constrained Analysis of Principal Coordinates (CAP) 

Cap_cluster<- capscale(bray.16S~Dim.2+Dim.4+Dim.5,#Only significant dimensions
                       data = mdat.16S_bray, add=FALSE, subset, na.action = na.omit)

##ANOVA-like permutation analysis
perm_test <- anova(Cap_cluster, permutations = 999)
p_value <- perm_test$Pr[1]
p_value

##Run variability table function
variability_table <- function(cca){

        chi <- c(cca$tot.chi,
                       cca$CCA$tot.chi, cca$CA$tot.chi)
        variability_table <- cbind(chi, chi/chi[1])
        colnames(variability_table) <- c("inertia", "proportion")
        rownames(variability_table) <- c("total", "constrained", "unconstrained")
        return(variability_table)
}


##Generate variability table
var_tbl <- variability_table(Cap_cluster)
ci <- quantile(perm_test$F.perm, c(.05,.95))*perm_test$chi[1]/var_tbl["total", "inertia"]
eig_CCA <- Cap_cluster$CCA$eig
eig_CA <- Cap_cluster$CA$eig
variance <- var_tbl["constrained", "proportion"]
variance_NMDS <- var_tbl["unconstrained", "proportion"]

##Extract unweighted and  unweighted CCA values and unconstrained CA values (here you basically extract the positions of your samples in the ordination space defined by the constrained/unconstrained axes)

###Extract colnames
colnames <- c(paste("u", colnames(Cap_cluster$CCA$u)[1], sep=""), paste("u", colnames(Cap_cluster$CCA$u)[2], sep=""), paste("w", colnames(Cap_cluster$CCA$wa)[1], sep=""), paste("w", colnames(Cap_cluster$CCA$wa)[2], sep=""), paste(colnames(Cap_cluster$CA$u)[1], sep=""), paste(colnames(Cap_cluster$CA$u)[2], sep=""))
  
 ### Extract datapoints 
points = data.frame(c(Cap_cluster$CCA$u[, 1]),
                    c(Cap_cluster$CCA$u[, 2]),
                    c(Cap_cluster$CCA$w[, 1]),
                    c(Cap_cluster$CCA$w[, 2]),
                    c(Cap_cluster$CA$u[, 1]),
                    c(Cap_cluster$CA$u[, 2]))
colnames(points) <- colnames
rownames(points)


##Calculate axis explained variation
eig_tot <- sum(Cap_cluster$CCA$eig)
eigenvalues <- Cap_cluster$CCA$eig/eig_tot
eigenvalues

##Bind with metadata


df <- cbind(points, mdat.16S_bray[match(rownames(points), rownames(mdat.16S_bray)),])


#Extract Arrows

#script for extracting genetic distance arrows
arrow <- data.frame(Cap_cluster$CCA$biplot)

#Plot
pwCPOA <- ggplot(df, aes(x=wCAP1, y=wCAP2)) +
  geom_point(aes(fill=genetic.distance), shape=22, size=3.5, na.value = "grey50") +
  scale_fill_manual(values = colors_var) + 
  labs(
    x = paste("wCPCoA 1 (", format(100 * eig_CCA[1] / sum(eig_CCA), digits=4), "%)", sep=""),
    y = paste("wCPCoA 2 (", format(100 * eig_CCA[2] / sum(eig_CCA), digits=4), "%)", sep="")
  ) +
  ggtitle(paste(format(100 * variance, digits=3), " % of variance; p=", format(p_value, digits=2), sep="")) +
  theme(
    legend.position = "right",
    panel.grid.major = element_line(color = "#EDEDED"),  
    panel.grid.minor = element_line(color = "#EDEDED"),  
    panel.background = element_rect(fill = "white"),   
    plot.background = element_rect(fill = "white")     
  ) +
  geom_segment(data = arrow, aes(x = 0, y = 0, xend = CAP1, yend = CAP2), size = 0.5, 
               arrow = arrow(type = "closed", length = unit(0.2, "inches"))) +
  geom_text(data = arrow, aes(x = CAP1, y = CAP2, label = row.names(arrow)), vjust = -1.5, hjust = NA)


pwCPOA
```

##### 3.4 - Figure S1

```
#Figure S1- Na inputation through phenotypic traits correlation

##1) transform every phenotypic trait as numeric
design$Seedling.cotiledon.shape<-as.numeric(design$Seedling.cotiledon.shape)
design$Seedling.anthocyanin.presence<-as.numeric(design$Seedling.anthocyanin.presence)
design$Leaf.shape.cultivated.material<-as.numeric(design$Leaf.shape.cultivated.material)
design$Leaf.blistering<-as.numeric(design$Leaf.blistering)
design$Leaf.margin.undulation<-as.numeric(design$Leaf.margin.undulation)
design$Leaf.venation<-as.numeric(design$Leaf.venation)
design$Leaf.division<-as.numeric(design$Leaf.division)
design$Leaf.color<-as.numeric(design$Leaf.division)
design$Leaf.color.intensity<-as.numeric(design$Leaf.color.intensity)
design$Leaf.anthocyanin.content<-as.numeric(design$Leaf.anthocyanin.content)
design$Side.shoot.formation.tendency<-as.numeric(design$Side.shoot.formation.tendency)
design$Tipburn.sensitivity<-as.numeric(design$Tipburn.sensitivity)
design$Plant.diameter<-as.numeric(design$Plant.diameter)
design$Head.shape<-as.numeric(design$Head.shape)
design$Head.leafs.overlap<-as.numeric(design$Head.leafs.overlap)
design$Head.height<-as.numeric(design$Head.height)
design$Heart.formation<-as.numeric(design$Heart.formation)
design$variety<-as.factor(design$variety)

##calculate correlations 
library(corrplot)
M = cor(design[25:41], use='pairwise.complete.obs')
corrplot(M, method = 'color', order = 'alphabet')
corrplot(M, method = 'number',number.cex = 0.4) # colorful number

## calculate p-values
cor.mtest <- function(mat, ...) {
    mat <- as.matrix(mat)
    n <- ncol(mat)
    p.mat<- matrix(NA, n, n)
    diag(p.mat) <- 0
    for (i in 1:(n - 1)) {
        for (j in (i + 1):n) {
            tmp <- cor.test(mat[, i], mat[, j], ...)
            p.mat[i, j] <- p.mat[j, i] <- tmp$p.value
        }
    }
  colnames(p.mat) <- rownames(p.mat) <- colnames(mat)
  p.mat
}
## matrix of the p-value of the correlation
p.mat <- cor.mtest(mdat.16S_bray[25:41])
head(p.mat)

##color palette
col <- colorRampPalette(c("#053061", "#2166AC", "#4393C3", "#92C5DE", "#D1E5F0", 
                                     "#FFFFFF", "#FDDBC7", "#F4A582", "#D6604D", "#B2182B", 
                                     "#67001F"))(200)

##significant correlation matrix with p-values
corrplot(M, method="color", col=col, order="hclust", 
         addCoef.col = "black", # Add coefficient of correlation
         tl.col="black", tl.srt=45, #Text label color and rotation
         # Combine with significance
         p.mat = p.mat, sig.level = 0.01, insig = "blank",
         number.cex = 0.4,
         # hide correlation coefficient on the principal diagonal
         diag=FALSE 
         )

##all correlations
corrplot(M, method="color", col=col, order="hclust",  # Add coefficient of correlation
         tl.col="black", tl.srt=45, diag=FALSE)
```

##### 3.5 - Figure 4

```
#the imputed dataset have been used to perform phenotypic traits analysis 

#Figure 4
Cap_cluster<- capscale(bray.16S~Heart.formation+Head.height+Head.shape+Leaf.venation+Head.leafs.overlap+Seedling.cotyledon.shape+Leaf.blistering+Side.shoot.formation.tendency+Tipburn.sensitivity+Leaf.division,
                       data = mdat.16S_bray, add=FALSE, subset, na.action = na.omit)

##ANOVA-like permutation analysis
perm_test <- anova(Cap_cluster, permutations = 999)
p_value <- perm_test$Pr[1]
p_value

##Run variability table function
variability_table <- function(cca){

        chi <- c(cca$tot.chi,
                       cca$CCA$tot.chi, cca$CA$tot.chi)
        variability_table <- cbind(chi, chi/chi[1])
        colnames(variability_table) <- c("inertia", "proportion")
        rownames(variability_table) <- c("total", "constrained", "unconstrained")
        return(variability_table)
}


##Generate variability table
var_tbl <- variability_table(Cap_cluster)
ci <- quantile(perm_test$F.perm, c(.05,.95))*perm_test$chi[1]/var_tbl["total", "inertia"]
eig_CCA <- Cap_cluster$CCA$eig
eig_CA <- Cap_cluster$CA$eig
variance <- var_tbl["constrained", "proportion"]
variance_NMDS <- var_tbl["unconstrained", "proportion"]

##Extract unweighted and  unweighted CCA values and unconstrained CA values (here you basically extract the positions of your samples in the ordination space defined by the constrained/unconstrained axes)

###Extract colnames
colnames <- c(paste("u", colnames(Cap_cluster$CCA$u)[1], sep=""), paste("u", colnames(Cap_cluster$CCA$u)[2], sep=""), paste("w", colnames(Cap_cluster$CCA$wa)[1], sep=""), paste("w", colnames(Cap_cluster$CCA$wa)[2], sep=""), paste(colnames(Cap_cluster$CA$u)[1], sep=""), paste(colnames(Cap_cluster$CA$u)[2], sep=""))
  
 ### Extract datapoints 
points = data.frame(c(Cap_cluster$CCA$u[, 1]),
                    c(Cap_cluster$CCA$u[, 2]),
                    c(Cap_cluster$CCA$w[, 1]),
                    c(Cap_cluster$CCA$w[, 2]),
                    c(Cap_cluster$CA$u[, 1]),
                    c(Cap_cluster$CA$u[, 2]))
colnames(points) <- colnames
rownames(points)


##Calculate axis explained variation
eig_tot <- sum(Cap_cluster$CCA$eig)
eigenvalues <- Cap_cluster$CCA$eig/eig_tot
eigenvalues

##Bind with metadata


df <- cbind(points, mdat.16S_bray[match(rownames(points), rownames(mdat.16S_bray)),])


#Extract Arrows

#script for extracting genetic distance arrows
arrow <- data.frame(Cap_cluster$CCA$biplot)

#Plot
pwCPOA <- ggplot(df, aes(x=wCAP1, y=wCAP2)) +
  geom_point(aes(fill=genetic.distance), shape=22, size=3.5, na.value = "grey50") +
  scale_fill_manual(values = colors_var) + 
  labs(
    x = paste("wCPCoA 1 (", format(100 * eig_CCA[1] / sum(eig_CCA), digits=4), "%)", sep=""),
    y = paste("wCPCoA 2 (", format(100 * eig_CCA[2] / sum(eig_CCA), digits=4), "%)", sep="")
  ) +
  ggtitle(paste(format(100 * variance, digits=3), " % of variance; p=", format(p_value, digits=2), sep="")) +
  theme(
    legend.position = "right",
    panel.grid.major = element_line(color = "#EDEDED"),  
    panel.grid.minor = element_line(color = "#EDEDED"),  
    panel.background = element_rect(fill = "white"),   
    plot.background = element_rect(fill = "white")     
  ) +
  geom_segment(data = arrow, aes(x = 0, y = 0, xend = CAP1, yend = CAP2), size = 0.5, 
               arrow = arrow(type = "closed", length = unit(0.2, "inches"))) +
  geom_text(data = arrow, aes(x = CAP1, y = CAP2, label = row.names(arrow)), vjust = -1.5, hjust = NA)


pwCPOA
```

***4 - Alpha diversity analysis*** # 4- Alpha
diversity analysis ### 4.1 - Create a table with Shannon and Observed
indexes

```
#r16S
#merged_phylo_tree
#r16S_raref
#r16S_L

#subset
r16S_L <- subset_samples(merged_phylo_tree, tissue %in% c("leaf"))
r16S_L= subset_samples(r16S_L, genotype2 != "11") #no tree coordinated
r16S_L= subset_samples(r16S_L, genotype2 != "71") #no belonging to genetic group
r16S_L= subset_samples(r16S_L, genotype2 != "119") #no belonging to genetic group
r16S_L= subset_samples(r16S_L, genotype2 != "92") #too much NA values 

library(picante)
#Table of alpha-diversity estimators
table_r16S <- estimate_richness(r16S_L, split = TRUE, measures=c("Observed", "InvSimpson","Shannon"))

#Bind design + table of alpha_div
sdr16S <- sample_data(r16S_L)
datar16S <- cbind(sample_data(sdr16S), table_r16S) 

#add phylogenetic info to table 
rgyrBPD <- as.data.frame.matrix(otu_table(r16S_L))
PD_matrix <- pd(samp = t(rgyrBPD),tree = phy_tree(r16S_L), include.root = F)

#Bind design + table of alpha_div
datar16S <- merge(datar16S, PD_matrix, by="row.names", all=TRUE)
```

##### 4.2 - Figures S3 - Variation in alpha diversity between varieties and groups of closely related genotypes

```
#libraries 
library(ggplot2)
library(tidyverse)
library(ggpubr)
library(rstatix)
library(tidyverse)
library(ggthemes)
library(multcompView)
library(agricolae)
library(dplyr)

#transforma as factor
datar16S$sampleID3 <- as.factor(datar16S$sampleID3)
datar16S$genotype2<-as.factor(datar16S$genetic.distance) #these are groups of closely related genotypes
datar16S$variety<- as.factor(datar16S$variety)
datar16S$block<-as.factor(datar16S$block)

#Figure S3B - variety

#Observed index
#Add Dunn comparisons to the graph
res_obs<-datar16S %>% 
  dunn_test(Observed ~ variety, p.adjust.method = "fdr")

stat.test <- res_obs %>% add_xy_position(x = "variety")


ggplot(datar16S, aes(variety, Observed),  na.rm = FALSE) + 
  geom_boxplot(aes(fill = variety), show.legend = TRUE) +
  geom_jitter(size=1)+
  labs(x="variety", y="Observed ASV richness") +
  scale_fill_manual(values = colors_var)+
  theme_bw() + 
  theme(panel.grid.major = element_blank(), panel.grid.minor = element_blank())+
  stat_pvalue_manual(stat.test, hide.ns = TRUE)->plot_16S_Observed_b

plot_16S_Observed_b

#Shannon index
res_obs<-datar16S %>% 
  dunn_test(Shannon ~ variety, p.adjust.method = "fdr")

stat.test <- res_obs %>% add_xy_position(x = "variety")


ggplot(datar16S, aes(variety, Shannon),  na.rm = FALSE) + 
  geom_boxplot(aes(fill = variety), show.legend = TRUE) +
  geom_jitter(size=1)+
  labs(x="variety", y="Shannon ASV richness") +
  scale_fill_manual(values = colors_var)+
  theme_bw() + 
  theme(panel.grid.major = element_blank(), panel.grid.minor = element_blank())+
  stat_pvalue_manual(stat.test, hide.ns = TRUE)->plot_16S_Shannon_b

plot_16S_Shannon_b

##Figure S3A - genetic.distance

#Observed index
#Add Dunn comparisons to the graph
res_obs<-datar16S %>% 
  dunn_test(Observed ~ genetic.distance, p.adjust.method = "fdr")

stat.test <- res_obs %>% add_xy_position(x = "genetic.distance")


ggplot(datar16S, aes(genetic.distance, Observed),  na.rm = FALSE) + 
  geom_boxplot(aes(fill = genetic.distance), show.legend = TRUE) +
  geom_jitter(size=1)+
  labs(x="genetic.distance", y="Observed ASV richness") +
  scale_fill_manual(values = colors_var)+
  theme_bw() + 
  theme(panel.grid.major = element_blank(), panel.grid.minor = element_blank())+
  stat_pvalue_manual(stat.test, hide.ns = TRUE)->plot_16S_Observed_b

plot_16S_Observed_b

#Shannon index
res_obs<-datar16S %>% 
  dunn_test(Shannon ~ genetic.distance, p.adjust.method = "fdr")

stat.test <- res_obs %>% add_xy_position(x = "genetic.distance")


ggplot(datar16S, aes(genetic.distance, Shannon),  na.rm = FALSE) + 
  geom_boxplot(aes(fill = genetic.distance), show.legend = TRUE) +
  geom_jitter(size=1)+
  labs(x="genetic.distance", y="Shannon ASV richness") +
  scale_fill_manual(values = colors_var)+
  theme_bw() + 
  theme(panel.grid.major = element_blank(), panel.grid.minor = element_blank())+
  stat_pvalue_manual(stat.test, hide.ns = TRUE)->plot_16S_Shannon_b

plot_16S_Shannon_b
```

#### 4.3 - Figures 6 and S6

```
#transform phenotypic traits to factors

datar16S$Seedling.cotyledon.shape<-as.factor(datar16S$Seedling.cotyledon.shape)
datar16S$Head.height<-as.factor(datar16S$Head.height)
datar16S$Seedling.anthocyanin.presence<-as.factor(datar16S$Seedling.anthocyanin.presence)
datar16S$Leaf.shape.cultivated.material<-as.factor(datar16S$Leaf.shape.cultivated.material)
datar16S$Leaf.blistering<-as.factor(datar16S$Leaf.blistering)
datar16S$Leaf.margin.undulation<-as.factor(datar16S$Leaf.margin.undulation)
datar16S$Leaf.venation<-as.factor(datar16S$Leaf.venation)
datar16S$Leaf.division<-as.factor(datar16S$Leaf.division)
datar16S$Leaf.color<-as.factor(datar16S$Leaf.division)
datar16S$Leaf.color.intensity<-as.factor(datar16S$Leaf.color.intensity)
datar16S$Leaf.anthocyanin.content<-as.factor(datar16S$Leaf.anthocyanin.content)
datar16S$Side.shoot.formation.tendency<-as.factor(datar16S$Side.shoot.formation.tendency)
datar16S$Tipburn.sensitivity<-as.factor(datar16S$Tipburn.sensitivity)
datar16S$Plant.diameter<-as.factor(datar16S$Plant.diameter)
datar16S$Head.shape<-as.factor(datar16S$Head.shape)
datar16S$Head.leafs.overlap<-as.factor(datar16S$Head.leafs.overlap)
datar16S$Head.height<-as.factor(datar16S$Head.height)
datar16S$Heart.formation<-as.factor(datar16S$Heart.formation)
datar16S$Observed<-as.numeric(datar16S$Observed)
datar16S$Shannon<-as.numeric(datar16S$Shannon)


#Calculate statistical differences in alpha-diversity inside each phenotypical traits

datar16S %>% 
 pivot_longer(., cols=29:45, names_to= "trait", values_to = "values") -> data_phen

library(ggplot2)

#Shannon index differences
res_Shannon<-data_phen %>% 
  group_by(trait) %>% 
  dunn_test(Shannon ~ values, p.adjust.method = "fdr")
stat.test <- res_Shannon %>% add_xy_position(x = "trait")

#Observed index differences
res_obs<-data_phen %>% 
  group_by(trait) %>% 
  dunn_test(Observed ~ values, p.adjust.method = "fdr")
stat.test <- res_obs %>% add_xy_position(x = "trait")

#Visualize alpha-diversity plot

#Figure 5A- Shannon diversity between heart formation groups

Heart.palette <- c("#ffffe5", "#f7fcb9", "#d9f0a3", "#addd8e", "#78c679", "#41ab5d", "#238443", "#006837", "#004529")

gg_traits <- datar16S %>% 
  ggplot(aes(x = Heart.formation, y = Shannon, fill = Heart.formation)) + 
  geom_boxplot(na.rm = TRUE) +  # Ignora i NA nel boxplot
  geom_jitter(na.rm = TRUE) +
  theme_classic() +
  scale_fill_manual(values = Heart.palette)

gg_traits

#Figure S6A- Observed diversity between heart formation groups
gg_traits <- datar16S %>% 
  ggplot(aes(x = Heart.formation, y = Observed, fill = Heart.formation)) + 
  geom_boxplot(na.rm = TRUE) +  # Ignora i NA nel boxplot
  geom_jitter(na.rm = TRUE) +
  theme_classic() +
  scale_fill_manual(values = Heart.palette)
gg_traits

#Figure 5B- Shannon diversity between Head height groups

Height.palette <- c("#ffffd4", "#fee391", "#fec44f", "#fe9929", "#d95f0e", "#993404")


gg_traits <- datar16S %>% 
  ggplot(aes(x = Head.height, y = Shannon, fill = Head.height)) + 
  geom_boxplot(na.rm = TRUE) +  
  geom_jitter(na.rm = TRUE) +
  theme_classic() +
  scale_fill_manual(values = Height.palette)

#Figure S4B- Observed diversity between head height groups
gg_traits <- datar16S %>% 
  ggplot(aes(x = Head.height, y = Observed, fill = Head.height)) + 
  geom_boxplot(na.rm = TRUE) +  
  geom_jitter(na.rm = TRUE) +
  theme_classic() +
  scale_fill_manual(values = Height.palette)

#Figure 5C- Shannon diversity between Leaf venation groups

#palette
colors <- c("#d8b36580", "#5ab4ac80")
dark_colors <- c("#d8b365", "#5ab4ac")

# Violin plot leaf venation Shannon
gg_traits <- datar16S %>% 
  ggplot(aes(x = Leaf.venation, y = Shannon)) + 
  geom_violin(aes(fill = Leaf.venation), na.rm = TRUE, alpha = 0.2) +  
  geom_boxplot(aes(fill = Leaf.venation), width = 0.1, na.rm = TRUE, fill = "white", color = "black") +  
  geom_jitter(aes(fill = Leaf.venation, color = Leaf.venation), width = 0.2, size = 3, shape = 21, stroke = 1.5, na.rm = TRUE, alpha = 0.6) +  
  theme_classic() +
  scale_fill_manual(values = colors) +
  scale_color_manual(values = dark_colors) +  
  guides(fill = FALSE) 
gg_traits

#Figure S6C- Observed diversity between head shape groups

#palette
Head.shape.palette <- c("#f1eef6", "#d4b9da", "#c994c7", "#df65b0", "#dd1c77")


gg_traits <- datar16S %>% 
  ggplot(aes(x = Head.shape, y = Observed, fill = Head.shape)) + 
  geom_boxplot(na.rm = TRUE) +
  geom_jitter(na.rm = TRUE) +
  theme_classic() +
  scale_fill_manual(values = Head.shape.palette)
```

##### 4.4 - Figures 6A-B and S6A-B- Spearman correlations between phenotypic traits and alpha- diversity indexes

```
library(ggplot2)
library(Kendall)

#transform to factor
datar16S$Head.height<-as.numeric(datar16S$Head.height)
datar16S$Heart.formation<-as.numeric(datar16S$Heart.formation)

#Figure 6A- spearman's Rho correlation coefficient for heart formation and Shannon index

res<-cor.test(datar16S$Heart.formation, datar16S$Shannon, method ="spearman")
print(res) 

#scatter plot
ggplot(datar16S, aes(x = Heart.formation, y = Shannon)) +
  geom_point() +  # Color of the points
  geom_smooth(method = "lm", color = "#006837", fill = "#addd8e") 
  labs(x = "Heart formation", y = "Shannon index") +
  theme_minimal()

#Figure S6A- spearman's Rho correlation coefficient for heart formation and Observed index
  
res<-cor.test(datar16S$Heart.formation, datar16S$Observed, method ="spearman")
print(res) 
  
#scatter plot
ggplot(datar16S, aes(x = Heart.formation, y = Observed)) +
  geom_point() +  # Color of the points
  geom_smooth(method = "lm", color = "#006837", fill = "#addd8e") + 
  labs(x = "Heart formation", y = "Observed index") +
  theme_minimal()

#Figure 5B- spearman's Rho correlation coefficient for Head height and Shannon index

res<-cor.test(datar16S$Head.height, datar16S$Shannon, method ="spearman")
print(res) 

#scatter plot
ggplot(datar16S, aes(x = Head.height, y = Shannon)) +
  geom_point() +
  geom_smooth(method = "lm", color = "#d95f0e", fill = "#fee391")
  labs(x = "Heart formation", y = "Shannon index") +
  theme_minimal()

#Figure S6B- spearman's Rho correlation coefficient for Head height and Observed index

res<-cor.test(datar16S$Head.height, datar16S$Observed, method ="spearman")
print(res) 

#scatter plot
ggplot(datar16S, aes(x = Head.height, y = Observed)) +
  geom_point() + 
  geom_smooth(method = "lm", color = "#d95f0e", fill = "#fee391")+
  labs(x = "Heart formation", y = "Observed index") +
  theme_minimal()
```

##### 4.5 - Figure S8

```
#Custom X-axis samples so that they are ordered according to the sampling order

#Function
custom_order <- function(x) {
  num <- as.numeric(str_extract(x, "\\d+"))
  letter <- str_extract(x, "[A-Za-z]")
  temp_df <- data.frame(sampleID3 = x, num = num, letter = letter)
  temp_df <- temp_df %>% arrange(letter, num)
  temp_df$new_index <- 1:nrow(temp_df)
  return(temp_df)
}

#Apply function and order samples (sampleID3)
ordered_data <- custom_order(datar16S$sampleID3)
datar16S <- datar16S %>%
  left_join(ordered_data, by = c("sampleID3" = "sampleID3"))

#Calculate the Spearman coefficient 
res <- cor.test(datar16S$new_index, datar16S$Shannon, method = "spearman")
print(res)

#color palette
block_colors <- c(
  "A" = "#fde0dd",
  "B" = "#fa9fb5",
  "C" = "#c51b8a"
)

#Figure S8A- Scatter plot

gg_traits <- ggplot(datar16S, aes(x = new_index, y = Shannon, color = block)) +
  geom_point(size=6) +  
  geom_smooth(method = "lm", color = "#5B3A29", fill = "#BFAE9A", se = TRUE) +
  labs(x = "Sampling order", y = "Shannon index") +
  scale_x_continuous(breaks = ordered_data$new_index, labels = ordered_data$sampleID3) + 
  scale_color_manual(values = block_colors) + 
  theme_minimal() +
  theme(
    axis.text.x = element_text(angle = 90, hjust = 1, vjust = 0.5, size = 5), 
    panel.grid.major = element_blank(),
    panel.grid.minor = element_blank(),
    panel.border = element_blank() 
  )

#Figure S8B - Box plot

gg_traits <- ggplot(datar16S, aes(x = new_index, y = Shannon, color = block)) +
  geom_boxplot() +
  geom_point(size = 4) +  # Points colored by block parameter
  labs(x = "Sampling order", y = "Shannon index") +
  scale_x_continuous(breaks = ordered_data$new_index, labels = ordered_data$sampleID3) +  # Maintain original labels
  scale_color_manual(values = block_colors) +  # Assign colors to blocks
  theme_minimal() +
  theme(
    axis.text.x = element_text(angle = 90, hjust = 1, vjust = 0.5, size = 5),  # Rotate x-axis text
    panel.grid.major = element_blank(),  
    panel.grid.minor = element_blank(),  
    panel.border = element_blank() 
  )

gg_traits

res_obs<-datar16S %>% 
  dunn_test(Shannon ~ block, p.adjust.method = "fdr")

#Figure S8C 

#Shannon by blocks
gg_traits <- datar16S %>% 
  ggplot(aes(x = Heart.formation, y = Shannon)) + #change Heart.formation with Head.height with height or Leaf.venation
  geom_boxplot() +  
  geom_jitter(aes(color = block), size = 3, na.rm = TRUE) +  # Points colored by block
  theme_classic() +
  scale_color_manual(values = block_colors)
```

##### 4.6 - Figure S5A

```
#bind micro/macro nutrients' coordinates to alpha-diversity table
datar16S$sampleID  <- rownames(datar16S)
rownames(datar16S) <- NULL
datar16S <- cbind(sampleID = datar16S$sampleID, datar16S[, -ncol(datar16S)])
datar16S$sample_ID <- NULL
sampleID3_values <- rownames(datar16S)
var <- var %>% mutate(sampleID3 = rownames(var))
datar16S <- left_join(datar16S, var, by = "sampleID3")
rownames(datar16S) <- datar16S[,1]

#Calculate Spearman coefficient between Shannon index and dimensions
#Example for Dim.1
res<-cor.test(datar16S$Dim.1, datar16S$Shannon, method ="spearman")
res

## scatter plot 
ggplot(datar16S, aes(x = Dim.1, y = Shannon)) +
  geom_point() +
  geom_smooth(method = "lm", color = "#2c7fb8", fill = "#7fcdbb") + 
  labs(x = "Dim.1", y = "Observed index") +
  theme_minimal()

#Calculate Spearman coefficient between Observed index and dimensions
#Example for Dim.1
res<-cor.test(datar16S$Dim.1, datar16S$Observed, method ="spearman")
res

## scatter plot 
ggplot(datar16S, aes(x = Dim.1, y = Observed)) +
  geom_point() +
  geom_smooth(method = "lm", color = "#2c7fb8", fill = "#7fcdbb") + 
  labs(x = "Dim.1", y = "Observed index") +
  theme_minimal()
```

***5-FAPROTAX*** # 5- FAPROTAX analysis to
uncover association with mammal and human gut-associated bacteria:
Figure S7A

```
#libraries
library(phyloseq)
library(microeco)
library("file2meco")

###########First part: prepare the dataset

#Transform the phyloseq object into microtable object
meco_dataset <- phyloseq2meco(r16S_L)

#prepare tables
t2 <- trans_func$new(meco_dataset)
t2$cal_spe_func(prok_database = "FAPROTAX")
t2$res_spe_func
t2$cal_spe_func_perc(abundance_weighted = FALSE)
t2$trans_spe_func_perc()
t2$plot_spe_func_perc()

#We want to know the number of OTU assigned to each function
func_table0 <- t2$res_spe_func

#Number of OTUs without any associated function in FAPROTAX
#and number of unensured functions
length(which(rowSums(func_table0) == 0)) 
length(which(colSums(func_table0) == 0)) 
func_table0 <- func_table0[-which(rowSums(func_table0) == 0), -which(colSums(func_table0) == 0)]
write.csv(func_table0, file = "Bacterial_functional_annotation_FAPROTAX_for_16S_zOTUs.csv")

length(which(rowSums(func_table0) == 0)) 
colSums(func_table0)
ASV_table<- as.data.frame(otu_table(r16S_L))

#Calculate the percentage of OTUs within a community that ensure the same function
#First, list the OTUs that are present in each site
list_sites_otu <- apply(ASV_table, 2, function(x) rownames(ASV_table)[which(x>0)])
#Create an empty table with functions in columns and sites in rows 
df_func_site <- as.data.frame(matrix(0, ncol = ncol(func_table0), nrow = ncol(ASV_table), dimnames = list(colnames(ASV_table), colnames(func_table0))))
#For each function and each site, find the number of OTUs present in the site
#that ensure the given function 
for(i in 1:ncol(df_func_site)){
  for(j in 1:nrow(df_func_site)){
    df_ind_prim <- func_table0[which(func_table0[, colnames(df_func_site)[i]] >0),]
    df_ind1_prim <- df_ind_prim[which(rownames(df_ind_prim) %in% list_sites_otu[[j]]),]
    df_func_site[j,i] <- sum(ASV_table[rownames(ASV_table) %in% rownames(df_ind1_prim), j])/colSums(ASV_table)[1]
  }
}
func_table_perc_16S <- df_func_site

save(func_table_perc_16S, file = "Bacterial_functional_groups_16S_zOTUs.RData")

write.table(func_table_perc_16S, "/yourPath/new_func_table_perc.txt", sep=",")

write.csv(func_table_perc_16S,'new_func_table_perc.csv')

#############end of the first part################

#part 2- statistical analysis

library(car)
library(ggplot2)
library(ggpubr)
library(rstatix)
library(psych)

#import tables
functab <- read.table("FAPROTAX/new_func_table_perc.csv", 
                       sep = ";",
                       dec = ".",
                       header = TRUE) 

#transform to factor
functab$sampleID3<-as.factor(functab$sampleID3)
functab$genetic.distance<-as.factor(functab$genetic.distance)
functab$variety<-as.factor(functab$variety)
functab$block<-as.factor(functab$block)
functab$Seedling.cotyledon.shape<-as.factor(functab$Seedling.cotyledon.shape)
functab$Seedling.anthocyanin.presence<-as.factor(functab$Seedling.anthocyanin.presence)
functab$Leaf.shape.cultivated.material<-as.factor(functab$Leaf.shape.cultivated.material)
functab$Leaf.blistering<-as.factor(functab$Leaf.blistering)
functab$Leaf.margin.undulation<-as.factor(functab$Leaf.margin.undulation)
functab$Leaf.venation<-as.factor(functab$Leaf.venation)
functab$Leaf.division<-as.factor(functab$Leaf.division)
functab$Leaf.color<-as.factor(functab$Leaf.division)
functab$Leaf.color.intensity<-as.factor(functab$Leaf.color.intensity)
functab$Leaf.anthocyanin.content<-as.factor(functab$Leaf.anthocyanin.content)
functab$Side.shoot.formation.tendency<-as.factor(functab$Side.shoot.formation.tendency)
functab$Tipburn.sensitivity<-as.factor(functab$Tipburn.sensitivity)
functab$Plant.diameter<-as.factor(functab$Plant.diameter)
functab$Head.shape<-as.factor(functab$Head.shape)
functab$Head.leafs.overlap<-as.factor(functab$Head.leafs.overlap)
functab$Head.height<-as.factor(functab$Head.height)
functab$Heart.formation<-as.factor(functab$Heart.formation)

#before doing dunn post hoc test check for normality to see if you can apply ANOVA. Our data were not normally distributed. 

#statistical tests
dunn_test(mammal_gut ~ Leaf.venation, data = functab) 
#substitute mammal_gut with human_gut associated bacteria or human associated bacteria (or any other variable in the dataset)
#results are reported in Figure S6B

#################end of the second part###############

#part 3: graphical representation

#Figure S7A

#mammal gut associated bacteria

gg_traits <- functab_filtered %>% 
  ggplot(aes(x = Leaf.venation, y = mammal_gut)) + 
  geom_violin(aes(fill = Leaf.venation), na.rm = TRUE, alpha = 0.2) +
  geom_boxplot(aes(fill = Leaf.venation), width = 0.1, na.rm = TRUE, fill = "white", color = "black") +  
  geom_jitter(aes(fill = Leaf.venation, color = Leaf.venation), width = 0.2, size = 3, shape = 21, stroke = 1.5, na.rm = TRUE, alpha = 0.6) + 
  theme_classic() +
  scale_fill_manual(values = colors) +
  scale_color_manual(values = dark_colors) + 
  guides(fill = FALSE) 
gg_traits

#Human gut associated bacteria

gg_traits <- functab%>% 
  ggplot(aes(x = Leaf.venation, y = human_gut)) + 
  geom_violin(aes(fill = Leaf.venation), na.rm = TRUE, alpha = 0.2) + 
  geom_boxplot(aes(fill = Leaf.venation), width = 0.1, na.rm = TRUE, fill = "white", color = "black") + 
  geom_jitter(aes(fill = Leaf.venation, color = Leaf.venation), width = 0.2, size = 3, shape = 21, stroke = 1.5, na.rm = TRUE, alpha = 0.6) +  
  theme_classic() +
  scale_fill_manual(values = colors) +
  scale_color_manual(values = dark_colors) + 
  guides(fill = FALSE) 
gg_traits

#Human associated bacteria

gg_traits <- functab%>% 
  ggplot(aes(x = Leaf.venation, y = human_associated)) + 
  geom_violin(aes(fill = Leaf.venation), na.rm = TRUE, alpha = 0.2) + 
  geom_boxplot(aes(fill = Leaf.venation), width = 0.1, na.rm = TRUE, fill = "white", color = "black") +  
  geom_jitter(aes(fill = Leaf.venation, color = Leaf.venation), width = 0.2, size = 3, shape = 21, stroke = 1.5, na.rm = TRUE, alpha = 0.6) + 
  theme_classic() +
  scale_fill_manual(values = colors) +
  scale_color_manual(values = dark_colors) + 
  guides(fill = FALSE) 
gg_traits
```

***6-Log2Fold change analysis***

### 6- Log2Fold change analysis with Dseq2: Figures 7A and S9A

```
##part 1: prepare the dataset with phenotypic traits in binary form

#heart formation: groups 1-2 are considered "without the heart formation capability", group 7 is considered capable of forming a tight heart
sample_data(r16S_L)$Heart.formation_binary <- ifelse(
  is.na(sample_data(r16S_L)$Heart.formation), 
  NA,
  ifelse(sample_data(r16S_L)$Heart.formation %in% c("1", "2"), "absent", 
         ifelse(sample_data(r16S_L)$Heart.formation %in% c("7"), "present", NA))
)

#head height: plants in group 3 are considered short, in group 5 tall.
sample_data(r16S_L)$Head.height_binary <- ifelse(
  is.na(sample_data(r16S_L)$Head.height), # Check if Head.height is NA
  NA,
  ifelse(sample_data(r16S_L)$Head.height == "3", "short",
         ifelse(sample_data(r16S_L)$Head.height %in% c("5"), "high", NA))
)

#Leaf venation is already binary

##part 2: apply the Dseq2 function

library( "DESeq2" )
library(ggplot2)
library(tidyverse)
library(ggrepel)

#in the following script substitute "Heart.formation_binary" with the phenotypic trait you want to study

#select the data in the binary format
r16S_L_selected= subset_samples(r16S_L, Heart.formation_binary != "NA")

#tranform as factor
sample_data(r16S_L_selected)$Heart.formation_binary<-as.factor(sample_data(r16S_L_selected)$Heart.formation_binary)

#apply sizeFactors(diagdds) function
diagdds = phyloseq_to_deseq2(r16S_L_selected, ~Heart.formation_binary)

diagdds = DESeq(diagdds, test="Wald", fitType="parametric")

res = results(diagdds, cooksCutoff = FALSE)

#to see the levels' order in the graph
head(res)

#set the p-value
alpha = 0.05
sigtab = res[which(res$pvalue < alpha), ]
sigtab = cbind(as(sigtab, "data.frame"), as(tax_table(r16S_L)[rownames(sigtab), ], "matrix"))

dimnames
head(sigtab)
dim(sigtab)

##part 3: graphical representation. Figures 6A and S8A

t #set the theme
theme_set(theme_bw() + 
          theme(panel.grid.major = element_blank(),
                panel.grid.minor = element_blank(),
                panel.border = element_rect(color = "black", fill = NA),
                axis.title.x = element_text(size = 12),
                axis.title.y = element_text(size = 12),
                axis.text.x = element_text(size = 7),
                axis.text.y = element_text(size = 7),
                legend.title = element_text(size = 10), 
                legend.text = element_text(size = 8)))

# Reorder the Log2Foldchange values in the plot
sigtab <- sigtab[order(sigtab$log2FoldChange, decreasing = FALSE), ]
sigtab$ASV <- factor(sigtab$ASV, levels = sigtab$ASV)  

#
gg <- ggplot(sigtab, aes(x = log2FoldChange, y = ASV, color = Order, size = baseMean)) +
  geom_point() +
  geom_vline(xintercept = 0, linetype = "dashed", color = "black") +
  scale_size_continuous(range = c(1, 10)) + 
  scale_color_brewer(palette = "Set1") +   
  labs(x = "log2 Fold Change", y = "Taxa", size = "Base Mean", color = "Order") +
  theme(axis.text.x = element_text(angle = 90, vjust = 0.5, hjust = 1, size = 7))
```

***7-Network analysis*** # 7- Network analysis:
Figures 7B-C and S9B-C

```
# Libraries
library(phyloseq)
library(tidyverse)
library(igraph)

#In the following script, substitute "Heart.formation_binary" with the other phenotypic traits you want to study, and "present" with the other phenotypic forms that were identified in section 6 part1

r16S_L_present= subset_samples(r16S_L, Heart.formation_binary == "present")
view(sample_data(r16S_L_present))

# Extract tax_table from phyloseq object
taxa_table <- as.data.frame(tax_table(r16S_L_present))
taxa_table <- tibble::rownames_to_column(taxa_table, var = "species")

# Extract metadata table from phyloseq object
metadata <- sample_data(r16S_L_present)
metadata <- as.data.frame(metadata)
metadata$rownames <- rownames(metadata)
metadata <- as_tibble(metadata) %>%
  column_to_rownames(var = "rownames")

# Extract OTU table from phyloseq object and convert to relative abundance
otu_table <- as.data.frame(otu_table(r16S_L_present))

# Convert OTU table to relative abundance
sp_ratio <- otu_table %>%
  mutate(across(everything(), ~ ./sum(.))) %>%
  rownames_to_column(var = "species") %>% 
  as_tibble()

# Display the first few rows of the sp_ratio
head(sp_ratio)

# Transform sp_ratio for network analysis
sp_ratio <- sp_ratio %>% 
  as_tibble() %>% 
  rename_all(tolower) %>% 
  mutate(species = str_replace_all(species, "__", "")) %>% 
  column_to_rownames(var = "species")

min.prevalence= 5
#for heart formation present: min.prevalence set at 6
#for heart formation absent: min.prevalence set at 6
#for head height short: min.prevalence set at 10
#for head height high: min.prevalence set at 4
#for leaf venation 1 : min.prevalence set at 12
#for leaf venation 2 : min.prevalence set at 5
incidence=sp_ratio
incidence[incidence>0]=1
sp_ratio_filtered <- sp_ratio[which(rowSums(incidence)>=min.prevalence),] ### end of prevalence filtering

sp_correl <- sp_ratio_filtered %>% 
  t() %>% 
  cor(method = "spearman")  ### correlation calculations

sp_correl[abs(sp_correl)<0.65]=0  ### define threshold for correlations

net_work <- graph_from_adjacency_matrix(sp_correl,mode="lower",weighted=TRUE, diag=FALSE)  ## create network file
net_work <- delete.vertices(net_work, degree(net_work)==0) #remove nodes without edges

plot(net_work, vertex.label = NA, edge.width = 5, vertex.size=10) ## first plot

###annotations

taxa_tbl <- taxa_table %>%
  as_tibble() %>%
  mutate(species = as.character(species), 
         species = ifelse(is.na(species), "unknown", species),
         species = str_replace_all(species, "__", ""))  #tidy taxonomic infos

net_work_used <- V(net_work)$name %>% 
   as_tibble() %>% 
  mutate(species = value) %>% 
  select(species) #extract species represented in the network

v_attr <- sp_ratio %>% ### we create a table of attributes for nodes (vertex)
  rownames_to_column( var = "species") %>% 
  as_tibble() %>% 
  pivot_longer(-species, names_to = "sample_id", values_to = "ratio" ) %>% 
  mutate(species = str_replace_all(species, "__", "")) %>% 
  group_by(species) %>% 
  summarise(rel_abundance = sum(ratio)) %>% 
  inner_join(net_work_used, by = "species") %>% 
  inner_join(taxa_tbl, by = "species") %>%  
  mutate(rel_abundance = abs(exp (rel_abundance))) # we join taxonomic infos, relative abundance and species that were only represented in the network to create the attribute table

  
network_table <- igraph::as_data_frame(net_work, 'both') ##we convert the network in data frames
     
network_table$vertices <- network_table$vertices %>%
    as_tibble() %>% 
    inner_join(v_attr, by = c("name"="species"))  # we add our attribute in the data frames 
  
  net_work1 <- graph_from_data_frame(network_table$edges,
                                     directed = F,
                                     vertices = network_table$vertices) # we

################plot
  
mom_data <- V(net_work1)$Order %>% 
  as_tibble_col(column_name = "species") #formating the class variable as factor (is needed for coloring edges)
mom_data$species <- as_factor(mom_data$species)

color_easy <-  c("#FF0000", "#000080", "#32CD32", "#FFD700","#00FFFF", "#800080",  "#FF00FF", "#4682B4", "#98FB98", "#FF7F50", "#FF1493", "#00BFFF", "#8B0000", "#228B22", "#B22222", "#0047AB",  "#808000", "#FFFF00", "#CC5500", "#4B0082",  "#00CED1", "#D8BFD8", "#4169E1", "#7CFC00",  "#8B4513", "#2F4F4F", "#FFFACD", "#20B2AA",  "#C71585", "gray" ,"#FFA500")[mom_data$species] #creating a color palette to represent each class levels

V(net_work1)$color <-  color_easy ## we have now the color attributes based on class

E(net_work1)$sign <- E(net_work1)$weight ##we create another attribute from the weight

E(net_work1)$weight <- abs(E(net_work1)$weight) ## we then use absolute value of weight because of the specific layout we will use

######network plot: Figure 6B and S9B-C

plot(net_work1, 
     vertex.size = 10, 
     edge.width = abs(E(net_work1)$weight) * 2, 
     vertex.label= NA, 
     edge.color = ifelse(E(net_work1)$sign > 0, "blue", "red"),
     vertex.color = (net_work1)$Order, 
     layout = layout_with_kk(net_work1)) 

title_legend1 <- V(net_work1)$Order %>% 
  as_factor() %>% 
  levels() 

legend(x = 1, y = 1, title_legend1,
       pch = 21, pt.bg = c("#FF0000", "#000080", "#32CD32", "#FFD700","#00FFFF", "#800080",  "#FF00FF", "#4682B4", "#98FB98", "#FF7F50", "#FF1493", "#00BFFF", "#8B0000", "#228B22", "#B22222", "#0047AB",  "#808000", "#FFFF00", "#CC5500", "#4B0082",  "#00CED1", "#D8BFD8", "#4169E1", "#7CFC00",  "#8B4513", "#2F4F4F", "#FFFACD", "#20B2AA",  "#C71585", "gray" ,"#FFA500"), 
       pt.cex = 1.5, bty = "n", ncol = 1)


######hub ASVs plot: Figures 6C and S9B-C

#identify the hub score
hs <- hub_score(net_work1)$vector

#plot
plot(net_work1, 
     vertex.size = hs*20, 
     edge.width = abs(E(net_work1)$weight) * 2, 
     vertex.label.cex= 0.6, 
     edge.color = ifelse(E(net_work1)$sign > 0, "blue", "red"),
     vertex.color = (net_work1)$Order, 
     layout = layout_with_kk(net_work1)) 

title_legend1 <- V(net_work1)$Order %>% 
  as_factor() %>% 
  levels() 

legend(x = 1, y = 1, title_legend1,
       pch = 21, pt.bg = c("#FF0000", "#000080", "#32CD32", "#FFD700","#00FFFF", "#800080",  "#FF00FF", "#4682B4", "#98FB98", "#FF7F50", "#FF1493", "#00BFFF", "#8B0000", "#228B22", "#B22222", "#0047AB",  "#808000", "#FFFF00", "#CC5500", "#4B0082",  "#00CED1", "#D8BFD8", "#4169E1", "#7CFC00",  "#8B4513", "#2F4F4F", "#FFFACD", "#20B2AA",  "#C71585", "gray" ,"#FFA500"), 
       pt.cex = 1.5, bty = "n", ncol = 1)
```

***8-Varpart analysis*** # 8- Varpart analysis:
Figure 5 and Tables S12-13

```
#library
library(vegan)

# Table S12- Create the model
mod <- varpart(bray.16S, ~genetic.distance, ~ variety,~ Dim.2+Dim.4+Dim.5,~ Heart.formation+Head.height+Head.shape+Leaf.venation+Head.leafs.overlap+Seedling.cotyledon.shape+Leaf.blistering+Side.shoot.formation.tendency+Tipburn.sensitivity+Leaf.division,
    data=mdat.16S_bray, transfo="hel")
mod
summary(mod)


#Table S13 - Redundancy analysis (RDA) model using Hellinger-transformed variables

#genetic distance
rda.result <- rda(decostand(bray.16S, "hell") ~ genetic.distance, data = mdat.16S_bray)
anova(rda.result)
#variety
rda.result <- rda(decostand(bray.16S, "hell") ~ variety, data = mdat.16S_bray)
anova(rda.result)
#mineral content
rda.result <- rda(decostand(bray.16S, "hell") ~ Dim.2+Dim.4+Dim.5, data = mdat.16S_bray)
anova(rda.result)
#phenotypic traits
rda.result <- rda(decostand(bray.16S, "hell") ~ Heart.formation+Head.height+Head.shape+Leaf.venation+Head.leafs.overlap+Seedling.cotyledon.shape+Leaf.blistering+Side.shoot.formation.tendency+Tipburn.sensitivity+Leaf.division,
    data=mdat.16S_bray)
anova(rda.result)

#Figure 5A- UpSetR plot

library(UpSetR)

input <- c(
  "Genetic distance" = 3.5,
  "Variety" = 4.5,
  "Mineral content" = 2.4,
 "Phenotypic traits" = 6.5,
  "Variety&Genetic distance" = 7,
  "Genetic distance&Mineral content" = 5.8,
  "Genetic distance&Phenotypic traits" = 8.7,
  "Variety&Mineral content" = 6.8,
  "Variety&Phenotypic traits" = 9.6,
  "Mineral content&Phenotypic traits" = 8.6,
  "Mineral content&Genetic distance&Variety" = 9.2,
  "Genetic distance&Variety&Phenotypic traits" = 11.8,
  "Genetic distance&Mineral content&Phenotypic traits" = 10.9,
 "Variety&Mineral content&Phenotypic traits"= 11.7,
 "Genetic distance&Variety&Mineral content&Phenotypic traits"=13.9)


names_list <- sub("=.*", "", names(input))
numeric_values <- as.numeric(input)

# Combine names and values
sorted_data <- data.frame(names = names_list, values = numeric_values)
sorted_data <- sorted_data[order(-sorted_data$values), ]

# Create a bar plot using ggplot2
ggplot(sorted_data, aes(x = factor(names, levels = sorted_data$names), y = values)) +
  geom_bar(stat = "identity", fill = "gray23", width = 0.7) +
  labs(title = "Total variance explained by each factor", x = "Category", y = "% of variation") +
  theme(axis.text.x = element_text(angle = 45, hjust = 1), 
        panel.border = element_blank(),  # Remove border around the image
        panel.grid.major = element_blank(),  # Remove major grid lines
        panel.grid.minor = element_blank(),  # Remove minor grid lines
        panel.background = element_blank(),  # Remove gray background
        plot.background = element_blank(),   # Remove plot background
        axis.line = element_line(color = "black")) + 
  geom_text(aes(label = values), vjust = -0.5, size = 3)

library(UpSetR)
set_order <- c("Phenotypic traits", "Variety", "Genetic distance", "Mineralcontent")
                
upset<-upset(fromExpression(input), 
      nintersects = 15, 
      nsets = 15, 
      order.by = "freq", 
      decreasing = T, 
      mb.ratio = c(0.6, 0.4),
      number.angles = 0, 
      text.scale = 1.5, 
      point.size = 2.8, 
      line.size = 1
      )
upset
```

***9-HEATMAP*** # 9- Figure S10 -heatmap

```
#Creation of an heatmap with differential relative abundance between the different heart formation stages

gpt <- subset_taxa(r16S_L, Kingdom == "d__Bacteria")

# Etract OTU matrix and taxonomical data 
otu_table_df <- as.data.frame(otu_table(gpt))
tax_table_df <- as.data.frame(tax_table(gpt))

#Be sure rownames are the same 
otu_table_df$ASV <- rownames(otu_table_df)
tax_table_df$ASV <- rownames(tax_table_df)

# join OTU table with taxa table 
otu_taxa_df <- merge(otu_table_df, tax_table_df, by = "ASV")
otu_taxa_df <- otu_taxa_df[, -c(224:226)] #delete eventual columns like "confidence", "interval", "kingdom"

# Convert data in long format
library(tidyr)
#as character
otu_taxa_df[] <- lapply(otu_taxa_df, function(x) if(is.numeric(x)) as.character(x) else x)

otu_table_long <- pivot_longer(
  otu_taxa_df, 
  cols = -c(ASV, Phylum, Class, Order, Family, Genus, Species), 
  names_to = "SampleID", 
  values_to = "Abundance"
)

# Add SampleID colum in metadata 
metadata_df <- as.data.frame(sample_data(gpt))
metadata_df$SampleID <- rownames(metadata_df)

# Debugging: check if it works
print(names(metadata_df))
head(metadata_df)

# add metadata to the long-format table
add_metadata <- function(otu_table_long, metadata) {
  metadata_df <- metadata
  metadata_df <- metadata_df[, c("SampleID", "Heart.formation")]
  
  # function to add metadata
  otu_table_long$Heart.formation <- NA
  for (i in seq_len(nrow(otu_table_long))) {
    sample_id <- otu_table_long$SampleID[i]
    if (sample_id %in% metadata_df$SampleID) {
      otu_table_long$Heart.formation[i] <- metadata_df$Heart.formation[metadata_df$SampleID == sample_id]
    }
  }
  
  return(otu_table_long)
}


# apply function
otu_table_long <- add_metadata(otu_table_long, metadata_df)
otu_table_long$Family<-as.factor(otu_table_long$Family)
otu_table_long$Heart.formation<-as.factor(otu_table_long$Heart.formation)
otu_table_long$Abundance<-as.numeric(otu_table_long$Abundance)

otu_table_long %>% 
  filter(!is.na(Heart.formation)) %>% 
  droplevels()

#Select only relevant columns
otu_table_long_1 <- otu_table_long[, c(5,9, 10)]

otu_table_long_1 %>%
  group_by(Family, Heart.formation) %>%
  summarise (Abundance= mean(Abundance)) -> otu_table_long_1

#Transform to wide
otu_table_long_1_wide <- otu_table_long_1 %>%
  pivot_wider(
    names_from = Heart.formation,
    values_from = Abundance,
    names_vary = "slowest"
  )

#Convert to data frame
otu_table_long_1_wide <- as.data.frame(otu_table_long_1_wide)

#Remove NA in ASV column 
otu_table_long_1_wide <- na.omit(otu_table_long_1_wide)

#Set rownames
row.names(otu_table_long_1_wide) <- otu_table_long_1_wide[, 1]

#Remove col1
otu_table_long_1_wide <- otu_table_long_1_wide[, -1]

#Convert to matrix
otu_table_long_1_wide_matrix <- as.matrix(otu_table_long_1_wide)

#Quantile breakes function
quantile_breaks <- function(xs, n = 10) {
  xs <- na.omit(xs)
  zero_included <- 0
  positive_values <- xs[xs > 0]
  quantiles <- quantile(positive_values, probs = seq(0, 1, length.out = n - 1), na.rm = TRUE)
  breaks <- unique(c(zero_included, quantiles))
  breaks <- sort(breaks)
  return(breaks)
}
mat_breaks <- quantile_breaks(otu_table_long_1_wide_matrix, n = 10)


#pheatmap: Figure S9
pheatmap(
  mat               = otu_table_long_1_wide_matrix,
  color             = colorRampPalette(c("#f7fcf0", "#e0f3db","#ccebc5", "#a8ddb5", "#7bccc4", "#4eb3d3","#2b8cbe","#0868ac", "#084081"   ))(10),
 breaks            = mat_breaks,
  border_color      = NA,
  show_colnames     = TRUE,
  show_rownames     = TRUE,
  cluster_cols = F,
 cellwidth         = 30, 
 # annotation_col    = mat_col,
  #annotation_colors = mat_colors,
  drop_levels       = TRUE,
  fontsize          = 10,
 fontsize_row      = 2,    
  fontsize_col      = 10, 
 cutree_rows = 2,
  main              = "Family relative abundance across heart formation capacities"
)
```

***10-FEAST***

### 10- FEAST analyis, unravelling origins: Figure S2

##### 10.1- Dataset preparation

```
#To do this part we used the 1) leaf dataset with 2) no rarefaction and 3) no Abundance threshold. 

Denoising <- read.csv("Output_qiime_qzv/Denoising.tsv", header=TRUE, sep = "\t",  dec=".")

Denoising

#Download packages

library("phyloseq")
library("ggplot2")      
library("readxl")       
library("dplyr")        
library("tibble")
library("tidyverse")
library("magrittr")


#Upload tables
##ASV Table
asv16S <- read.table(file="yourPath/16S_table-FINAL.txt",sep="\t",dec = ".",header=TRUE) 
##Tax Table
tax16S <- read.table(file="yourPath/taxonomy.tsv",sep="\t",dec = ".",header=TRUE)
##Design
  design <- read.table(file="yourPath/Design.txt",sep="\t",dec = ",",header=TRUE)

##Sequences
library(Biostrings)
refseq16S <- readDNAStringSet("yourPath/dna-sequences.fasta")

#Manage data for Phyloseq
asv16S <- asv16S %>%
    tibble::column_to_rownames("OTU.ID") 
tax16S <- tax16S %>% 
    tibble::column_to_rownames("OTU.ID")

#Collapse Tax Table
library(stringr)
tax16S[c("Kingdom","Phylum", "Class", "Order", "Family", "Genus", "Species")] <- str_split_fixed(tax16S$Taxon, ';', 7)
tax16S[tax16S == ""] <- NA 
tax16S[tax16S == " "] <- NA  
design <- design %>% 
    tibble::column_to_rownames("sampleID") 

otu_mat <- as.matrix(asv16S)
tax_mat <- as.matrix(tax16S)

#Create Phyloseq Object  #phylogenetic info to be added
ps16S_0 <- phyloseq(tax_table(tax_mat),
                  otu_table(otu_mat, taxa_are_rows = TRUE), sample_data(design), refseq(refseq16S))

#TAXA
# Calculate the number of taxa before filtering
num_taxa_before <- ntaxa(ps16S_0)


#Filtered ASV based on taxonomy
ps16S_1a <- subset_taxa(ps16S_0, !is.na(Taxon) & !Taxon %in% c("Unassigned"))
ps16S_1b <- subset_taxa(ps16S_1a, !Order %in% c(" o__Chloroplast"))
ps16S_1c<- subset_taxa(ps16S_1b, !Order %in% c("o__Mitochondria"))
ps16S_1d<- subset_taxa(ps16S_1c, !Family %in% c("f__Mitochondria"))
ps16S_1e<- subset_taxa(ps16S_1d, !Family %in% c("f__Chloroplast"))
ps16S_1f<- subset_taxa(ps16S_1e, !Genus %in% c(" g__Mitochondria"))
ps16S_1g<-subset_taxa(ps16S_1f, !Taxon %in% c("d__Eukaryota"))
ps16S_1h<-subset_taxa(ps16S_1g, !Kingdom %in% c("d__Eukaryota"))
ps16S_1i<-subset_taxa(ps16S_1h, !Kingdom %in% c("d__Archaea"))

##decontamination from blank control samples
#libraries
library(here); packageVersion("here")
library(decontam); packageVersion("decontam")
library(phyloseq); packageVersion("phyloseq")
library(Biostrings); packageVersion("Biostrings")
library(tidyverse); packageVersion("tidyverse")

d16S_leaves <- subset_samples(ps16S_1i, tissue %in% c("leaf"))
d16S_leaves <- subset_taxa(d16S_leaves, !taxa_sums(d16S_leaves) == 0)
d16S_soil <- subset_samples(ps16S_1i, tissue %in% c("soil"))
d16S_soil <- subset_taxa(d16S_soil, !taxa_sums(d16S_soil) == 0)
d16S_endophytes <- subset_samples(ps16S_1i, tissue %in% c("endophytes"))
d16S_endophytes <- subset_taxa(d16S_endophytes, !taxa_sums(d16S_endophytes) == 0)
d16S_seeds <- subset_samples(ps16S_1i, tissue %in% c("seed"))
d16S_seeds <- subset_taxa(d16S_seeds, !taxa_sums(d16S_seeds) == 0)

#import clean phyloseq from blanks
phyloseq_bianchi <- readRDS("ps_blanks.rds") 

#prepare dataset
ps16S_1i.2<- merge_phyloseq(d16S_leaves, phyloseq_bianchi)
sample_data(ps16S_1i.2)$FEAST[394:400] <- paste("blank")
base::as.data.frame(phyloseq::sample_data(ps16S_1i.2))
base::as.data.frame(phyloseq::sample_data(d16S_leaves))
view(sample_data(ps16S_1i.2))

#inspect libraries sizes

library("gtable")

df <- as.data.frame(sample_data(ps16S_1i.2))
df$LibrarySize <- sample_sums(ps16S_1i.2)
df <- df[order(df$LibrarySize),]
df$Index <- seq(nrow(df))
df$FEAST<-as.factor(df$FEAST)
ggplot(data=df, aes(x=Index, y=LibrarySize, color=FEAST)) + geom_point()+
  theme(
    legend.title = element_text(size = 5), 
    legend.text = element_text(size = 5),    
    legend.key.size = unit(0.5, "cm"),        
    legend.key.width = unit(0.5, "cm")        
  )

#Prevalence based-decontamination
 sample_data(ps16S_1i.2)$is.neg <- sample_data(ps16S_1i.2)$FEAST == "blank"
contamdf.prev <- isContaminant(ps16S_1i.2, method="prevalence", neg="is.neg")
table(contamdf.prev$contaminant)

# Make phyloseq object of presence-absence in negative controls and true samples
ps.pa <- transform_sample_counts(ps16S_1i.2, function(abund) 1*(abund>0))
ps.pa.neg <- prune_samples(sample_data(ps.pa)$FEAST == "blank", ps.pa)
ps.pa.pos <- prune_samples(sample_data(ps.pa)$FEAST == "leaf", ps.pa)
# Make data.frame of prevalence in positive and negative samples
df.pa <- data.frame(pa.pos=taxa_sums(ps.pa.pos), pa.neg=taxa_sums(ps.pa.neg),
                      contaminant=contamdf.prev$contaminant)
ggplot(data=df.pa, aes(x=pa.neg, y=pa.pos, color=contaminant)) + geom_point() +
  xlab("Prevalence (Negative Controls)") + ylab("Prevalence (True Samples)")

#trim contaminants

ps.noncontam <- prune_taxa(!contamdf.prev$contaminant, ps16S_1i.2)
ps.noncontam

ps16S_1i.5<- merge_phyloseq(ps.noncontam, d16S_endophytes,d16S_seeds, d16S_soil)
view(otu_table(ps16S_1i.5))

ps16S_1i.5= subset_samples(ps16S_1i.5, FEAST != "blank")

#remove samples with less than 100 reads
ps16S_1i.4 <- prune_samples(sample_sums(ps16S_1i.5)>=100, ps16S_1i.5)#Before 
ps16S_1i.5 <- filter_taxa(ps16S_1i.4, function(x) sum(x) > 0, TRUE)
view(sample_data(ps16S_1i.5))
```

##### 10.2- One sink: Table S14 B

```
Packages <- c("Rcpp", "RcppArmadillo", "vegan", "dplyr", "reshape2", "gridExtra", "ggplot2", "ggthemes")
lapply(Packages, library, character.only = TRUE)
library(FEAST)

FEAST_dataset = merge_samples(ps16S_1i.5, "FEAST")
otu_table<- as.matrix(t(otu_table(FEAST_dataset)))
write.csv(otu_table, "otu_table.csv")

#upload the metadata table
metadata <- Load_metadata(metadata_path = "C:/yourPath/metadata.txt")

#upload the otu table
otus <- Load_CountMatrix(CountMatrix_path = "C:/yourPath/otu_table.txt")

#This output is reported in Table S14 B
FEAST_output <- FEAST(C = otus, metadata = metadata, EM_iterations = 1000, different_sources_flag = 1, dir_path = "C:/yourPath/FEAST/", outfile="demo")
```

##### 10.3- Multiple sinks: Table S14 A and Figure S2A

```
Packages <- c("Rcpp", "RcppArmadillo", "vegan", "dplyr", "reshape2", "gridExtra", "ggplot2", "ggthemes")
lapply(Packages, library, character.only = TRUE)
library(FEAST)

FEAST_dataset = merge_samples(ps16S_1i.5, "FEAST2")

sample_data<- as.matrix(sample_data(FEAST_dataset))
otu_table<- as.matrix(otu_table(t(FEAST_dataset)))
write.csv(otu_table, "otu_table_byblocks.csv")
write.csv(sample_data, "sample_byblocks.csv")

metadata <- Load_metadata(metadata_path = "C:/yourPath/metadata.txt")

otus <- Load_CountMatrix(CountMatrix_path = "C:/yourPath/otu_table_byblocks.txt")
#otus <- otus[-c(1, 2), ]

#This output is reported in Table S14 A
FEAST_output <- FEAST(C = otus, metadata = metadata,  EM_iterations = 10000, different_sources_flag = 1, dir_path = "C:/yourPath/", outfile="demo_prova")

  
################################boxplot###############################
#in this case the FEAST function have to be applied for every sample. Since tables are large, there is a first step to help in this process.

#metadata preparation

#extract samples in your metadata table
sample_data<- as.matrix(sample_data(ps16S_1i.5))

# Use samples to create the following vector
colonna_F <- c("01A", "02A", "03A", "04A", "06A", "07A", "08A", "09A", "11A", "13A", 
  "14A", "16A", "17A", "19A", "20A", "21A", "22A", "23A", "24A", "25A", 
  "26A", "27A", "28A", "29A", "30A", "31A", "32A", "33A", "34A", "35A", 
  "36A", "37A", "38A", "39A", "40A", "41A", "42A", "43A", "44A", "45A", 
  "46A", "47A", "48A", "49A", "50A", "51A", "53A", "54A", "55A", "56A", 
  "57A", "58A", "59A", "62A", "63A", "64A", "65A", "66A", "67A", "68A", 
  "69A", "70A", "71A", "72A", "73A", "74A", "75A", "76A", "77A", "78A", 
  "79A", "80A", "81A", "82A", "83A", "84A", "85A", "86A", "87A", "88A", 
  "89A", "90A", "91A", "92A", "93A", "94A", "95A", "96A", "97A", "98A", 
  "99A", "100A", "101A", "102A", "103A", "104A", "105A", "106A", "107A", 
  "108A", "109A", "110A", "111A", "112A", "113A", "114A", "115A", "116A", 
  "117A", "118A", "119A", "120A", "121A", "122A", "123A", "124A", "125A", 
  "126A", "127A", "128A", "129A", "130A", "131A", "132A", "134A", "135A", 
  "137A", "138A", "139A", "495A", "01B", "02B", "03B", "04B", "06B", 
  "07B", "08B", "09B", "11B", "13B", "14B", "16B", "17B", "19B", "20B", 
  "21B", "22B", "23B", "24B", "25B", "26B", "27B", "28B", "29B", "30B", 
  "31B", "32B", "33B", "34B", "35B", "36B", "37B", "38B", "39B", "40B", 
  "41B", "42B", "43B", "44B", "45B", "46B", "47B", "48B", "49B", "50B", 
  "51B", "53B", "54B", "55B", "56B", "57B", "58B", "59B", "62B", "63B", 
  "64B", "65B", "66B", "67B", "68B", "69B", "70B", "71B", "72B", "73B", 
  "74B", "75B", "76B", "77B", "78B", "79B", "80B", "81B", "82B", "83B", 
  "84B", "85B", "86B", "87B", "88B", "89B", "90B", "91B", "92B", "93B", 
  "94B", "95B", "96B", "97B", "98B", "99B", "101B", "102B", "103B", 
  "104B", "105B", "106B", "107B", "108B", "109B", "110B", "111B", "112B", 
  "113B", "115B", "116B", "117B", "118B", "119B", "120B", "121B", "122B", 
  "123B", "124B", "125B", "126B", "127B", "128B", "129B", "130B", "131B", 
  "132B", "134B", "135B", "136B", "137B", "138B", "139B", "495B", "01C", 
  "02C", "03C", "04C", "06C", "07C", "08C", "09C", "11C", "14C", "16C", 
  "19C", "20C", "21C", "22C", "23C", "24C", "25C", "26C", "27C", "28C", 
  "29C", "30C", "31C", "32C", "33C", "34C", "35C", "36C", "37C", "38C", 
  "39C", "40C", "41C", "42C", "43C", "44C", "45C", "46C", "47C", "48C", 
  "49C", "50C", "51C", "53C", "54C", "55C", "56C", "57C", "58C", "59C", 
  "62C", "63C", "64C", "65C", "66C", "67C", "68C", "69C", "70C", "71C", 
  "72C", "73C", "74C", "75C", "76C", "77C", "78C", "79C", "80C", "81C", 
  "82C", "83C", "84C", "85C", "86C", "87C", "88C", "89C", "90C", "91C", 
  "92C", "93C", "94C", "95C", "96C", "97C", "98C", "99C", "100C", "101C", 
  "102C", "103C", "104C", "105C", "106C", "107C", "108C", "109C", "110C", 
  "111C", "112C", "113C", "114C", "115C", "116C", "118C", "119C", "120C", 
  "121C", "122C", "123C", "124C", "125C", "126C", "127C", "128C", "129C", 
  "130C", "131C", "132C", "134C", "135C", "136C", "137C", "138C", "139C", 
  "495C")

# Function to generate the table
generate_table <- function(env_list) {
  sampleID <- c()
  Env <- c()
  SourceSink <- c()
  id <- c()
  
  # ID counter
  id_counter <- 1
  
  for (env in env_list) {
    # Sink insertion
    sampleID <- c(sampleID, paste0("A", id_counter))
    Env <- c(Env, env)
    SourceSink <- c(SourceSink, "Sink")
    id <- c(id, id_counter)
    
    #Source insertion
    for (source in c("soil 1", "soil 2", "soil 3", "seed", "endophytes")) {
      sampleID <- c(sampleID, paste0("A", id_counter + 1))
      Env <- c(Env, source)
      SourceSink <- c(SourceSink, "Source")
      id <- c(id, id_counter)
      
      # To increase the ID row by row
      id_counter <- id_counter
    }
    
    id_counter <- id_counter + 1
  }
  
  df <- data.frame(sampleID, Env, SourceSink, id, stringsAsFactors = FALSE)
  return(df)
}

#Create a data frame with our sample names
df <- generate_table(colonna_F)

# Modify ID to have an increase of 6 columns after every sample
df$id <- rep(seq(1, ceiling(nrow(df) / 6)), each = 6, length.out = nrow(df))

# Save the dataframe as CSV
write.csv(df, "tabella_generata.csv", row.names = FALSE)

###otu_table

library(dplyr)
library(tidyr)

#otu_table for sink samples
otu_table<- as.matrix(otu_table(ps16S_1i.5))
write.csv(otu_table, "otu_table_byblocks.csv") 

#otu_table for source samples
otu_sources = merge_samples(ps16S_1i.5, "FEAST")
otu_table_sources<- as.matrix(t(otu_table(otu_sources)))
write.csv(otu_table_sources, "otu_table_sources.csv")

#now the otu of source samples have to be inserted after the otu of each sink samples. The following script create a table with 5 spaces after each sample, to simplify the process of table creation

# Read the .txt files
otu_table_byblocks <- read.table(file.path(file_path, "otu_table_byblocks.txt"), header = TRUE, sep = "\t")

# Set the first column as row names for both tables
otu_table_byblocks <- otu_table_byblocks %>%
  column_to_rownames(var = names(otu_table_byblocks)[1])

# Function to insert empty columns
insert_empty_columns <- function(df, n) {
  df <- as.data.frame(df)
  # Create an empty list to hold the columns
  expanded_list <- list()
  
  # Iterate through each column in the original data frame
  for (i in 1:ncol(df)) {
    expanded_list[[length(expanded_list) + 1]] <- df[[i]]
    if (i < ncol(df)) { # Add empty columns only if it's not the last column
      expanded_list <- c(expanded_list, replicate(n, NA, simplify = FALSE))
    }
  }
  
  # Combine the list into a new data frame
  expanded_df <- do.call(cbind, expanded_list)
  return(expanded_df)
}

# Create a copy of the blocks table and add empty columns
otu_table_expanded <- insert_empty_columns(otu_table_byblocks, 5)

write.csv(otu_table_expanded, "otu_table_expanded.csv")

#now insert the sources'otus after the otus of each sample

##########################apply function


metadata <- Load_metadata(metadata_path = "C:/yourPath/metadata.txt")

otus <- Load_CountMatrix(CountMatrix_path = "C:/yourPath/otu_table.txt")

FEAST_output <- FEAST(C = otus, metadata = metadata,  EM_iterations = 10000, different_sources_flag = 1, dir_path = "C:/yourPath/", outfile="demo_prova")

##IMPORT OUTPUT
  FEAST_output <- read.table(file="C:/yourPath/demo_prova_source_contributions_matrix.txt",sep="\t",dec = ",",header=TRUE)
  
FEAST_output<-as.data.frame(FEAST_output)
  
write.csv(FEAST_output, "FEAST_output.csv")


#script to delete the NA from the FEAST_output file

shift_left <- function(df) {
  t(apply(df, 1, function(row) {
    non_na_values <- row[!is.na(row)]
    c(non_na_values, rep(NA, length(row) - length(non_na_values)))
  }))
}


df_shifted <- as.data.frame(shift_left(FEAST_output))

write.csv(df_shifted, "df_shifted.csv", row.names = FALSE, na = "")
#join this output with the metadata table

####graphical representation

#import your new metadata table
Origin_dataset <- read.table(file="C:/yourPath/sample_byblocks.txt" ,sep="\t",dec = ",",header=TRUE)

Origin_dataset$A2_soil.1<-as.numeric(Origin_dataset$A2_soil.1)
Origin_dataset$A3_soil.2<-as.numeric(Origin_dataset$A3_soil.2)
Origin_dataset$A4_soil.3<-as.numeric(Origin_dataset$A4_soil.3)
Origin_dataset$A5_seed<-as.numeric(Origin_dataset$A5_seed)
Origin_dataset$A6_endophytes<-as.numeric(Origin_dataset$A6_endophytes)
Origin_dataset$Unknown<-as.numeric(Origin_dataset$Unknown)
Origin_dataset$genotype2<-as.factor(Origin_dataset$genotype2)
Origin_dataset$variety<-as.factor(Origin_dataset$variety)
Origin_dataset$block<-as.factor(Origin_dataset$block)
Origin_dataset$genetic.distance<-as.factor(Origin_dataset$genetic.distance)
Origin_dataset$sampleID3<-as.factor(Origin_dataset$sampleID3)

#remove Na in "genetic distance" column
Origin_dataset <- Origin_dataset %>%
  filter(!is.na(genetic.distance) & genetic.distance != "" & genetic.distance != "Na") %>%
  droplevels()

# Reshape data
library(reshape2)

dati_long <- melt(Origin_dataset, id.vars = c("sampleID3", "variety", "genetic.distance", "block"), measure.vars = c("A2_soil.1", "A3_soil.2", "A4_soil.3", "A5_seed", "A6_endophytes"))

dati_long$value<-as.numeric(dati_long$value)
dati_long$variable<-as.factor(dati_long$variable)
dati_long$value <- round(dati_long$value, 2)
dati_long$log_value <- log(dati_long$value)

#Figure S2A
#by block
# Imposta l'ordine desiderato per 'variable'
ordine_variabili <- c("A5_seed", "A4_soil.3", "A2_soil.1", "A3_soil.2", "A6_endophytes")
dati_long$variable <- factor(dati_long$variable, levels = ordine_variabili)

block_colors <- c(
  "A" = "#fcbba1",
  "B" = "#fa9fb5",
  "C" = "#c51b8a"
)

#boxplot
gg1 <- ggplot(dati_long, aes(x = variable, y = value, color=block)) +
  geom_boxplot() +
  geom_jitter(aes(color = block), size = 1,  position = position_jitterdodge(jitter.width = 0.3, dodge.width = 0.8)) +
  labs(x = "Sources", y = "Average source contribution") +
  facet_wrap(~ variable, scales = "free_x", ncol=5) +
  scale_color_manual(values = block_colors) + 
  theme_classic() +
  ggtitle("Average source contribution") +
  theme(axis.text.x = element_text(angle = 90, vjust = 0.5, hjust = 1),
        strip.text = element_text(size = 12),
        legend.position = "bottom",
        panel.spacing = unit(1, "lines"))
```
